# Iterative improvement of deep learning models using synthetic regulatory genomics

**DOI:** 10.1101/2025.02.04.636130

**Authors:** André M. Ribeiro-dos-Santos, Matthew T. Maurano

## Abstract

Deep learning models can accurately reconstruct genome-wide epigenetic tracks from the reference genome sequence alone. But it is unclear what predictive power they have on sequence diverging from the reference, such as disease- and trait-associated variants or engineered sequences. Recent work has applied synthetic regulatory genomics to characterized dozens of deletions, inversions, and rearrangements of DNase I hypersensitive sites (DHSs). Here, we use the state-of-the-art model Enformer to predict DNA accessibility and RNA transcription across these engineered sequences when delivered at their endogenous loci. At high level, we observe a good correlation between accessibility predicted by Enformer and experimental data. But model performance was best for sequences that more resembled the reference, such as single deletions or combinations of multiple DHSs. Predictive power was poorer for rearrangements affecting DHS order or orientation. We use these data to fine-tune Enformer, yielding significant reduction in prediction error. We show that this fine-tuning retains strong predictive performance for other tracks. Our results show that current deep learning models perform poorly when presented with novel sequence diverging in certain critical features from their training set. Thus an iterative approach incorporating profiling of synthetic constructs can improve model generalizability and ultimately enable functional classification of regulatory variants identified by population studies.

## Introduction

Most genetic associations with human diseases and traits lie within noncoding regulatory DNA (Maurano et al. 2012). Genome-scale methods to analyze the function of noncoding regulatory elements within their relevant context would improve our ability for direct assessment of disease-associated variation. Profiling of allelic transcription factor (TF) activity across multiple cell and tissue types has shown that both genomic and tissue context modulates the penetrance of noncoding variants (Halow et al. 2021), but direct biochemical assessment of each variant in every relevant cell type and state remains impractical.

An alternative approach is to train machine learning models on reference genomic sequence and then apply them to genetic variation data. Recent machine learning models including gkmSVM (Lee et al. 2015), DeepSEA (Zhou and Troyanskaya 2015), Enformer (Avsec et al. 2021a), and BPNet (Avsec et al. 2021b) have demonstrated impressive performance in predicting functional genomics signals such as expression and DNA accessibility from sequence alone. Those models were trained on large functional genomics datasets (e.g. DNase-seq or CAGE) across a wide variety of human and mouse cell and tissue types generated by large projects such as ENCODE (The ENCODE Project Consortium et al. 2020; Meuleman et al. 2020), the Roadmap Epigenomics Mapping Project (Roadmap Epigenomics Consortium et al. 2015), and FANTOM (The FANTOM Consortium and the RIKEN PMI and CLST (DGT) et al. 2014). Analysis of current models on inter-individual genetic variation data has yielded little improvement relative to more parsimonious approaches (Maurano et al. 2015; Dey et al. 2020; Karollus et al. 2023; Toneyan et al. 2022; Sasse et al. 2023; Huang et al. 2023). The opacity of the more complex models reduces their contribution to mechanistic understanding, and could mask overfitting resulting in degraded performance under new circumstances. Furthermore, training sequence is universally drawn from the human or mouse reference genomes, whose genomic features can be highly correlated, particularly for long-range interactions (Xi and Beer 2018; Whalen and Pollard 2019). Such an approach limits the explored sequence space due to constraints imposed by evolution and could reduce model performance on mutations and engineered sequences diverging significantly from reference. In principle, genomic sequences from different species (Kelley 2020; Cochran et al. 2022) or random sequence (de Boer and Taipale 2024) could expand the sequence space for training. Thus, the problem remains how to generate sufficient functional data reflects function at the endogenous genomic locus.

Sources of training data beyond the reference sequence are limited by the scale of sequence amenable to manipulation, and its relevance to the regulation architecture of the endogenous locus. Massively parallel reporter assays (MPRAs) assay transiently transfected plasmids or randomly integrated sequences with little genomic context and thus cannot accurately model the effect of surrounding regulatory sites at the endogenous locus (Kircher et al. 2019; Kosicki et al. 2025). Large-scale CRISPR-Cas-based genome engineering strategies including saturation genomic editing overcome the inability of MPRAs to reproduce accurate genomic context (Findlay et al. 2014; Martyn et al. 2025). However, each CRISPR guide provides control over a limited genomic range (100-1000 bp). Thus these approaches are more suited to the deletion of existing sequence than its rearrangement or the addition of novel sequence and are difficult to use for multi-step engineering of diploid cells. Training on allelic variation in expression (Sasse et al. 2023), chromatin accessibility (Maurano et al. 2015; Halow et al. 2021), or genome-wide association study variants (Mostafavi et al. 2023) are limited by the standing genetic variation and the effects of selection. A synthetic regulatory genomic approach is uniquely able to uncover novel biology, and these new technologies will move the field towards consideration of regulatory function at a locus scale.

Recent work applying synthetic regulatory genomics has highlighted the influence of genomic context on enhancer function (Ribeiro-Dos-Santos et al. 2022; Brosh et al. 2023), and the discrepancy between reporter assays and function at the endogenous locus (Brosh et al. 2023). The Big-IN platform enables targeted integration of large BACs upwards of 160 kb into human and murine embryonic stem cells, overcoming previous limitations of scalability and portability across genomic loci and cellular contexts (Brosh et al. 2021). Integration of a modular landing pad (LP) enables subsequent recombinase-mediated cassette exchange (RMCE) of large payloads and efficient positive/negative selection for correct clones. Hundreds of different payloads have been delivered to these cell lines at multiple loci (Brosh et al. 2021; Mitchell et al. 2021; Pinglay et al. 2022; Brosh et al. 2023; Ordoñez et al. 2024; Zhang et al. 2023). Compared to editing methods such as saturation genome editing, synthetic regulatory genomics is not limited to derivatization of the reference sequence, but can deliver a nearly unlimited set of sequences.

Here, we explore the application of deep learning models to predict regulatory features at loci engineered through synthetic regulatory genomics. We evaluate the ability of these models to predict the functional behavior of regulatory elements when subject to complex deletions and rearrangements. Further, we evaluate the model prediction of functional genomics for novel synthetic sequences and long-range regulatory elements interaction. Finally, we demonstrate the potential design for iterative improvement of such models by integrating synthetic regulatory dataset. Our work suggests a key role for synthetic regulatory genomics in training future genomic deep learning models.

## RESULTS

### Enformer performance predicting synthetic payloads

To probe the varied performance of deep learning models across different regimes, we analyzed the Enformer model due to its high accuracy and wide receptive field that allows investigation of large regulatory landscapes (Avsec et al. 2021a). Based on previous work showing that Enformer retains most of its predictive performance with shorter input windows (Karollus et al. 2023), we adapted the published Enformer model to use flexible input and output lengths to enable capturing relevant regulatory elements while maintaining high computational efficiency, as well as outputting predictions for large synthetic payloads (**Methods**). We confirmed the adapted model produces nearly identical results to the published version, including at different input window sizes (**Supplemental Fig. S1, Supplemental Fig. S2**).

We then evaluated the predictive power of Enformer on synthetic sequences related to the reference genome but differing in significant aspects. We started with a published analysis which dissected the *Sox2* locus in mouse embryonic stem cells (mESC) (Brosh et al. 2023). 80 synthetic payloads were delivered replacing either the full 143-kb locus or 43-kb locus control region (LCR) required for *Sox2* expression in mESC, and included multiple categories of variation, such as deletions, duplications, rearrangements, inversions, and surgical TF site deletions (**Fig. 1A**). As the published Enformer model did not include CAGE predictions in mESC (Avsec et al. 2021a), we trained Enformer on publicly available CAGE data produced at RIKEN (**Methods**, **Supplemental Fig. S3**). We then predicted transcription at the *Sox2* transcription start site (TSS) for the synthetic payloads (**Supplemental Fig. S4A**). However, comparison with measured expression showed no correlation, and in particular, Enformer did not predict the effect of deleting the LCR, which should result in total ablation of *Sox2* expression in mESC (Zhou et al. 2014; Li et al. 2014). We focused on the 70 payloads delivered replacing the LCR. We predicted DNA accessibility at the LCR for each payload using Enformer, and summed accessibility at LCR DNase I hypersensitive sites (DHSs) 19-28 as a proxy for the overall LCR activity (**Supplemental Table S1, Supplemental Table S2**). Predicted LCR accessibility showed a good correlation with experimentally measured *Sox2* expression, confirming that accessibility can serve as a good proxy for expression in this context (**Fig. 1B**).

**Fig. 1.**
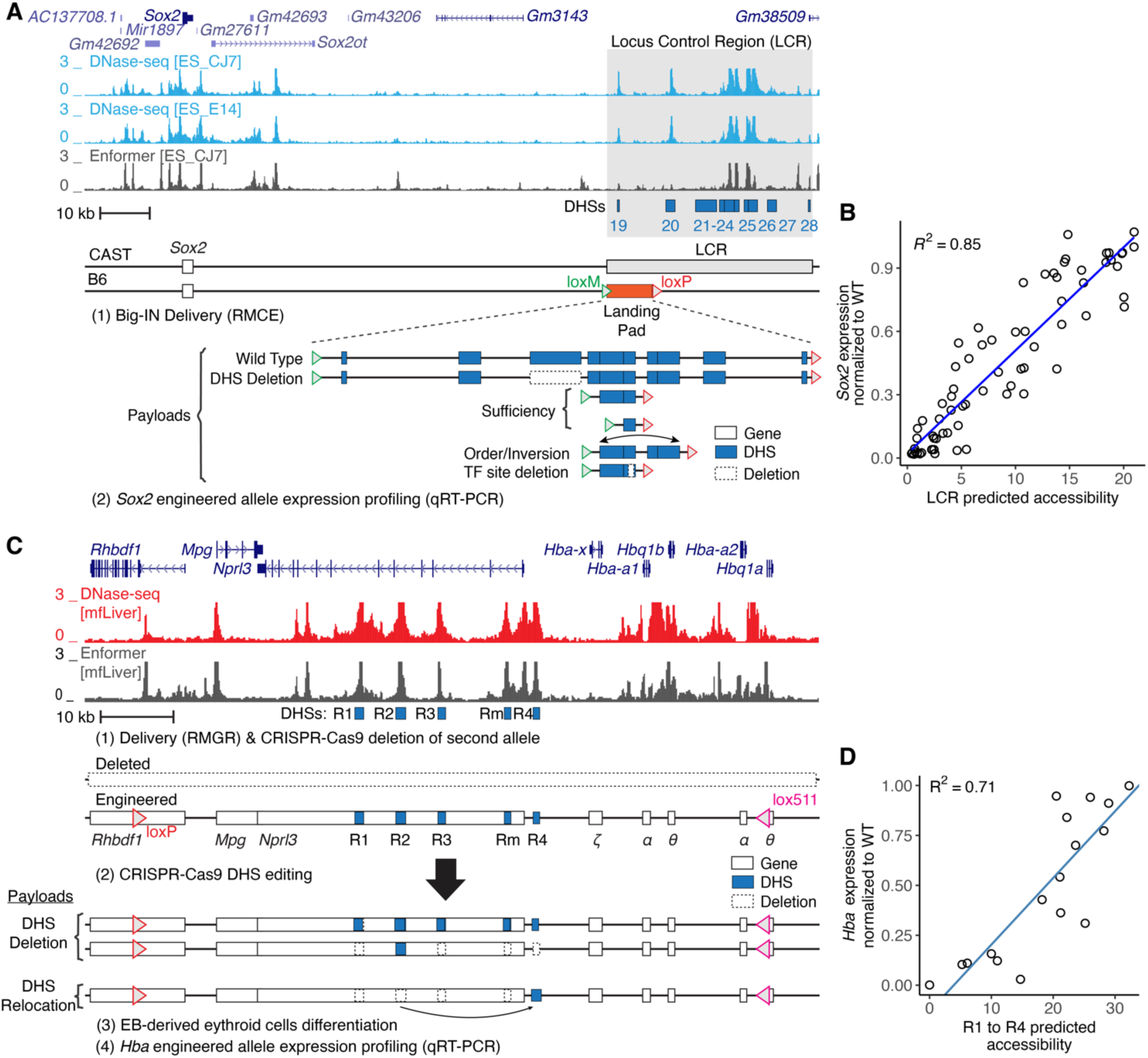
Assessment of Enformer performance on synthetic sequences. (A) Schematic of *Sox2* locus showing DNase-seq in mouse embryonic stem cells (mESC) and Enformer prediction Engineered CAST/B6 mESC cells harbor a Big-IN landing pad replacing one allele of the *Sox2* LCR for delivery of multiple synthetic payloads (Brosh et al. 2023). Locus control region (LCR) is highlighted in gray. DHSs are indicated by blue boxes. DHS deletions are indicated by dashed boxes. (B) *Sox2* expression was characterized by allele-specific qRT-PCR. Each point represents experimentally measured *Sox2* expression and Enformer prediction of a synthetic payload (N=70). Engineered *Sox2* allele expression was scaled between 0 (ΔSox2) and 1 (WT). LCR activity was measured as the summed accessibility over all LCR (DHSs 19-28) in mESC_CJ7. (C) Schematic of *α-globin* locus showing DNase-seq of mouse fetal liver (mfLiver), Enformer predictions, and *α-globin* locus engineering strategy by CRISPR-Cas9 editing and characterization using qRT-PCR (Blayney et al. 2023). Enhancers R1-R4 are indicated by blue boxes. Deleted regions are represented by dashed boxes. (D) *Hba* expression of the resulting cells was profiled by qRT-PCR, normalized to *Hbb* and scaled as a proportion of WT expression. Each point represents experimentally measured *Hba* expression and Enformer prediction of an enhancer configuration (N=17). *Hba* expression was measured by qRT-PCR, normalized to *Hbb* and scaled as a proportion of WT expression. Predicted enhancers (R1-R4) activity was measured as the sum of maximum predicted accessibility at enhancers R1 to R4. Blue lines indicate linear regressions (B and D).

To evaluate our accessibility prediction approach on a different locus and cell type, we analyzed a series of enhancer deletions and relocations at the α-globin locus (**Supplemental Table S1**) characterized in mouse embryonic body-derived erythroid cells (**Fig. 1C**) (Blayney et al. 2023). Starting in mESCs, one α-globin allele was deleted using CRISPR-Cas9, while the other was engineered to support subsequent delivery of engineered *α-globin* payloads ranging up to 85 kb through recombinase-mediated genomic replacement (RMGR). Enhancer deletions and relocations were then engineered with CRISPR-Cas9. We verified that Enformer accessibility predictions matched the previously published experimentally measured ATAC-seq tracks (**Supplemental Fig. S5**). Similarly to the predictions on the *Sox2* dataset, the sum of predicted accessibility of all five enhancers (**Supplemental Table S2**) presented a good correlation to experimentally measured *Hba* expression (**Fig. 1D**).

We then investigated predictive performance for individual TF recognition sequences identified within *Sox2* LCR DHSs 23-24, (Brosh et al. 2023). We used Enformer to perform an in silico scanning deletion analysis of those DHSs replacing 16 bp with Ns (**Fig. 2A**). The accessibility predictions correctly identified two regions identified as essential in the experimental analysis. However, Enformer completely missed a key cluster of TF recognition sequences (TF sites 23.5– 23.8) (**Fig. 2A**), which experimental deletion analysis showed to be essential (Brosh et al. 2023). Furthermore, published analysis has shown that DHSs 23 and 24 have positive synergy, in that the activity of one DHS is greatly increased by the presence of another DHS in superadditive fashion (Brosh et al. 2023). Enformer predicted no synergy between DHSs 23 and 24 as deletions in one DHS were predicted to have no effect on accessibility of the neighboring DHS (**Fig. 2A**). We compared Enformer predictions to specific payloads harboring single and multiple TF site deletions or mutations within DHSs 23 and 24. While Enformer predicted the activity payloads with fully disabling deletions, its worst prediction was for the DHS23–24 pair, and it overestimated the activity of medium-effect TF site deletions (**Fig. 2B**), suggesting it had difficulty quantitatively predicting the relationship between multiple TF sites. Finally, the CAGE model failed to predict an effect for any deletions (**Supplemental Fig. S4B**). Thus, while deep learning models can recapitulate gross features of synthetic sequences derived from genomic reference sequence, their performance has gaps, most notably the synergy among neighboring regulatory elements.

**Fig. 2.**
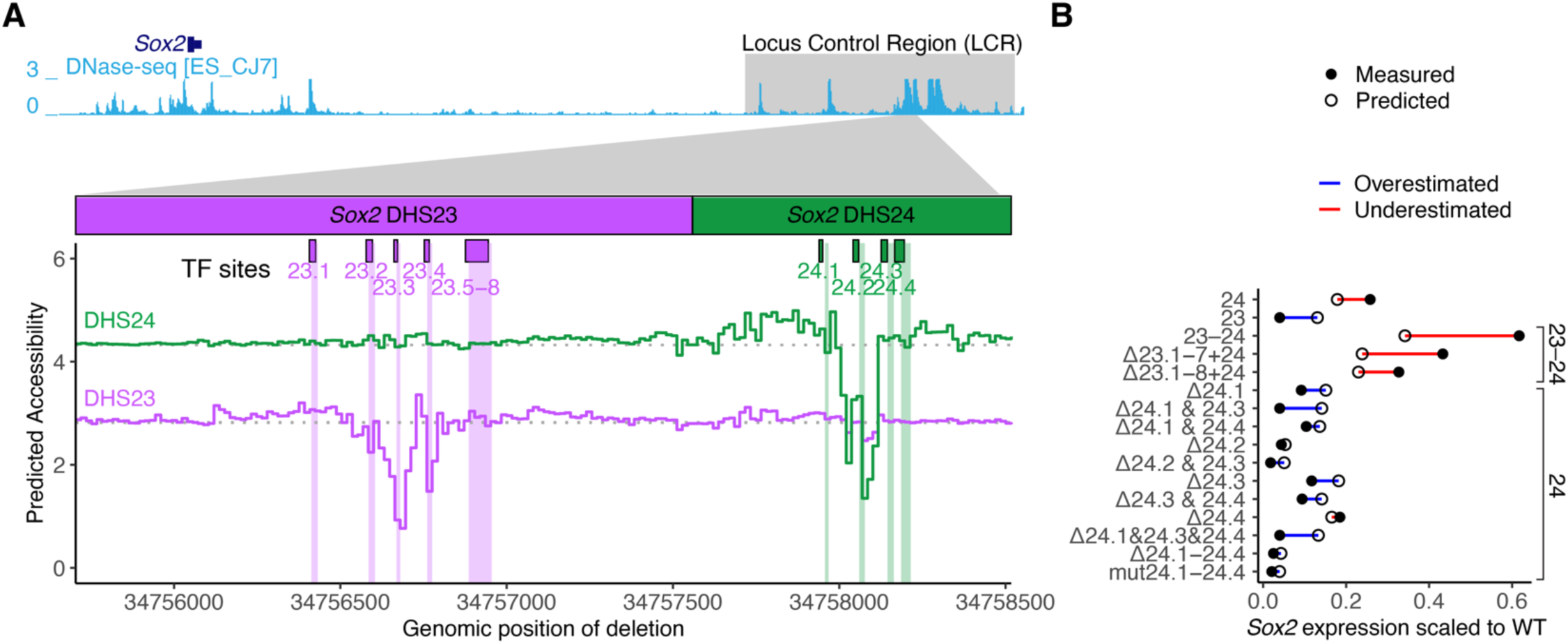
Enformer poorly models individual TF recognition site deletions. (A) In silico scanning deletion analysis of *Sox2* DHS23 and DHS24 accessibility. Enformer was used to predict accessibility (mESC_CJ7 DNase-seq) for a series of DHS23-24 virtual payloads replacing the *Sox2* LCR. 16-bp deletions were encoded by replacing the payload sequence with Ns sliding across the length of DHS23 and DHS24. Colored lines indicate the DHS accessibility for each deletion location (x-axis). Shown are DHS23 and DHS24 accessibility according to deletion position are represented by the purple and green lines respectively. Horizontal dotted lines indicate baseline accessibility. Boxes above indicate relevant TF recognition sequences. (B) Comparison of experimentally measured and predicted accessibility at payloads including DHSs 23 and/or 24 with TF recognition sequences deleted (Δ) or mutated (mut), delivered in place of the *Sox2* LCR, and profiled for expression (Brosh et al. 2023). Difference between measured (closed circle) and predicted (open circle) accessibility is shown by a line and colored by direction of difference. Predicted expression was scaled to WT using a linear regression fitted to all payload examples in Fig. 1B.

Finally, to explore performance of deep learning models on sequences even further diverged from the reference genome, we investigated a synthetic “backwards” sequence in which the *HPRT1* gene sequence was reversed but not complemented (Camellato et al. 2024). This synthetic sequence shares many features with the native *HPRT1* (e.g., base composition, homopolymer runs, and other short repeats) but disrupts higher-order features such as coding sequence and TF recognition sites (except palindromes). mESCs were engineered to replace the endogenous mouse *Hprt* locus with three synthetic payloads ranging from 95-101 kb (**Fig. 3A**): (i) *HPRT1* containing the native human sequence; (ii) SynHPRT1R containing its reversed but not complemented sequence; and (iii) SynHPRT1R^noCpG^ wherein all CpG sites were removed (**Supplemental Table S1**). For the *HPRT1* payload in native hESC, Enformer correctly predicted activity at two experimentally observed DHSs (**Fig. 3B**). We then compared Enformer predictions to experimentally measured DNA accessibility (ATAC-seq) in mESC for all three payloads (**Fig. 3C**). The *HPRT1* payload only showed accessibility by ATAC-seq at the gene promoter, and both the reversed payloads (SynHPRT1R and SynHPRT1R^noCpG^) showed no accessibility at all. In comparison, Enformer predicted false positive peaks in both *HPRT1* and SynHPRT1R^noCpG^ payloads, and it predicted above background accessibility throughout SynHPRT1R (**Fig. 3C**). The false-positive peaks in *HPRT1* corresponded to peaks observed in the experimental data for hESC but not mESC, suggesting that the same sequence behaves differently in the two cellular contexts, and that the Enformer predictions do not accurately reflect the mESC trans-regulatory environment. These SynHPRT1R false positives might be related to sequence CpG content, since they were absent in SynHPRT1R^noCpG^. These results reinforce model limitations when assessing novel synthetic sequences.

**Fig. 3.**
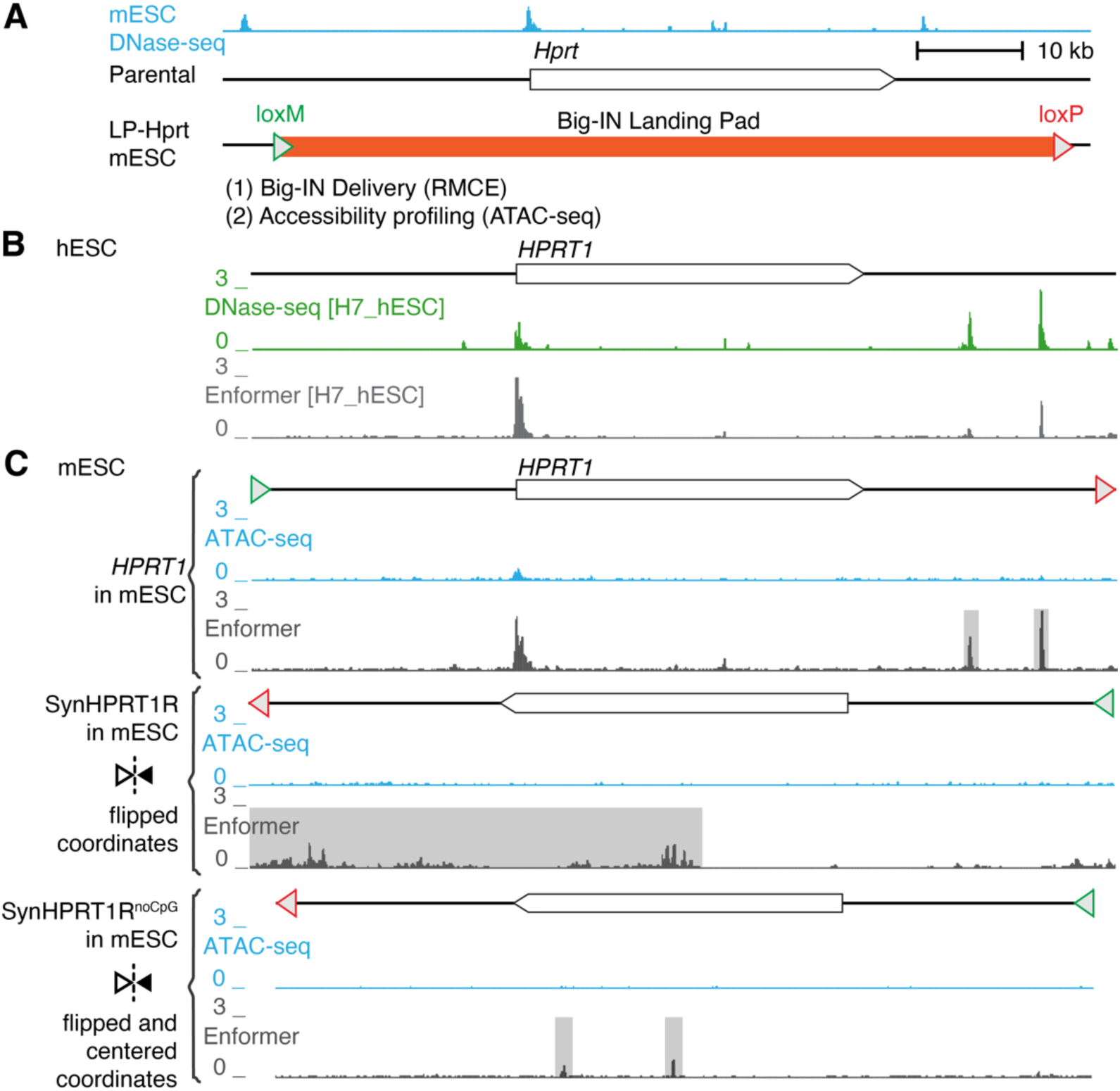
Limited predictive performance on novel synthetic sequences. (A) Schematic of mouse *Hprt* locus and synthetic payload profiling strategy (Camellato et al. 2024). mESC DNase-seq is shown at top. mESCs were engineered to replace their *Hprt* locus with Big-IN landing pad, which was used to deliver three synthetic payloads: human *HPRT1* locus, SynHPRT1R (reversed but not complemented human *HPRT1* locus) and SynHPRT1R^noCpG^ (SynHPRT1R with all CpG removed) payloads. Engineered cells were characterized by ATAC-seq. (B) Schematic of human *HPRT1* locus. Shown is DNase-seq of human H7_hESC and corresponding Enformer prediction. (C) ATAC-seq and Enformer accessibility predictions for engineered mESC. Shown synthetic payloads (*HPRT1*, SynHPRT1R, and SynHPRT1R^noCpG^) profiled and predicted accessibility. Schematics of each synthetic payload are shown above accessibility tracks. Reversed payloads (SynHPRT1R, and SynHPRT1R^noCpG^) are flipped horizontally and SynHPRT1R^noCpG^ was centered to match *HPRT1* coordinates. Enformer false positive predictions are highlighted in gray.

### Enformer does not accurately model distal promoter-enhancer regulation

Previous reports have shown that activation of *Sox2* by its LCR decreases with enhancer-gene distance (Zuin et al. 2022). We tested whether Enformer was able to replicate the LCR-*Sox2* distance dependence curve by simulating placement of the *Sox2* LCR at 10-kb intervals both downstream and upstream of the *Sox2* promoter. For each placement, we predicted mESC CAGE at the *Sox2* promoter and DNase-seq signal at the promoter and LCR (**Fig. 4A**). We found that predicted accessibility remained mostly unchanged and was independent of enhancer-gene distance (**Fig. 4B**, **Supplemental Fig. S6**). To evaluate the impact of genomic distances in a different sequence context, we flanked the *Sox2* proximal region with two copies of SynHPRT1R^noCpG^ (upstream and downstream) and again evaluated the impact of inserting the whole *Sox2* LCR at 10-kb intervals (**Fig. 4C**). This experiment yielded similar results, in that Enformer did not predict a significant distance effect (**Fig. 4D**). In contrast to predictions at the endogenous locus, the presence of the LCR at any position in SynHPRT1R^noCpG^ was predicted to increase *Sox2* promoter accessibility above the ΔLCR background. Thus, Enformer incorrectly predicts no effect of distance between the *Sox2* promoter and LCR, inconsistent with the experimental data. This is consistent with other reports that existing transformer models do not effectively model long-range interactions despite their wide receptive fields (Karollus et al. 2023).

**Fig. 4.**
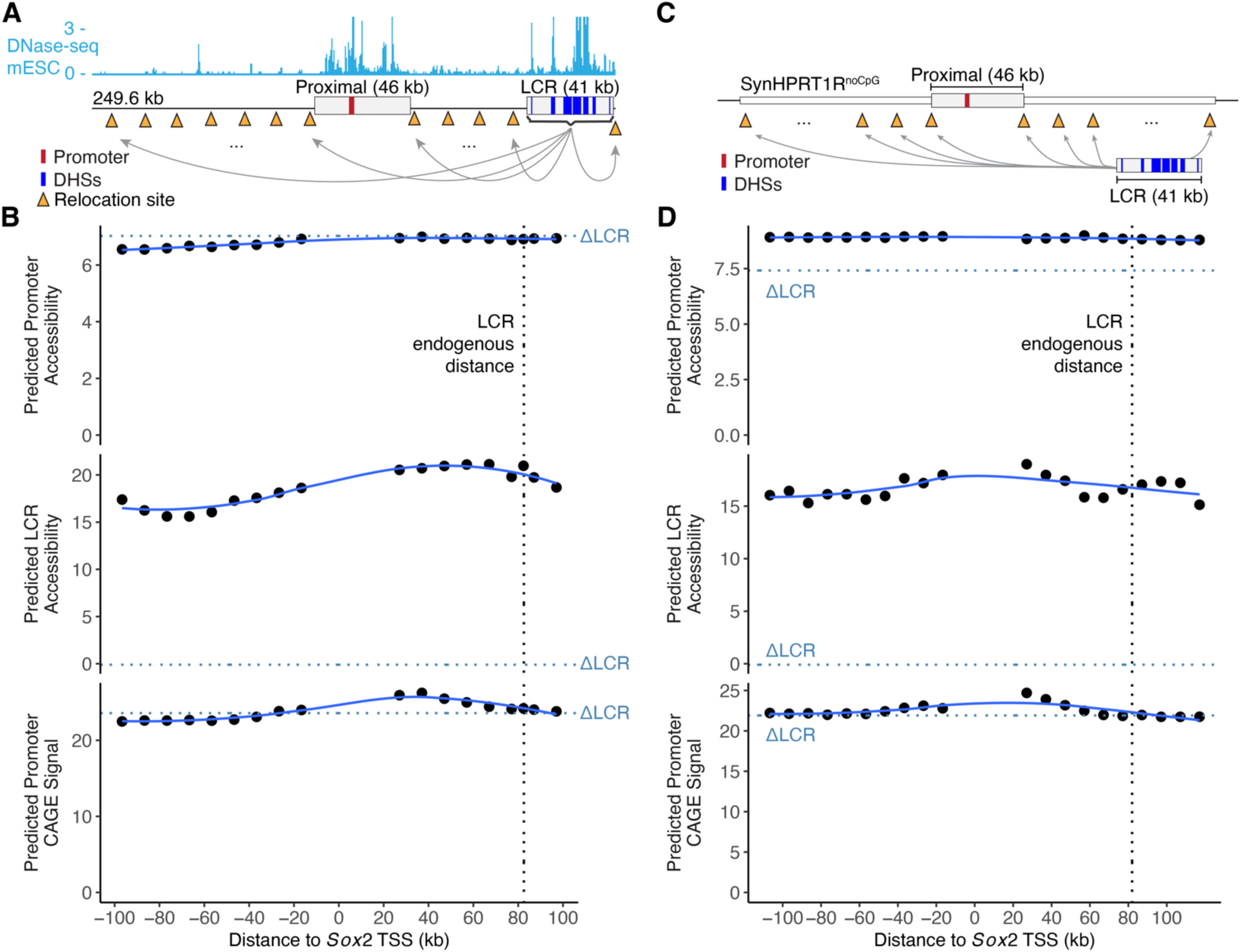
Enformer does not predict effect of LCR relocation on *Sox2* accessibility or expression. (A,C) Schematics of relocation simulation of the *Sox2* LCR in 10-kb intervals surrounding the promoter in its endogenous locus (A) or neutral synthetic sequence (SynHPRT1R^noCpG^) (C); red rectangles indicate *Sox2* promoter and blue rectangles LCR DHSs 19-28. (B, D) Enformer predictions for a 249.6-kb window encompassing the whole virtual locus and any remaining space was filled with Ns. Each point represents predicted accessibility of *Sox2* promoter (top) or LCR (middle), or promoter CAGE signal (bottom) for a given distance between *Sox2* TSS and LCR. Promoter CAGE signal and accessibility was measured as the site maximum predicted signal, and LCR accessibility as the sum of DHSs 19-28 maximum predicted signal. Vertical black dotted line indicates endogenous LCR distance to TSS. Horizontal dotted blue lines indicate prediction when LCR was replaced by SynHPRT1R^noCpG^ fragment of the same size. Solid blue lines show LOESS fits.

### High-level Enformer predictions depend on background sequence properties

This lack of distance dependence contrasts with a prior report that Enformer predicts a key role for enhancer-promoter distance (Toneyan and Koo 2024). To explore this apparent inconsistency, we reproduced the effect of the position of their 74 candidate enhancers on CAGE signal at their target promoter in K562 cells. We built a virtual sequence for each enhancer-promoter pair by extracting a 196,608-bp window centered at the targeted gene transcription start site (TSS) from the human reference genome. We randomly shuffled dinucleotides within the targeted region sequence and restored the TSS block (defined as the central 5-kb region). We then generated a series of virtual sequences by inserting the candidate enhancer in 10-kb intervals from the TSS. The TSS CAGE signal was measured for each TSS-enhancer position by averaging the signal of ten random dinucleotide shuffles. Consistent with the published analysis, all promoter-enhancer pairs showed a steep reduction of CAGE signal with increasing TSS-enhancer distance (**Supplemental Fig. S7A**).

We reasoned that the contrast with the *Sox2* distance analysis might be related to the choice of background sequences. Thus, we repeated the experiment using 10 randomly generated background sequences based on the dinucleotide content of SynHPRT1R^noCpG^. In this context, we observed a selective distance response on TSS CAGE signal, with many DHS relocations producing little to no impact on TSS CAGE signal (**Supplemental Fig. S7B**). We confirmed all promoter/enhancer pairs were still expressed in this background (**Supplemental Fig. S7C**). Thus, we observed completely different responses depending on the sequence context. To more specifically attribute this context dependence to sequence features, we investigated the nucleotide composition at the endogenous loci used. We found that sites with G+C content < 47% showed the same distance dependence regardless of context, while the sequences that differed when tested in SynHPRT1R^noCpG^ all had G+C ≥47% (**Supplemental Fig. S7D**). Notably, the human and mouse reference genomes, and SynHPRT1R^noCpG^ all had lower G+C (41%, 42%, and 38%, respectively) suggesting the in silico distance dependence is disrupted by a mismatch in nucleotide composition. This highlights the importance of background generation strategy and the impact of key genome sequence parameters such as G+C content on evaluation of model predictions.

### Deep learning models struggle to predict cell-type selective DHSs

Current deep learning models are trained to predict multiple regulatory tracks (i.e. cell types) at the same time (Kelley et al. 2016, 2018; Avsec et al. 2021b). We reasoned that this multi-task strategy could favor detection of pervasive regulatory sites present across multiple cell types to the detriment of more cell-selective features (Schreiber et al. 2020; Kathail et al. 2024). We compared peaks called from Enformer predicted accessibility against matching experimental DNase-seq peaks. Peaks were partitioned according to their activity across 141 cell and tissue types from reference DNase-seq datasets which included including 105 human tracks and 36 mouse tracks (**Supplemental Table S3**). This analysis showed that tissue-specific peaks were highly enriched for false positives, while true positives tended to be constitutively active sites (**Fig. 5A-B**). Enformer presented increasingly higher positive predictive value (PPV) with decreasing tissue-specificity (**Fig. 5C**). Similar trends were observed when investigating mESC promoter, CTCF and non-CTCF sites (**Supplemental Fig. S8**), suggesting the lack of sensitivity to cell-type selective sites is not limited to particular classes of genomic regulatory elements. The relatively low proportion of false positive sites among the more cell-type specific sites suggests that some of them can be explained by features bleeding through from other training tracks.

**Fig. 5.**
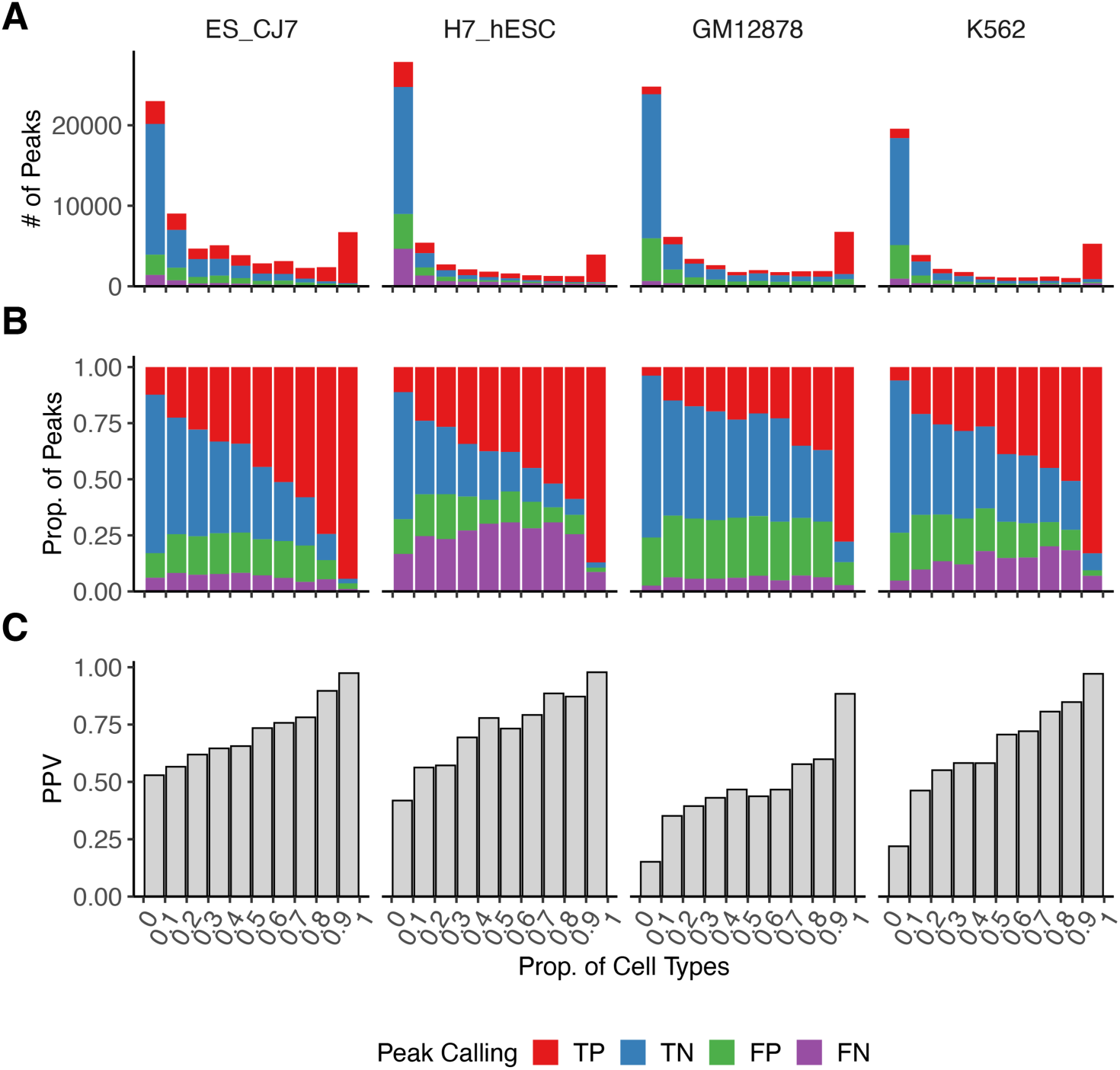
Enformer favors detection of shared regulatory sites. (A-C) Peak calling comparison between mESC_CJ7, H7_hESC, GM12878, and K562 DNase-seq and their respective Enformer predictions (detailed in **Methods**). Peaks were binned according to the proportion of cell types represented in a collection of mouse (N=36) and human (N=105) reference DNase-seq datasets (**Supplemental Table S3**). Peaks are classified as true positives (TP), when found in both reference DNase-seq and Enformer predictions; true negative (TN), when not found in either; false positive (FP), when found only in Enformer predictions; and false negative (FN), when found only in reference DNase-seq. A 25.6-kb input window was used for predictions. (A) Total number of peaks colored by category. (B) Relative proportion of categories per bin. (C) Positive predictive value (PPV) per bin.

### Fine-tuning on synthetic regulatory genomics datasets improves predictive performance

We then developed a fine-tuning strategy to improve model performance through incorporation of synthetic regulatory genomics datasets. We added a new independent output layer that uses the baseline Enformer feature extraction trunk to predict *Sox2* expression (**Fig. 6A**). The new output layer is composed of a self-attention layer to capture relevant features independently of position and a dense layer to combine the resulting signal into a single expression prediction value. This allowed us to fine-tune Enformer internal feature extraction without affecting mouse or human regulatory track heads. We evaluated three configurations of the new output self-attention layer: SingleHead 64/64, SingleHead 64/128, and MultiHead 64/64 (detailed in **Methods**). They all presented similar performance and any can be considered an adequate architecture.

**Fig. 6.**
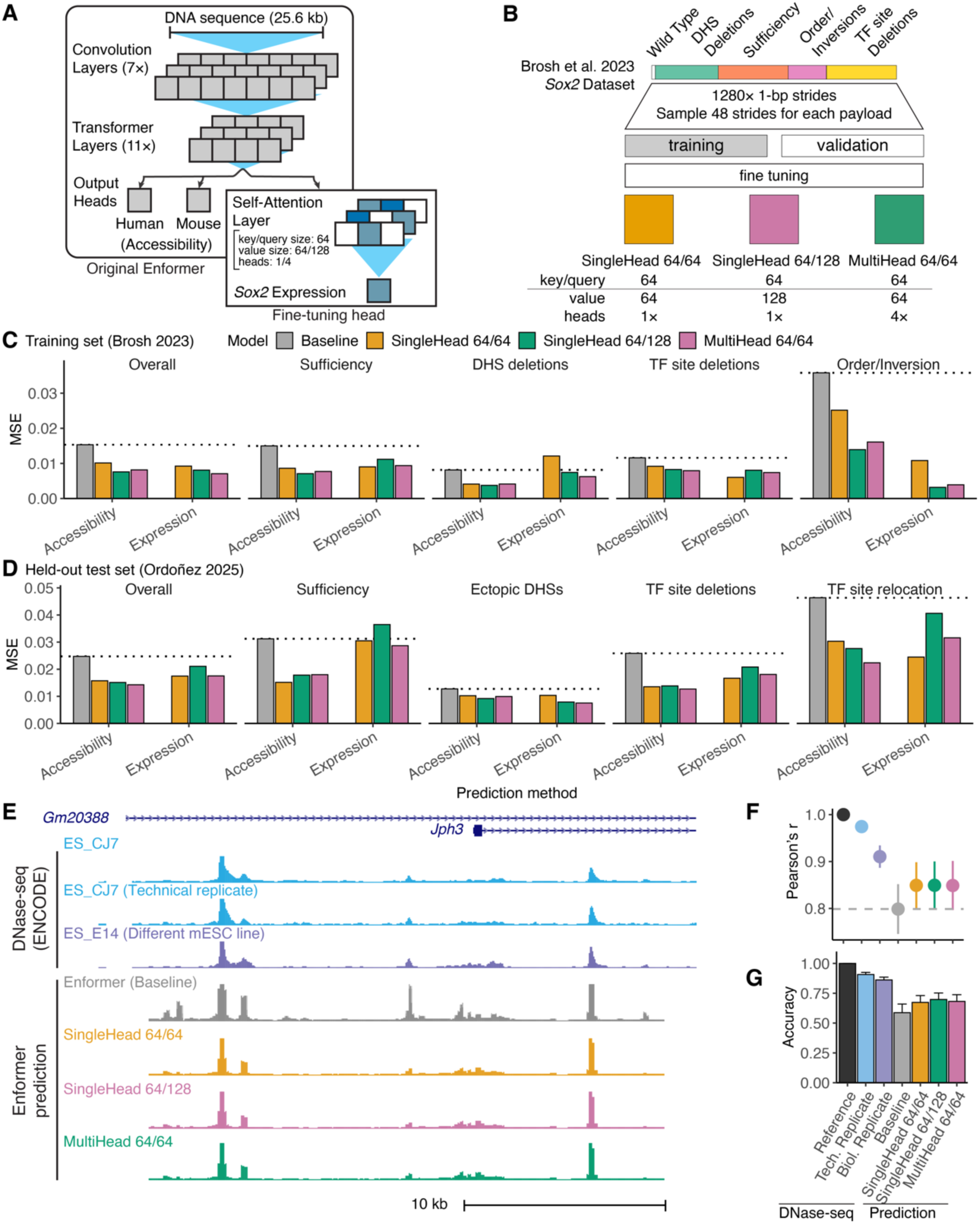
Improved model prediction by incorporating synthetic regulatory genomics data. (A) Enformer fine-tuning architecture. (B) Dataset augmentation and training strategy. (C-D) Predictive performance of baseline and fine-tuned Enformer models on training (Brosh et al. 2023) (C) and held-out test (Ordonez et al. 2025) (D) sets. Expression was estimated either by summing the maximum predicted accessibility of all LCR DHS (Accessibility) or directly from the new output head (Expression). MSE, mean squared error. Results presented for overall dataset and payload categories containing more than two payloads. Horizontal dotted lines show baseline performance. (E) Experimental and predicted mESC DNase-seq at the *Jph3* locus. Shown is DNase-seq of the prediction target (mESC_CJ7), a technical replicate, an independent cell line (mESC_E14), and accessibility predictions made with baseline and fine-tuned Enformer models. (F-G) Genome-wide predictive performance, showing experimental mESC replicates to indicate maximum performance. Error bars indicate ±1 s.d.. (F) Mean correlation (*r*) with the prediction target mESC_CJ7 at peaks. Peak signal was measured as the maximum at bins overlapping each hotspot. (G) Mean peak calling accuracy from predicted signal.

For training, we sampled from different windows centered on the experimentally assessed payloads inserted to replace the endogenous *Sox2* LCR in the mouse reference genome (**Fig. 6B, Supplemental Fig. S9, Methods**). To assess whether our results were sensitive to overfitting from training and evaluating our model on the same collection of payloads, we developed a more conservative training strategy that randomly partitioned payloads into independent training and validation datasets (**Supplemental Fig. S9C**). Training ten batches of models using this partitioned strategy produced similar results to the full dataset training, but with higher variance due to the reduced numbers of payloads per category (**Supplemental Fig. S10**). Thus, we employed the model trained with the full dataset for all subsequent analysis.

Across payload categories, fine-tuned models outperformed baseline Enformer predictions, especially for complex sequence modification such as DHS ordering and inversion (**Fig. 6C**). To confirm performance of the fine-mapped model on a held-out test set, we analyzed a second dataset comprised of 86 synthetic payloads including further *Sox2* DHS combinations, TF deletions and relocations, and combinations of DHS from *Sox2*, *Nanog*, *Sall1*, and *Prdm14* (Ordonez et al. 2025). Analysis of showed similar improvements (**Fig. 6D**). When comparing baseline and fine-tuned genome-wide accessibility predictions, fine-tuning eliminated several baseline false positives (**Fig. 6E**). Genome-wide assessment of mESC accessibility predictions demonstrate that fine-tuned models presented higher peak correlation to source DNase-seq and higher peak calling accuracy (**Fig. 6F-G**). Predictions from the fine-tuned Enformer retained high predictive performance across multiple assay categories (**Supplemental Fig. S11**). The final tuning step for Enformer was towards the human dataset (Avsec et al. 2021a), which may have slightly favored detection of human regulatory features. Fine-tuned predictions outperformed baseline for all mouse assays and showed only minimal performance decay against human assays (**Supplemental Fig. S12**). The fine-tuned model showed no improvement in predicting distance effects, suggesting the potential for future work to explicitly incorporate distance effects when training new models (**Supplemental Fig. S13**). Our results demonstrate a unique approach to iteratively improve the predictive performance of pre-trained deep learning models through incorporation of high-yield synthetic regulatory assay data.

## Discussion

We showed that Enformer has good performance modeling basic aspects of synthetic regulatory data at both *Sox2* and *α-globin* loci, but it fails to capture DHS synergy and was only partially sensitive to TF site deletions. For sequences (i.e. SynHPRT1R) that were more diverged from the reference used to train Enformer, the model performed poorly. We found Enformer unable to effectively model long-range regulation of *Sox2* transcription, consistent with previous reports that predictive performance falls off rapidly distal to the target TSS (Avsec et al. 2021a; Karollus et al. 2023; Toneyan and Koo 2024). The architecture of Enformer penalizes feature interaction proportionally to the distance between elements, which may contribute to this shortcoming. Further, long-range interactions are relatively uncommon thus will not be favored during training (Karollus et al. 2023). This highlights the limitations imposed by a limited training sequence space and poses caveats to future analysis of novel sequences.

We generally found better predictive performance using predictions for DNA accessibility, even when the experimental data measured expression. In particular, Enformer CAGE predictions showed overall poor performance on the distal regulatory elements we tested. The mESC CAGE output head performed as well as those from the Enformer paper (**Supplemental Fig. S3**), but it is possible that CAGE predictions in mESC could be further optimized. Our results suggest that accessibility provides a useful intermediate more amenable to prediction, at least in the context of distal enhancer perturbations. This is consistent with other observations that predictions of DNA accessibility are more effective than those for CAGE for promoter-distal regulatory elements (Karollus et al. 2023; Martyn et al. 2025). However, there is the possibility that this strategy overlooks differences between molecular function on the level of DNA accessibility and transcription. For example, while we used *Sox2* expression data to fine-tune Enformer, we only evaluated predictive performance on DNA accessibility. While the focus on DNA accessibility was supported by our earlier analysis, we did not show directly that fine-tuning makes more accurate gene expression predictions. For models like Enformer which are trained to predict both DNA accessibility and transcription, it remains to be seen whether the model has a deep understanding of the distinct underlying biomolecular processes involved, or rather if a core understanding based on regulatory sequence motifs underpins superficial differences which tailor the output to a specific genomic track. Our work suggests that different model architectures and training data may be needed to predict local effects of functional variants vs. identification of their target genes.

Multitask training to predict multiple regulatory genomic assays across different cell and tissue types enables information sharing between the different contexts (Kelley et al. 2018). However, performance remains limited by the training sequence space. Here we demonstrated that predictive power is related to the degree of DHS activity across multiple cell and tissue types, consistent with previous reports (Kathail et al. 2024). Broadly active sites across regulatory assays and cell types are favored during training, since prediction errors for such features are weighed more strongly. Thus correlation structures among genomic features can impede model training (Whalen et al. 2022). Simply expanding the number of training tasks has limited benefit as the sequence space explored remains the same (Schreiber et al. 2020; Kathail et al. 2024).

Deep learning models are promising tools to expand our understanding of regulatory genomics and to interpret variant functional impact (Avsec et al. 2021a, 2021b; Zhou and Troyanskaya 2015). However, our results show they are currently limited by the limited scope of their training sequences, which largely reflect the reference genome. We argue that deep learning models would benefit from training on functional characterization of regulatory elements that have been systematically deleted, reordered, and inverted, and perturbed by disease- and trait-associated variation. An iterative training strategy in which these results feed into model development, which in turn directs future experimental exploration would allow systematic exploration of disease- and trait-relevant sequence space and improve model performance across contexts. As a proof of concept of this strategy, our results showed that fine-tuning improved prediction performance across all payload categories and improved genome-wide prediction performance, cleaning many false positives signals and producing higher peak calling accuracy. Thus, synthetic regulatory genomics and machine learning are highly complementary in genomics’ dual status as both a “big data” and experimental science to uniquely permit pushing the limits of model knowledge while maintaining generality through iterative large-scale testing of model predictions.

## Methods

### Enformer model

The published Enformer model is hosted in the kaggle repository (https://www.kaggle.com/models/deepmind/enformer) cited in (Avsec et al. 2021a). This model, referred to here as Avsec2021, requires a 393,216-bp input sequence and outputs predictions for the central 114,688-bp region (**Supplemental Fig. S1A**).

Our model implementation was based on an equivalent model hosted at the DeepMind github repository (https://github.com/google-deepmind/deepmind-research/tree/master/enformer) and pre-trained weights from Google storage (gs://dm-enformer/models/enformer/sonnet_weights/). Based on previous work showing that Enformer retains most of its predictive performance with shorter input windows (Karollus et al. 2023), we employed an adapted version lacking an internal cropping layer to allow a flexible input sequence size instead of the fixed 196,608 bp, and to generate predictions for the whole input (**Supplemental Fig. S1B**). We compared the Avsec2021 version to our adapted model with decreasing input sizes (196,608 bp, 114,688 bp, and 25,600 bp). We selected ten 114,688 bp sites containing at least one mESC_CJ7 hotspot V1 (FDR 0.01) and predicted their mESC_CJ7 DNase-seq density using all four implementations (**Supplemental Fig. S1C**). We found no significant prediction deviations between Avsec2021 and our adapted versions (**Supplemental Fig. S1D-E**). Furthermore, we found high correlation in predicted DNase-seq for delivered *Sox2* payloads using 25,600-bp and 196,608-bp windows (**Supplemental Fig. S2**).

Enformer predictions throughout the manuscript were measured as the mean of ten 1-bp strides (5 bp downstream and upstream) unless otherwise noted. Predicted accessibility and CAGE signal were measured as the maximum signal across all bins overlapping the promoter or DHSs as described. Synthetic sequences used for prediction are included in **Supplemental Data S1**.

### Synthetic regulatory biology datasets

#### Sox2

Expression data was previously published: TableS4 (Brosh et al. 2023) and (Ordonez et al. 2025). Briefly, engineered allele *Sox2* expression was measured by qRT-PCR using allele-specific primers. Fold-change was calculated as 2^ΔCT[CAST-B6]^ and scaled to yield expression values ranging from 0 (ΔSox2) and 1 (WT). Previously published DNase-seq data was taken from ENCODE: DS13320 (mESC_CJ7) and DS21450 (mESC_E14) (Vierstra et al. 2014).

When employing Enformer to predict mESC_CJ7 DNase-seq signal of these payloads, virtual sequences were built by replacing either the full 143-kb *Sox2* locus or 43-kb LCR in the mouse reference genome (mm10; **Supplemental Table S1**) with the synthetic payload sequence. Predictions were generated for a 196,608-bp window centered at the payload insertion site (**Supplemental Table S2**).

### α-globin

Expression data was previously published: Fig5A and Fig6B-D (Blayney et al. 2023). Engineering was performed in mESC (mDist cells derived from mESC_E14), and expression measured in embryonic body-derived erythroid cells. *Hba* expression in engineered cells was profiled by qRT-PCR, normalized to *Hbb* and scaled to a proportion of WT expression. Previously published embryonic liver DNase-seq data was taken from ENCODE: DS20937 (mfLiver-F3) (Vierstra et al. 2014) and ATAC-seq of engineered cells from GEO: GSE219056.

We employed Enformer to predict accessibility of those five enhancers (R1-R4; **Supplemental Table S1**) in mouse embryonic liver (mfLiver-F3) as a proxy for *Hba* expression. Different enhancer configurations were generated by deleting and relocating R1-R4 enhancers on the mouse reference genome (mm10). Predictions were generated for a 25.6-kb window centered around the R1-R4 enhancers.

### HPRT1R

ATAC-seq data was previously published: BS15734A, BS15738A, BS21951A were taken from GEO (GSM8001814, GSM8001817, and GSM8001825, respectively) (Camellato et al. 2024). Previously published DNase-seq data was taken from ENCODE: DS13320 (mESC_CJ7), and DS18873 (H7_hESC) (Vierstra et al. 2014; Meuleman et al. 2020).

Virtual sequences were built by replacing the mouse *Hprt* locus (**Supplemental Table S1**) in the mouse reference genome (mm10) with each of the three payloads. Accessibility in mESC_CJ7 was predicted in a 393,216-bp window centered at the *Hprt* locus. We similarly predicted human *HPRT1* locus accessibility in human H7_hESC.

### Training Enformer on mESC CAGE data

Previously published mESC CAGE tracks CNhs14104 and CNhs14109 were taken from the FAN-TOM website at https://fantom.gsc.riken.jp/5/datafiles/reprocessed/mm10_latest/basic/mouse.timecourse.hCAGE/ (Fraser et al. 2015), and GSM3852792, GSM3852793, and GSM3852794 were taken from GEO series GSE132191 (Bonetti et al. 2020). We trained a new output head that employs the Enformer trunk to predict these 5 new mESC CAGE tracks. The output head was trained, evaluated and tested on random 25.6 kb genomic intervals using a Poisson negative log-likelihood loss function. Training and evaluation datasets composed of 469,585 intervals from the mouse reference genome (mm10) randomly assigned in a 9:1 ratio to each dataset respectively. We used Adam optimizer from TensorFlow with a learning rate of 0.0005 and otherwise default settings to train for 20 epochs, each visiting 1,600 intervals in batches of 4 intervals. Only the new output head weights were updated during training, while preserving weights for the Enformer trunk.

Resulting model performance was evaluated by measuring Pearson’s correlation of CAGE signal (log(1+x) transformed) between original and predicted CAGE signals at nine ∼5.7-Mb genomic intervals of interest (**Supplemental Table S1, Supplemental Fig. S3**) in non-overlapping 196,608-bp blocks. We also measured Pearson’s correlation of seven other CAGE tracks from baseline Enformer at the same intervals (FANTOM: CNhs10466, CNhs10469, CNhs10471, CNhs10474, CNhs11297, CNhs12107, and CNhs13511; Enformer index 6601, 6604, 6606, 6609, 6823, 6830, and 6937). GSM3852792 was selected as representative for further analysis in mESC. Predictions used a 196,608-bp input window.

### Enformer fine-tuning

To fine-tune Enformer using the *Sox2* LCR synthetic payload dataset, we added a new output head to Enformer to predict *Sox2* expression based on *Sox2* LCR sequence. The new head employs a self-attention layer to capture relevant features from the Enformer prediction trunk independently of position. The resulting signal is combined into a scalar prediction of *Sox2* expression by a fully-connected layer using a sohttps://ftplus activation function (**Fig. 6A**).

Here we evaluated three configurations of the self-attention layer by varying the internal key/value matrix sizes, and number of independent attention projections (or heads). SingleHead 64/64 applies a single projection of 64 key and value matrices, SingleHead 64/128 applies a single projection of 64 key and 128 value matrices, and MultiHead 64/64 applies four independent projections of 64 key and value matrices.

Brosh 2023 and Ordoñez 2025 synthetic payload sequences were expanded to 1,280 entries by striding 1 bp at a time up to ±640 bp upstream and downstream through a 25.6-kb focal region centered at the *Sox2* LCR (**Supplemental Fig. S9A**). We included all Brosh 2023 payloads (n=70) during both training and evaluation. 48 strides from Brosh 2023 dataset were sampled to train the three fine-tuned model configurations and another 48 strides were sampled for evaluation (**Fig. 6B, Supplemental Fig. S9B**). For the 86 payloads in the held-out test Ordoñez 2025 dataset, 32 strides were sampled as a testing dataset.

In parallel, we trained models with Brosh 2023 payloads partitioned into training and evaluation datasets to avoid data leakage. Payloads were randomly assigned into a training or validation dataset in a 3:1 ratio for ten independent batches. Models were trained and their performance evaluated using each batch training and validation datasets (**Supplemental Fig. S9C**). Performance was average across training batches. Retrained models presented similar results to full dataset training, but higher variance likely attributable to the reduced size of the training sets.

For both strategies, fine-tuning was conducted using only the baseline Enformer trunk and the new output head. This strategy allows us to adjust Enformer internal feature extraction based on our synthetic dataset, while preserving mouse and human prediction heads. We employed the Adam optimization algorithm with a learning rate of 10^-5^ to fine-tune the models for 10 epochs, each visiting 400 sequences at time in batches of 4 sequences. Training error was measured by Mean-Squared Error (MSE).

Predictions for performance comparisons were made using a 25.6-kb input window. Expression was estimated directly from the new output head and indirectly by summing predicted accessibility of all LCR DHSs in mESC_CJ7. For comparison, accessibility estimates were scaled to match measured expression using a linear regression fitted with the training dataset. Predictions were averaged across all payload strides and error measured by Mean Squared-Error (MSE) across synthetic payloads (**Supplemental Table S1**).

### Genome-wide predictive performance

Genome-wide evaluations were conducted by comparing experimental DNase-seq results to baseline and fine-tuned models for 8 selected tracks (**Supplemental Table S4**). For each selected track, we evaluated predictions at 2,000 randomly selected 25.6-kb regions from autosomes containing at least one hotspot V1 (FDR 0.01) for the selected track. Within those regions, peaks were identified as continuous runs of DNase-seq density or prediction values above the track cutoff and larger than 128 bp. The track signal cutoff was established by calculating the signal 95^th^ percentile at 2,000 random 128-bp sites within the targeted regions. A final collection of evaluated sites was generated by merging peaks called for all DNase-seq assays (selected track, technical replicate and others), baseline and fine-tune predictions and combining it to non-overlapping random sites used to the signal cutoff. Activity across cell-types and tissues was estimated by annotating the number of tracks with overlapping hotspots V1 (FDR 0.01) among a collection of reference mouse (N=36) and human (N=105) ENCODE DNase-seq tracks (**Supplemental Table S3**).

## Software availability

Code for training Enformer on new cell types and for fine-tuning Enformer is available at https://github.com/mauranolab/finetune-enformer and as Supplemental Code. Weights for the fine-tuned Enformer and mESC CAGE predictions are available at https://doi.org/10.5281/zenodo.13363228.

## Competing interest statement

M.T.M. is listed as an inventor on a patent application describing Big-IN.

## Acknowledgements

We thank Raquel Ordoñez for helpful comments, and Ran Brosh and Brendan Camellato for sharing data. This work was partially funded by National Institutes of Health (NIH) grants RM1HG009491 and R01MH136353.

## Author contributions

M.T.M conceived and oversaw the project. A.R.S. performed analyses. A.R.S. and M.T.M wrote the paper.

## Notes

### Summary of Updates

Some minor updates and clarifications throughout the document

