## Supplemental Material for "Iterative improvement of deep learning models using synthetic regulatory genomics"

### Supplemental Figures

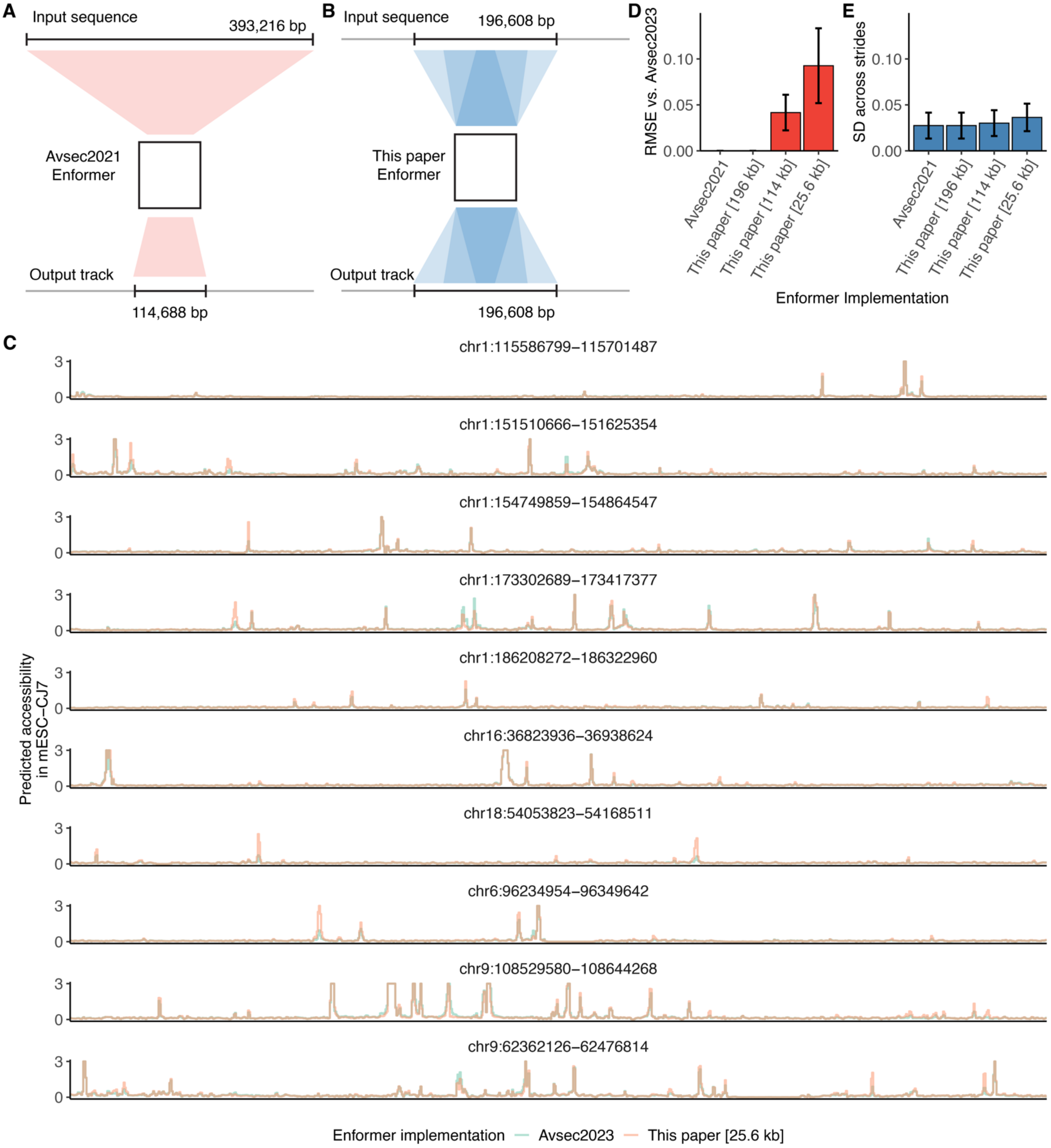

**Supplemental Fig. S1. Baseline and adapted Enformer prediction comparison.**

Comparison of Avsec2021 and adapted Enformer implementation in this paper to ensure no major differences were introduced. Here, we evaluated mESC_CJ7 accessibility prediction for these implementations at ten randomly selected 114,688 bp sites containing at least one mESC_CJ7 hotspot V1 (FDR 0.01). To accommodate larger input sequence requirements, we expanded the targeted sites to the required size. For smaller input sequence requirements, the targeted regions were broken down into non-overlapping windows. (A) Schematic of Avsec2021 baseline model (Avsec et al. 2021a). (B) Schematic of model in this paper (Methods). (C) Accessibility predictions for all evaluated sites for both Avsec2021 and this paper [25.6 kb]. (D-E) Comparison of Enformer implementation performance. Error bars indicate standard deviation across evaluated sites. (D) Deviation from Avsec2023 baseline measured by Root Mean Squared Error (RMSE). (E) Prediction standard deviation across ten 1-bp strides used to estimate accessibility.

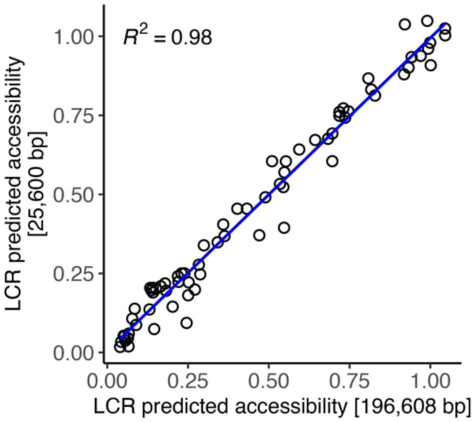

**Supplemental Fig. S2. Enformer accessibility predictions of *Sox2* payloads by input window size.**

Comparison of Enformer accessibility predictions using a 25,600 bp window centered at *Sox2* LCR and a 196,608-bp window centered at whole *Sox2* Locus. Each point represents accessibility prediction of a synthetic payload delivered in place of *Sox2* LCR. Predicted accessibility was measured as the summed signal of all included LCR DHSs (19-28) in MESC_CJ7. Blue line indicates linear regressions.

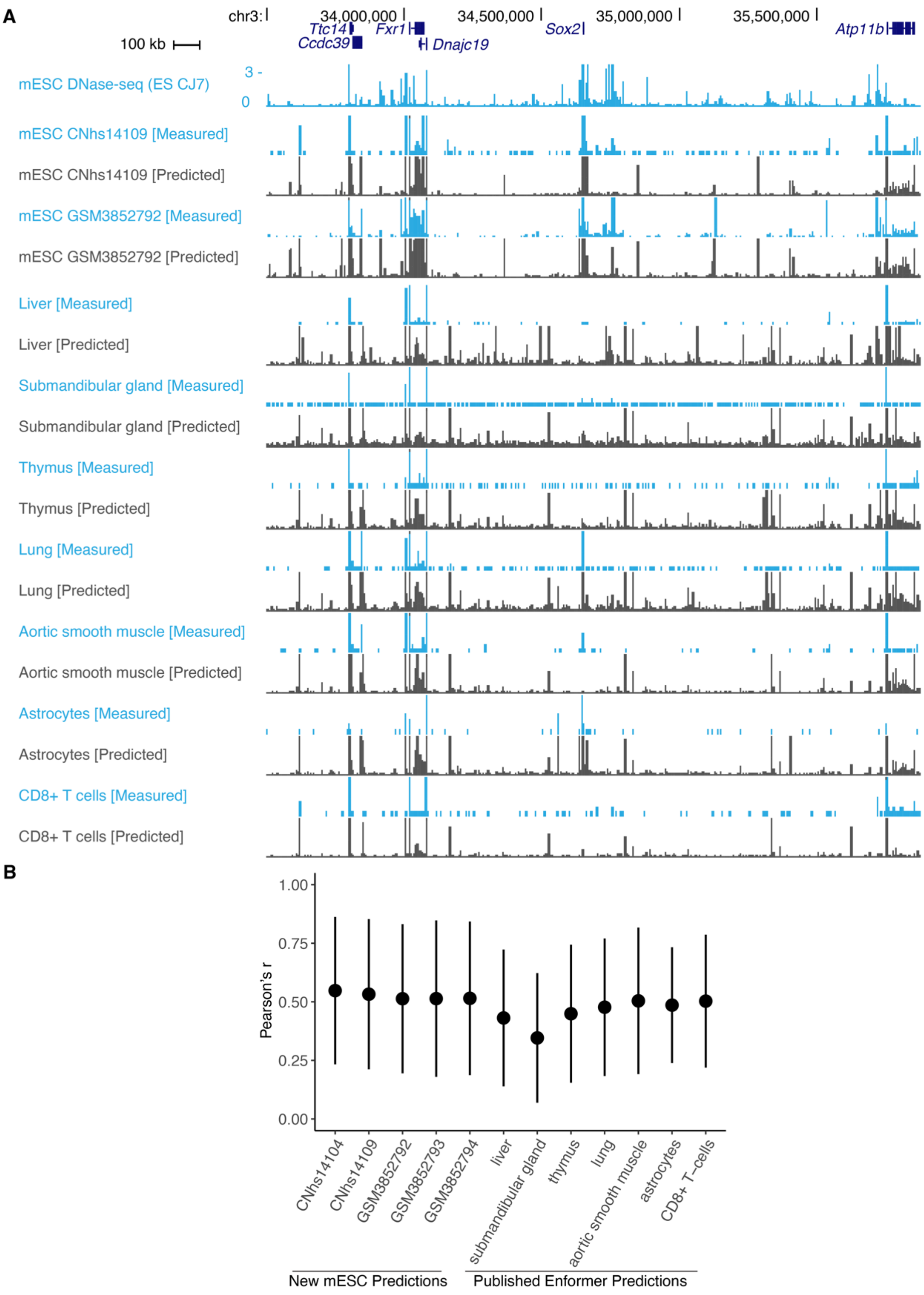

**Supplemental Fig. S3. Comparison of mESC CAGE predictions to training data.**

(A) Example browser shot showing experimentally measured and predicted signal for new mESC CAGE and published Enformer cell types at the wider *Sox2* locus. (B) Mean correlation between predicted and measured CAGE signal across test intervals (**Supplemental Table S1**).

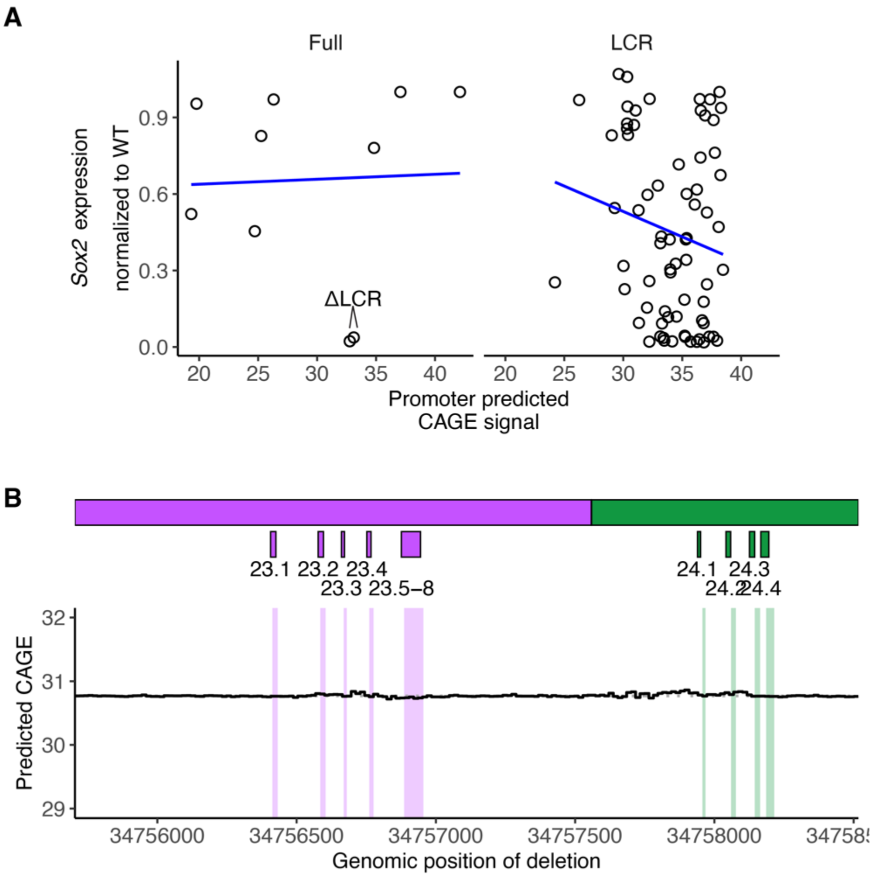

**Supplemental Fig. S4. Analysis of CAGE predictions at *Sox2*.**

(A) Comparison of experimentally measured *Sox2* expression and Enformer predicted CAGE signal at *Sox2* promoter. Shown are synthetic payloads replacing the full *Sox2* locus (left, n=10) or *Sox2* LCR (right, n=70). Expression was characterized by allele-specific qRT-PCR and scaled between 0 (ΔSox2) and 1 (WT). CAGE signal was measured as the maximum signal at *Sox2* promoter. Blue lines indicate linear regressions on data shown in each panel. (B) In silico scanning deletion analysis of *Sox2* expression. Enformer was used to predict CAGE signal at *Sox2* promoter for a series of DHS23-24 virtual payloads replacing the *Sox2* LCR. 16-bp scanning deletions were encoded by replacing the payload sequence with Ns sliding across the sliding across the length of DHS23 and DHS24. Black line indicates *Sox2* promoter maximum CAGE signal for each deletion position (x-axis). Horizontal dotted line indicates baseline expression. Boxes above indicate relevant TF recognition sequences. Compare to **Fig. 2A**.

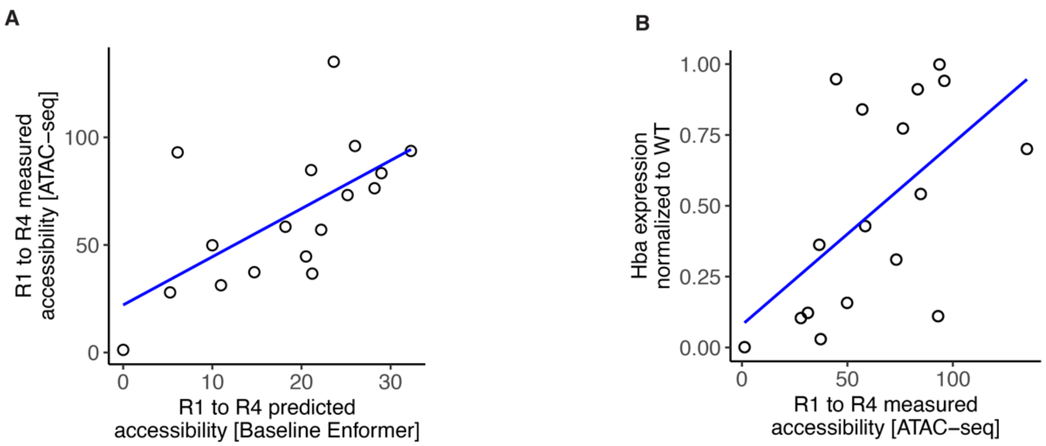

**Supplemental Fig. S5. Comparison of predicted and experimentally measured accessibility.**

Comparison with Enformer predicted accessibility of mouse embryonic liver (A) and *Hba* expression (B). Each point represents an *Hba* enhancer configuration from Blayney et al (2023) (N = 17). Predicted and measured enhancer accessibility were calculated as the sum of the maximum signal at each *Hba* enhancer (R1 to R4). Blue line indicates linear regression.

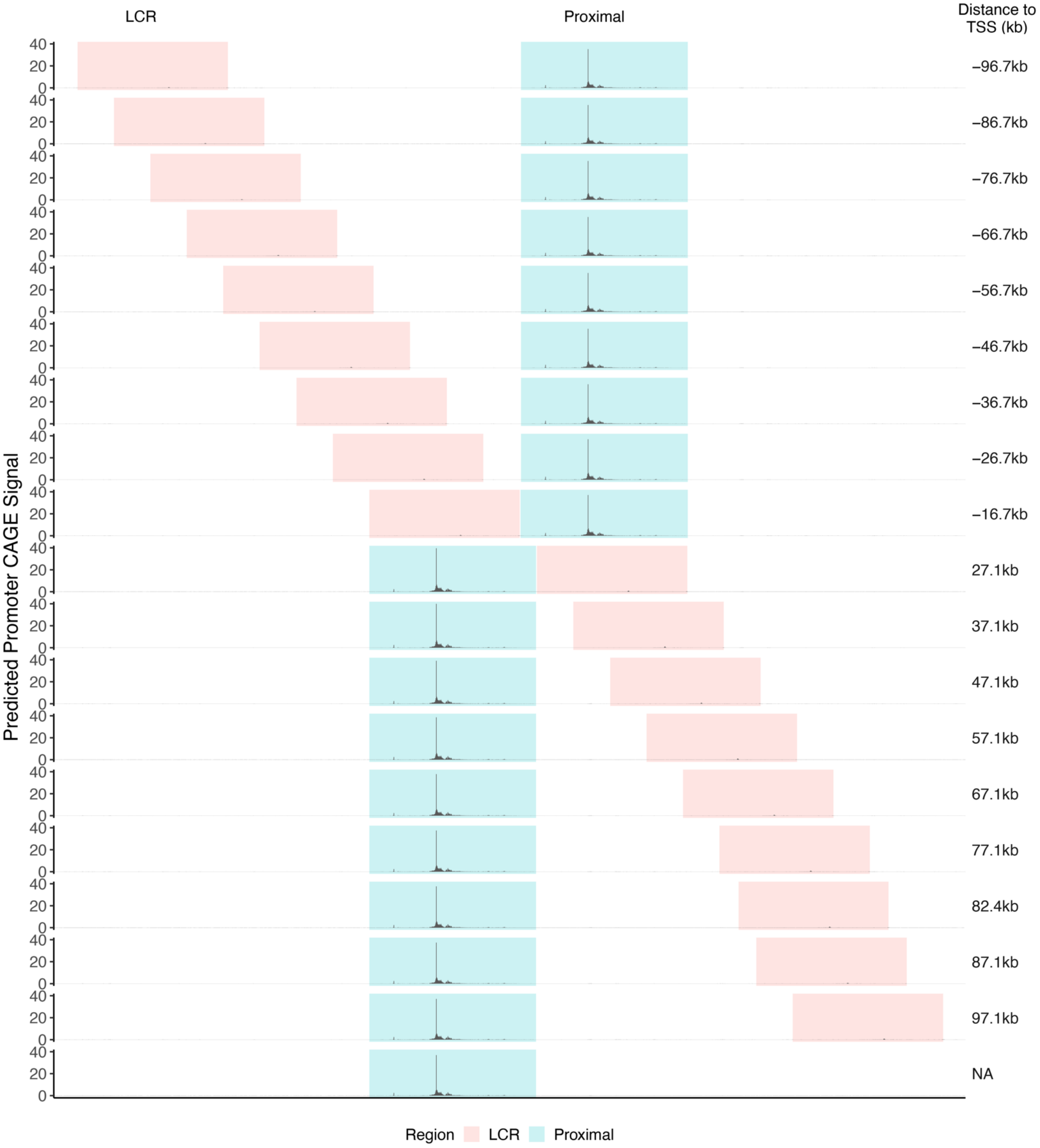

**Supplemental Fig. S6. Enformer predictions of in-locus *Sox2* LCR relocation.**

(A) Enformer prediction of in locus LCR relocation tracks. *Sox2* proximal region is highlighted in blue and LCR highlighted in gray. LCR distance to *Sox2* TSS is indicated to the right.

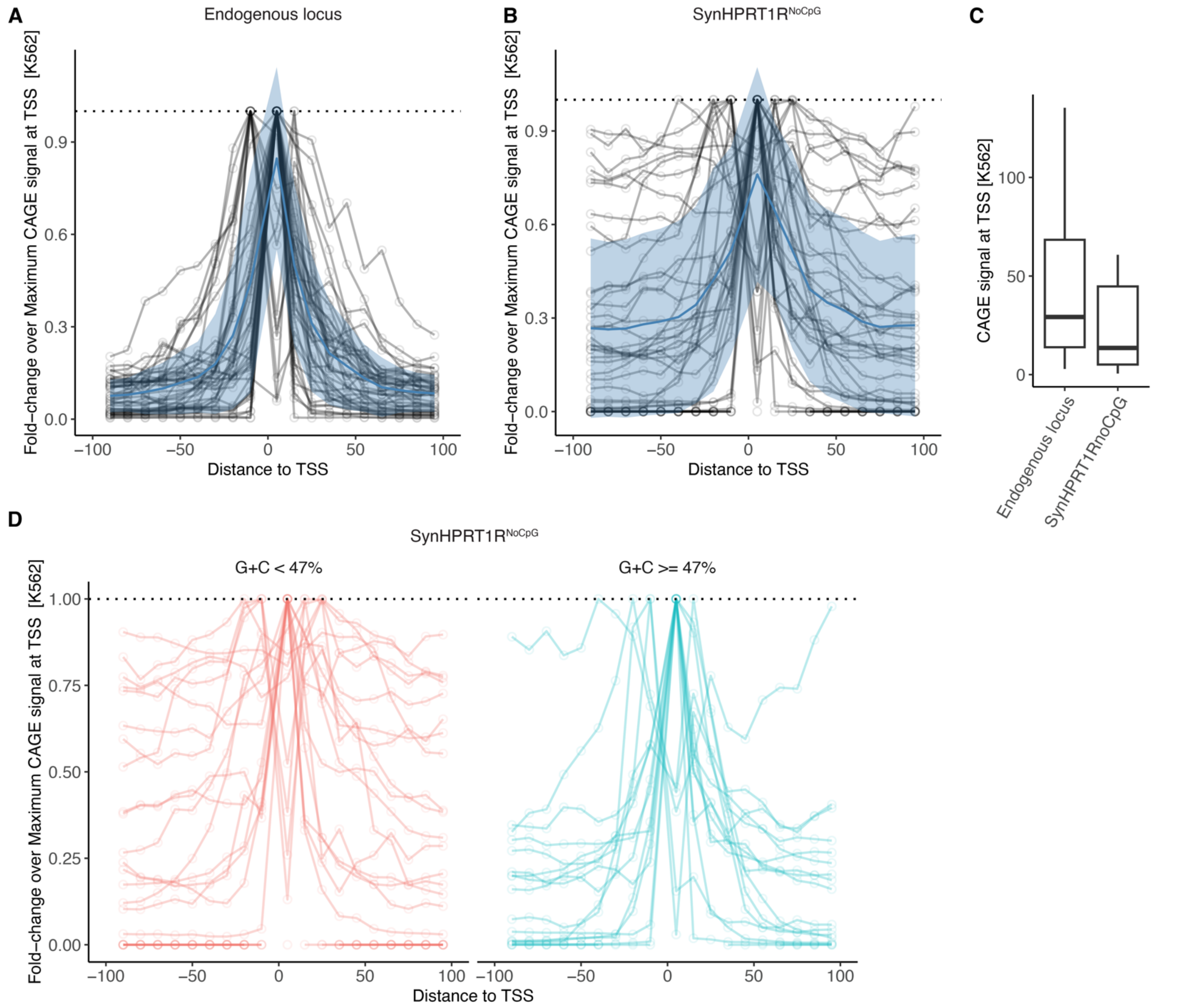

**Supplemental Fig. S7. Impact of background sequence on Enformer distance effect predictions.**

(A-B) Enformer estimates of transcription driven by 74 K562 candidate CREs (Toneyan and Koo 2024) as a function of their distance to targeted gene TSS. Simulations were conducted by moving CRE regions 10 kb downstream and upstream from their targeted TSS and measuring K562 CAGE signal at the TSS. A 5-kb window centered at both CRE and TSS were preserved while the remaining background sequence was generated by shuffling dinucleotides from the CRE endogenous loci (A) or from SynHPRT1R$^{noCpG}$ (B). CAGE signal was estimated as the mean from 10 random dinucleotides shuffles and normalized by the maximum estimated signal across all positions. Each line represents a CRE candidate, and each point a CAGE signal estimate. Solid blue line represents the mean signal across all CREs and the blue shaded area ±1 s.d.. (C) Predicted K562 CAGE signal at TSS in both contexts. (D) Analysis from (B) split by whether G+C of entire 196,608-bp endogenous window is above or below 47%.

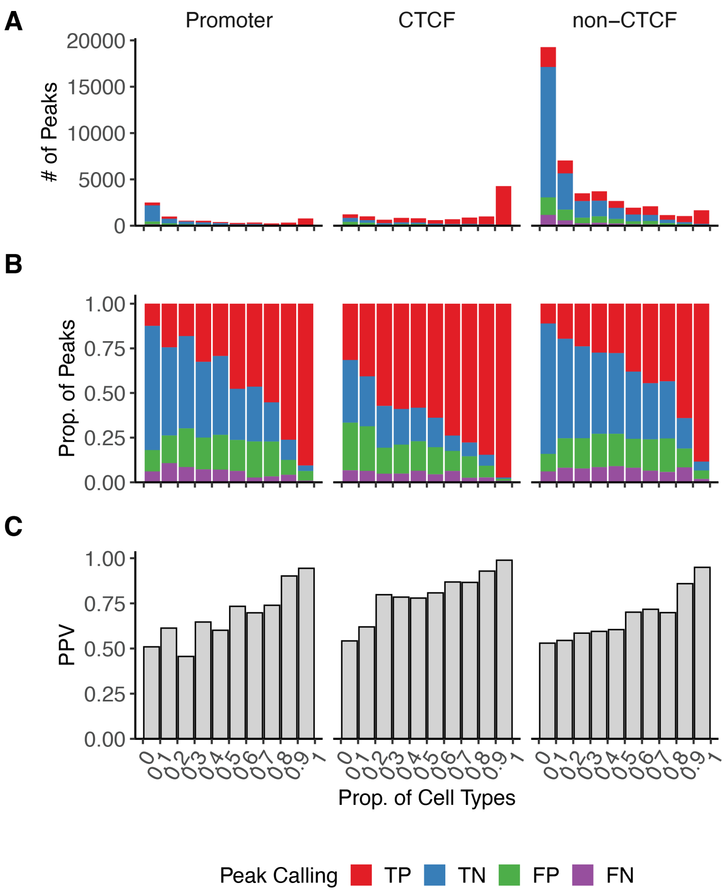

**Supplemental Fig. S8. Cell-type activity analysis at promoter, CTCF, and, non-CTCF sites.**

(A-C) Peak calling comparison of mESC_CJ7 DNase-seq track and Enformer prediction (detailed in **Methods**) of Promoter (within 2.5 kb of TSS), CTCF (overlapping CTCF hotspots) and non-CTCF sites. Peaks were binned according to the proportion of cell types represented in a collection of mouse (N=36) reference DNase-seq datasets (**Supplemental Table S3**). Peaks are classified as true positives (TP), when found in both reference DNase-seq and Enformer predictions; true negative (TN), when not found in either; false positive (FP), when found only in Enformer predictions; and false negative (FN), when found only in reference DNase-seq. (A) Total number of peaks colored by category. (B) Relative proportion of categories per bin. (C) Positive predictive value (PPV) per bin.

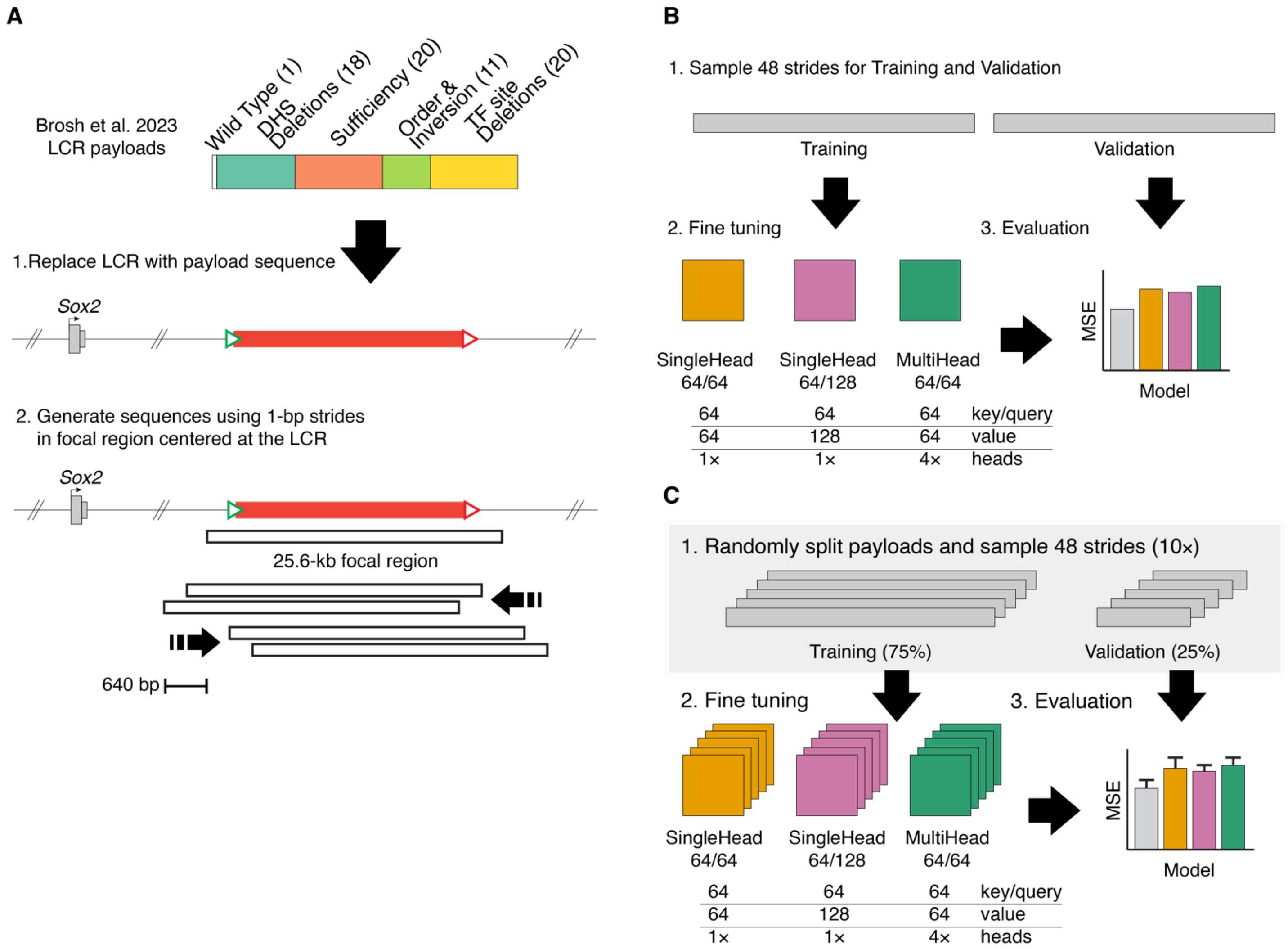

**Supplemental Fig. S9. Schematic of fine-tuning training and dataset augmentation.**

(A) *Sox2* LCR dataset augmentation strategy. Virtual input sequences were built by replacing *Sox2* LCR by each synthetic payload sequence. Dataset was augmented by striding a 25.6-kb focal region centered at the LCR 1 bp at a time (upstream and downstream) into a total of 1,280 sequences. (B-C) Fine-tuning strategy using all payloads (B) and partitioning payloads into training and validation datasets (C). (B) Fine-tuning strategy including all payloads. For each payload, 48 strides were randomly selected for training and another 48 strides were selected to evaluate the trained models. (C) Fine-tuning strategy on partitioned payloads. Payloads were randomly partitioned into training and validation datasets in ten independent batches and 48 strides of each payload selected. For each batch, models were trained using the training dataset and evaluated with their respective validation dataset.

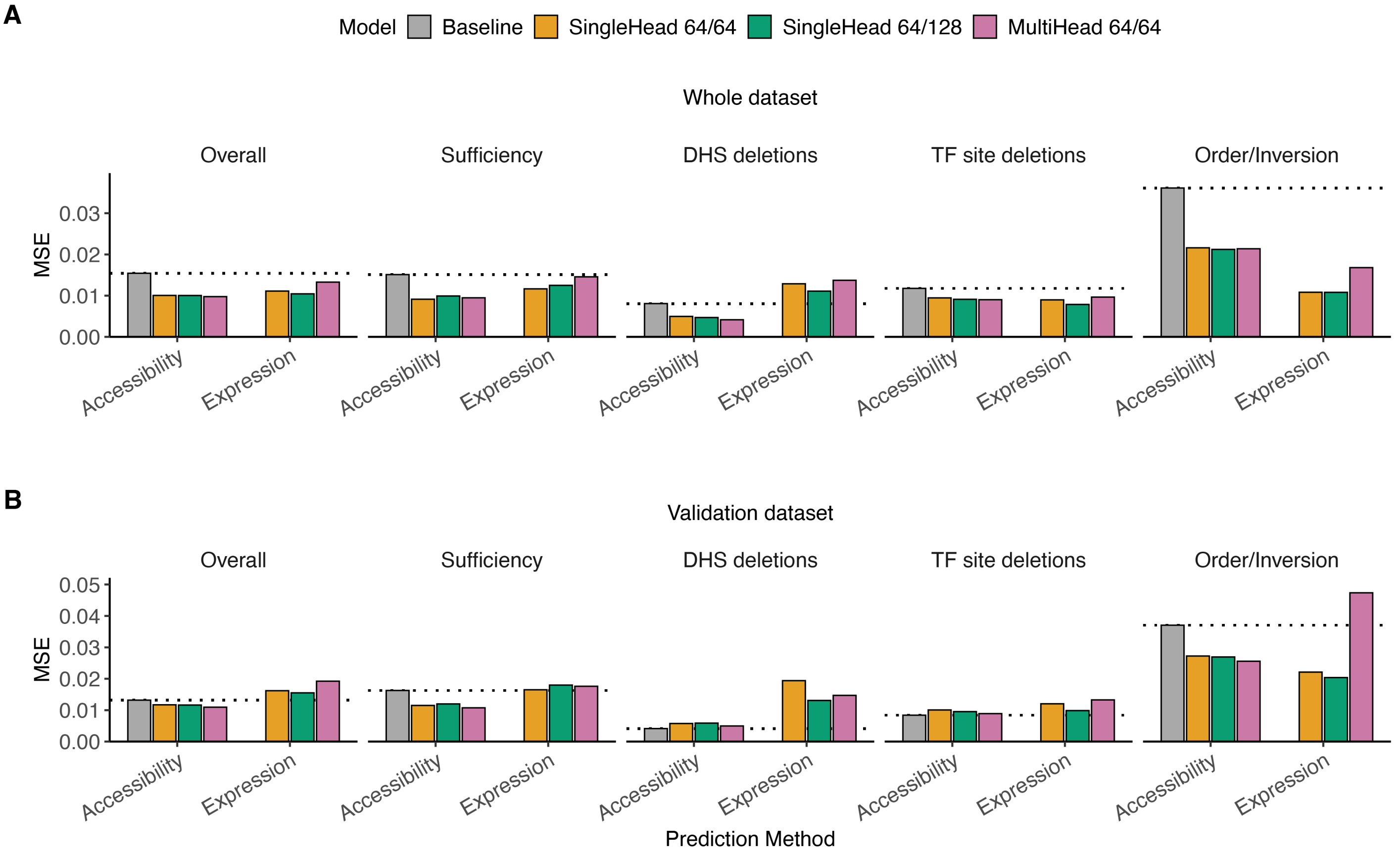

**Supplemental Fig. S10. Partitioned fine-tuning training performance.**

(A-B) Prediction performance of baseline and fine-tuned models trained using payload partitioning strategy. Expression was estimated either by summing the maximum predicted accessibility of all LCR DHS (Accessibility) or directly from the new output head (Expression). Model performance was measured as mean squared error (MSE) and averaged (mean) between ten independent partition batches. Presented the results for the overall dataset and payload categories with more than two payloads. Horizontal dotted lines indicate baseline performance. (A) Shows prediction performance for the whole training dataset, and (B) performance for the validation dataset.

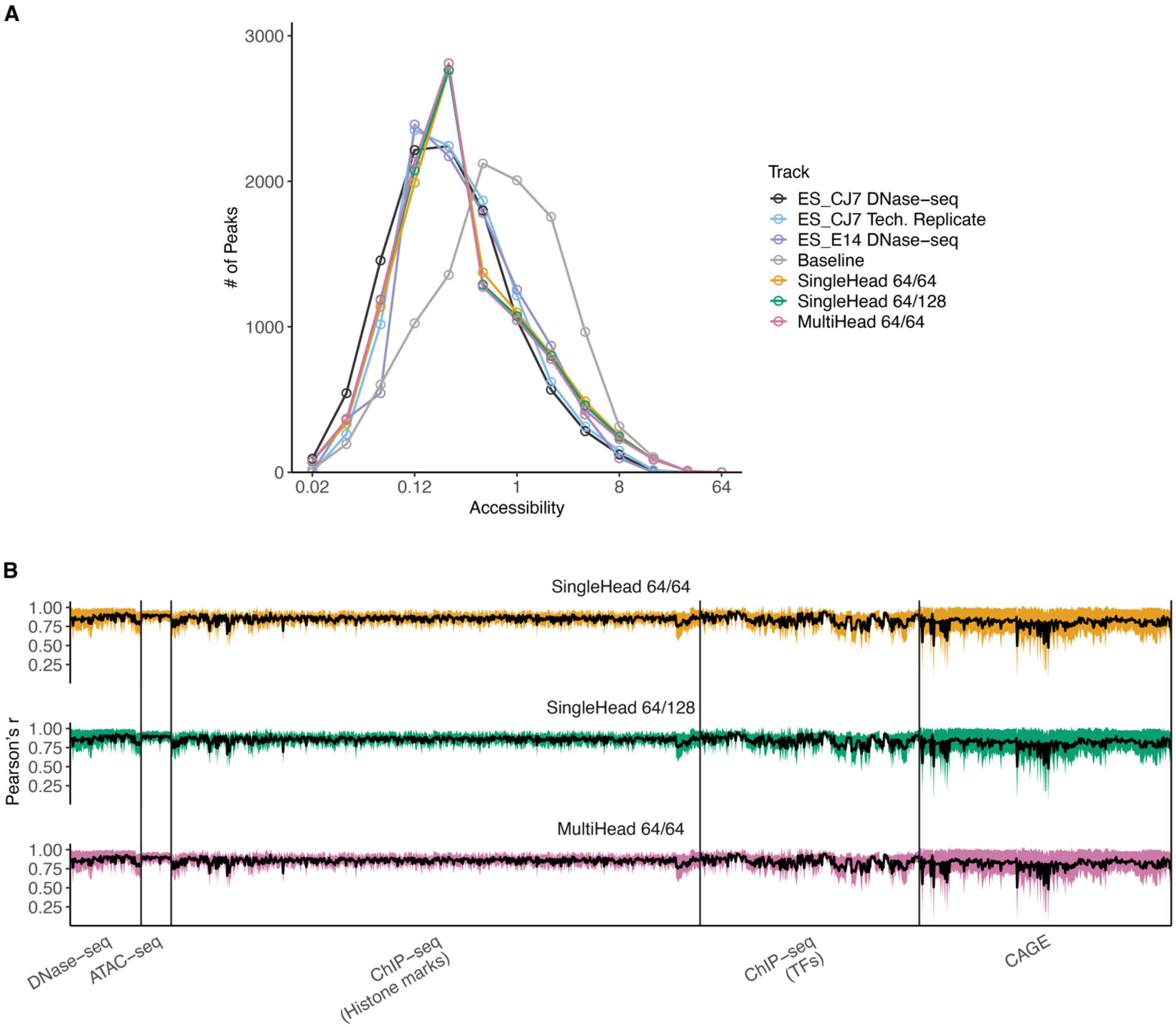

**Supplemental Fig. S11. mESC_CJ7 peak analysis.**

(A-B) DNase-seq density and Enformer prediction comparison at 2,000 randomly selected 25.6-kb regions containing at least one mESC_CJ7 hotspot V1 (FDR 0.01). (A) Peak signal distribution at randomly selected regions. Peak calling detailed in **Methods**. Peak signal was measured as maximum DNase-seq density or mESC_CJ7 predicted accessibility. Shown the signal distribution for DNase-seq data for mESC_CJ7, a technical replicate, and an independent cell line (mESC_E14), and Enformer predictions. (B) Fine-tuned models prediction correlation to baseline Enformer at randomly selected regions. Shown Pearson's correlation score (*r*) of fine-tuned models to baseline model prediction for all mouse output tracks (single-cell ATAC-seq were excluded). Black line indicates mean *r* across all evaluated regions and shaded area represents ±1 s.d.. Tracks are grouped according to their type indicated in x-axis.

**A**

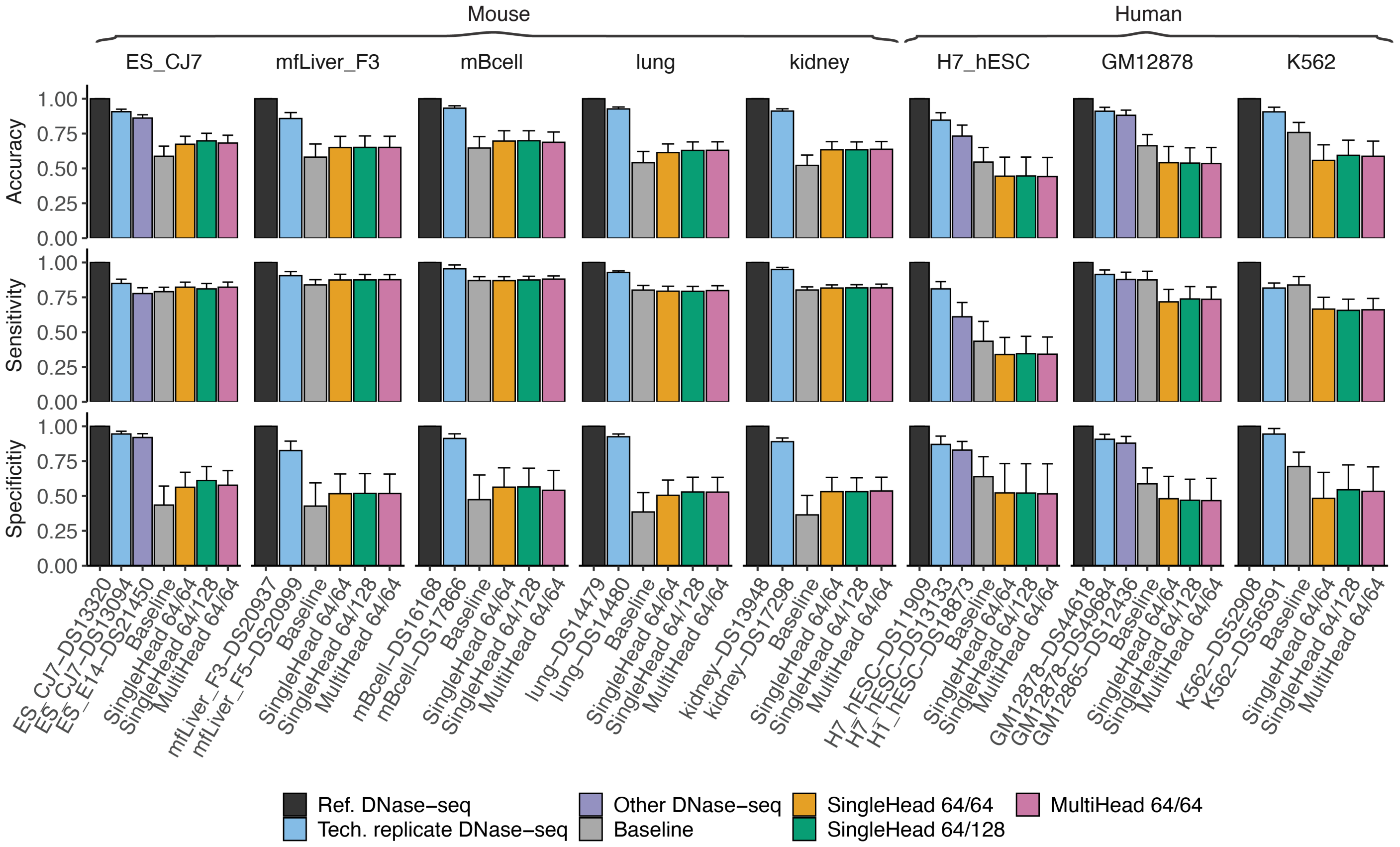

**Supplemental Fig. S12. Peak calling performance across multiple cells and tissues.**

(A) Peak calling accuracy, sensitivity (true positive ratio) and specificity (true negative ratio) for various targeted ENCODE cell/tissues DNase-seq tracks. Peak calling is detailed in **Methods**. Results shown for the targeted DNase-seq track, a technical replicate, an independent similar track, and predictions made by baseline and fine-tuned Enformer. Error bar indicates ±1 s.d..

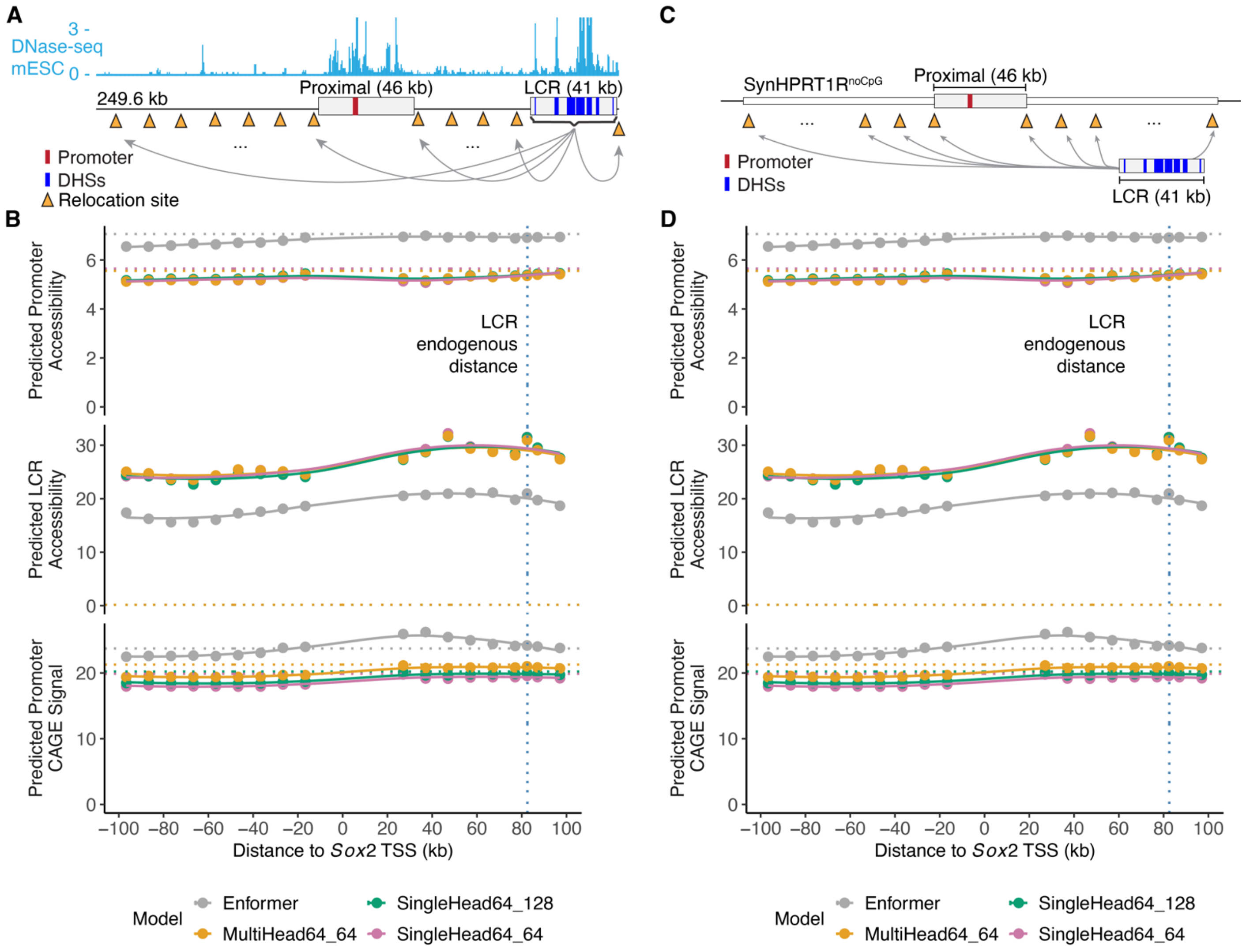

**Supplemental Fig. S13. Fine-tuned Enformer model does not predict effect of LCR relocation on *Sox2* accessibility or expression.**

(A,C) Schematics of relocation simulation of the *Sox2* LCR in 10-kb intervals surrounding the promoter in its endogenous locus (A) or neutral synthetic sequence (SynHPRT1R$^{noCpG}$) (C); red rectangles indicate *Sox2* promoter and blue rectangles LCR DHSs 19-28. (B, D) Enformer predictions for a 249.6-kb window encompassing the whole virtual locus and any remaining space was filled with Ns. Each point represents predicted accessibility of *Sox2* promoter (top) or LCR (middle), or promoter CAGE signal (bottom) for a given distance between *Sox2* TSS and LCR. Promoter CAGE signal and accessibility was measured as the site maximum predicted signal, and LCR accessibility as the sum of DHSs 19-28 maximum predicted signal. Vertical black dotted line indicates endogenous LCR distance to TSS. Horizontal dotted lines indicate prediction when LCR was replaced by SynHPRT1R$^{noCpG}$ fragment of the same size. Solid blue lines show LOESS fits. Line colors correspond to model, either baseline Enformer or the three fine-tuned Enformer models described in **Fig. 6**.

### Supplemental Tables

#### Supplemental Table S1. Genomic coordinates.

Genomic coordinates of DHSs and other relevant regions.

#### Supplemental Table S2. *Sox2* and *α-globin* accessibility predictions.

Enformer accessibility predictions of synthetic payloads delivered in place of *Sox2* LCR (Brosh et al. 2023) and CRISPR-Cas9 editing of *α-globin* enhancers (Blayney et al. 2023), and their experimentally characterized engineered gene expression (qRT-PCR). Payload.ID, unique descriptor of payload or CRISPR-Cas9 edit; Group, payload or edit category; Locus, indicator of *Sox2* or *α-globin* locus; Activity, normalized observed gene expression; AccessibilityPrediction, sum of DHSs maximum accessibility for all *Sox2* LCR DHSs 19-28 or *α-globin* DHSs (R1-R4); AdjustedAccessibilityPrediction, scaled sum of DHSs accessibility to fit gene activity (detailed in Methods); CAGEPrediction, predicted mESC CAGE signal at *Sox2* promoter.

#### Supplemental Table S3. ENCODE data used for cross-cell type activity analysis.

Collection of ENCODE mouse and human DNase-seq tracks employed to characterize cross-cell patterns. Name, cell or tissue label; DS and ENCODE, sample identifiers (see https://www.encodeproject.org); and Species.

**Supplemental Table S4. ENCODE tracks investigated.**

ENCODE tracks employed to compare baseline and fine-tune models. DS and ENCODE sample IDs are from https://www.encodeproject.org (Thurman et al. 2012; Vierstra et al. 2014; Meuleman et al. 2020). EnformerIndex lists track index among Enformer predictions. Replicate ID includes DS number of technical replicate; Other ID includes unique DS number for the other DNase-seq track to which the current was compared.

| Name | Sample ID | ENCODE ID | Species | Enformer Index | Replicate ID | Other ID |
|---|---|---|---|---|---|---|
| H7_hESC | DS11909 | ENCLB449ZZZ | human | 20 | DS13133 | DS18873 |
| GM12878 | DS44618 | ENCLB681AXO | human | 12 | DS49684 | DS12436 |
| K562 | DS52908 | ENCLB096YUZ | human | 625 | DS56591 | DS56591 |
| mESC_CJ7 | DS13320 | ENCLB163SYJ | mouse | 5323 | DS13094 | DS21450 |
| mESC_E14 | DS21450 | ENCLB890BTE | mouse | 5324 | DS18505 | DS13320 |
| mfLiver_F3 | DS20937 | ENCLB173ZFD | mouse | 5322 | DS20999 | - |
| MEL | DS13036 | ENCLB066PZV | mouse | 5339 | DS21963 | - |
| mBcell | DS16168 | ENCLB693PQW | mouse | 5315 | DS17866 | - |
| lung | DS14479 | ENCLB144OVE | mouse | 5338 | DS14480 | - |
| kidney | DS13948 | ENCLB388UBV | mouse | 5333 | DS17298 | - |

**Supplemental Data**

Supplemental Data S1 and Supplemental Code are provided as separate files.

**Supplemental Data S1. Evaluated synthetic sequences.**

Collection of synthetic sequences employed to evaluate and fine-tune Enformer models, including 166 synthetic sequences where *Sox2* or the *Sox2* LCR was replaced by the synthetic payloads (Brosh et al. 2023; Ordonez et al. 2025); 17 sequences where *Hba* locus was replaced by payloads (Blayney et al. 2023); 3 sequences where *Hprt* locus was replaced by payloads (Camellato et al. 2024); and 40 sequences from the *Sox2* relocation assay (**Fig 4**).

**Supplemental Code. Enformer fine-tuning code.**

Archive of code used to fine-tune Enformer and train Enformer on new CAGE data (also available from https://github.com/mauranolab/finetune-enformer).
