## Supplemental Code for "Iterative improvement of deep learning models using synthetic regulatory genomics": README.md

# Iterative improvement of deep learning models using synthetic regulatory genomics

We have included below a quick start guide broken down into the following sections:

- [Installation](#installation)
- [Training Enformer to output predictions for new tracks](#training-enformer-to-output-predictions-for-new-tracks)
- [Fine-tuning Enformer on Synthetic Regulatory Genomics data](#fine-tuning-enformer-on-synthetic-regulatory-genomics-data)

### Data availability

Pre-trained weights for the fine-tuned models and a new output head for five mESC (mouse embryonic stem cells) CAGE tracks are available under our [Zenodo repository](https://doi.org/10.5281/zenodo.13363228).

### Credits

This project Enformer implementation is based on their code available
[here](https://github.com/google-deepmind/deepmind-research/tree/master/enformer)
with only a minor modification to remove input/output size constraints (see
[enformer.py](src/models/enformer.py) at line 141).

### Citation

If you find this repository useful and use some of our code, please cite our
[preprint](https://www.biorxiv.org/content/10.1101/2025.02.04.636130v2):

> Ribeiro-dos-Santos, André M., and Matthew T. Maurano. Iterative Improvement
> of Deep Learning Models Using Synthetic Regulatory Genomics.
> bioRxiv, 21 Feb. 2025. bioRxiv, https://doi.org/10.1101/2025.02.04.636130.

---

## Installation

### Dependencies

We recommend using `python 3.9` due to several dependencies of Enformer original [code](https://github.com/google-deepmind/deepmind-research/tree/master/enformer).
All other library requirements are listed under `requirements.txt`.

To install them run the following:

```bash
## Create a virtual environment to isolate your packages
python3 -m venv .venv
source .venv/bin/activate
## Install requirements
pip install requirements.txt
## Check installation
python3 -m src.models.train --help
```

### Setup

The code requires two environment variables be defined, to locate Enformer
tensorhub and publication weights. They can be defined as follow:

```bash
## Address to Enformer tensorhub version. When using `tensorhub` model,
## the program will fetch and cache from the specified address.
ENFORMERTENSORHUB="https://www.kaggle.com/models/deepmind/enformer/TensorFlow2/enformer/1"
## Local path to Enformer publication weights from their
## [google storage](gs://dm-enformer/models/enformer/sonnet_weights/)
ENFORMERBASELINE="data/Avsec2021Weights/"
```

Other than those, you may consider define the following variables to simplify
usage:

```bash
SEQ_LENGTH=25600 ## Prediction input/output size in bp
SEQ_WINDOW=128   ## Prediction output bin size. Default produced by Enformer architecture

## Fine-tuned models attention layer specifications.
## When using our code, you may be required to define them whenever running a
## specific model.
## - SingleHead 64/64
SPEC_SH6464="--key-size 64  --value-size 64  --num-heads 1"
## - SingleHead 64/128
SPEC_SH64128="--key-size 64 --value-size 128 --num-heads 1"
## - MultiHead 64/64
SPEC_MH6464="--key-size 64  --value-size 64  --num-heads 4"
```

---

## Training Enformer to output predictions for new tracks

Given some bigwig files for some tracks we wish to make predictions, we can train an new output layer to generate similar predictions as Enformer without changing the model underlining weights.
For such purpose, we must first specify the new output tracks on a TSV such as the one below.

```tsv
# Example of tracks.tsv
index   identifier  file    clip    scale   sum_stat    description
0   CNhs14104   /path/to/data/CNhs14104.bw  384 1   sum mESC CAGE 0
1   CNhs14109   /path/to/data/CNhs14109.bw  384 1   sum mESC CAGE 1
2   GSM3852792  /path/to/data/GSM3852792.bw 384 1   sum mESC CAGE 2
3   GSM3852793  /path/to/data/GSM3852793.bw 384 1   sum mESC CAGE 3
4   GSM3852794  /path/to/data/GSM3852794.bw 384 1   sum mESC CAGE 4
```

All seven columns are required, specially `file`, `clip`, `scale`, and `sum_stat`.
These columns specify the following:
- `file` defines the path to the target track bigwig file.
- `clip` defines the value that clips each bin signal.
- `scale` defines a scaling factor to multiply each bin.
- `sum_stat` defines the function to aggregate the signal at each bin. We currently allow `mean` and `sum`.

### Build a training dataset

Next, we must construct a dataset of random genomic sequences and their respective track signals binned in 128 bp intervals.
Command below will construct such dataset while binning each track signal according to the `tracks.tsv` specifications.
It randomly samples regions within mappable intervals (specified below in `mappable.bed`) and measures the binned signal of each bigwig track.
Regions with standard deviation across bins below `{cutoff}` are excluded from the dataset.
This command will generate several output files, including a BED file (`{prefix}.bed`) with coordinates of every region included, and a H5 file for each dataset batch (`{prefix}.{0..batches}.h5`).

```bash
python3 -m src.models.make_track_dataset_h5 \
    ## Reference genome to extract DNA sequences
    --reference ${MM10REF} \
    ## Extracted sequences length in bp (must be multiple of width)
    --length 25600 \
    ## Sequence bin size [Enformer uses 128 bp]
    --width 128 \
    ## Minimal signal standard deviation, to drop blank examples
    ## Cutoff used in this example avoid dropping most sites
    --cutoff 0.00001 \
    ## Number of examples extracted per batches
    --size 25000 \
    ## Number of batches to try extract.
    --batches 20 \
    ## Random generator seed number to ensure reproducibility
    --seed 55 \
    ## This command will generate {prefix}.bed with all examples coordinates
    ## and a {prefix}.{0..batches}.h5 dataset for each batch
    --prefix out/mESC-CAGE/dataset \
    ## Coordinates to be considerate when sampling examples
    mappable.bed \
    ## Prediction track specifications
    tracks.tsv
```

### Train a new prediction head

Next, we use the generated dataset to train the new output head, while preserving the Enformer trunk weights, thus preserving its compatibility with original predictions.
The dataset batches are divided into training and validation datasets, based on the `validation` parameter.
During each training step, examples from the training dataset are presented in batches and the output layer weights are updated.
The training runs for a certain number of epochs, each consisting of a certain number of steps.

```bash
## Must define and export HEADSIZE as the number of tracks in the new head.
## It is required instantiate the new layer correctly
HEADSIZE=$(cat out/mESC-CAGE/tracks.tsv | wc -l)
export HEADSIZE

## Execute the following to train the new output head
python3 -m src.models.train_track_h5 \
    ## Number of batches created when constructing the training dataset.
    --dataset-batches 20 \
    ## Random generator seed number to ensure reproducibility
    --seed 55 \
    ## Training parameters
    --learning-rate 0.0005 \
    ## Number of training epochs to conduct
    --epochs 20 \
    ## Number of steps to be conducted on each epoch
    --steps 400 \
    ## Number of examples presented at each step.
    ## The total number of examples presented at each epoch is {steps} * {batch}
    --batch 4 \
    ## Ratio of dataset batches to be reserved for training validation
    --validation 0.1
    ## Where to save new head weights
    out/mESC-CAGE/model \
    ## Training dataset prefix
    out/mESC-CAGE/dataset \
```

### Predict new head results

Using the trained weights we can generate predictions for targeted regions using the command below.

```bash
## Generate predictions for the regions in `targets.bed` based on their
## genomic sequences from ${MM10REF}.
python3 -m src.models.predict_array \
    ## Enformer trunk model to use
    original 
    ## Bed file with targeted regions
    out/mESC-CAGE/targets.bed \
    ## Model output head to use, must be `human`, `mouse` or
    ## path for a new output head
    --head out/mESC-CAGE/model \
    ## Reference genome to extract DNA sequences
    --reference ${MM10REF} \
    ## Length of fragments when making predictions.
    ## It resizes the targeted regions to be a multiple of this length and
    ## breaks into non-overlapping fragments during prediction.
    --length 25600 \
    ## Number of 1-bp shifts centered at each fragment used to average the
    ## predicted signal
    --strides 10 \
    ## Numpy array where the signal are saved
    --output out/mESC-CAGE/targets.npy

## Define a number sequence for each track included in the new head
Tracks=$(echo $HEADSIZE | awk '{ print $1 - 1 }')
Tracks=$(seq -s' ' 0 $Tracks)

## Generate bedgraph tracks for each prediction track.
## It produces {prefix}.{0..track}.bedgraph for each track index in $Tracks
python3 -m src.data.array_to_bedgraph \
    out/mESC-CAGE/targets.bed \
    out/mESC-CAGE/targets.npy \
    --tracks $Tracks \
    --prefix out/mESC-CAGE/targets
```

---

## Fine-tuning Enformer on Synthetic Regulatory Genomics data

Code to fine-tuning on experimentally evaluated synthetic constructs delivered to a genomic locus.
We developed a fine-tuning strategy to improve performance by incorporating synthetic regulatory genomics datasets.
We added a new independent output layer that uses the baseline Enformer feature extraction trunk to predict our synthetic assays expression data.
The new output layer is composed of a self-attention layer to capture relevant features independently of position and a dense layer to combine the resulting signal into a single prediction value.
We evaluated three configurations of our new output self-attention layer: **SingleHead 64/64**, **SingleHead 64/128**, and **MultiHead 64/64**. **SingleHead 64/64** applies a single projection of 64 key and value matrices, **SingleHead 64/128** applies a single projection of 64 key and 128 value matrices, and **MultiHead 64/64** applies four independent projections of 64 key and value matrices.

### Building a fine-tuning dataset

First we prepare the environment and define some reference requirements.

```bash
## Define a seed to guarantee  reproducibility
SEED1=4200
SEED2=5200
MM10REF="/path/to/mm10.fasta"
Sox2LCR_COORDINATES="chr3:34732778-34772706"

## Create a dataset directory
mkdir -p data/dataset
```

Next we build a synthetic payload dataset by replacing the endogenous **Sox2 LCR** locus with our synthetic payloads replicating assays from [_Brosh et al. 2023_](https://www.cell.com/molecular-cell/fulltext/S1097-2765(23)00154-5).
By default, the generated sequences are 28,880bp long (25,600 target window +
1280 padding) centered at the replaced locus.

```bash
python3 -m src.data.make_dataset \
    ## Payload activity reference table. A TAB delimited file, requiring 
    ## `MenDel.Name`, `group`, `foldchange`, `activity` fields to be defined.
    data/raw/Brosh2023_TableS4.tsv \
    ## Payloads sequence fasta. Sequence names must correspond to `MenDel.Name`
    ## from activity table.
    data/raw/Brosh2023_Payload.fasta \
    ## BED Annotation of relevant locus from payloads.
    --annotation data/raw/Sox2_DistalLCR.bed \
    ## Reference genome which the payloads will be inserted.
    --reference ${MM10REF} \
    ## Genomic coordinates to be replaced by payloads.
    --context $Sox2LCR_COORDINATES \
    ## Seed to guarantee results reproducibility. It is relevant when assigning
    ## payloads to training or testing folds
    --seed ${SEED1} \
    ## folds assignment ratio. Due to payload diversity constraints we assigned
    ## everyone to a training fold and sampled two set of sequences to training
    ## and testing.
    --folds train=1 \
    --output data/dataset/Brosh2023.tsv
```

From this dataset, we sampled 48 random strides of 25,600bp for each payload to
be a training dataset and other 48 strides for testing.

```bash
## Will generate a sequence (`{prefix}-sequence.{fold}.npy`) and activity
## (`{prefix}-activity.{fold}.npy`) employing during training and testing.
python3 -m src.data.make_dataset_array data/dataset/Brosh2023.tsv \
    --prefix data/dataset/Brosh2023 \
    --seed $SEED1 \
    --stride 1 \
    --length 25600 \
    --sample 48

cat data/dataset/Brosh2023.tsv | sed 's/train/validation/' > tmp.tsv
python3 -m src.data.make_dataset_array tmp.tsv \
    --prefix data/dataset/Brosh2023 \
    --seed $SEED2 \
    --stride 1 \
    --length 25600 \
    --sample 48
```

### Fine-Tuning

After generating a training and validation dataset, we can fine tune Enformer
weights based on our dataset with the command below:

```bash
MODEL="out/SingleHead64_64" ## Path to model weights
## Fine-tuned attention layer specifications
MODELSPEC="--key-size 64  --value-size 64  --num-heads 1"

python3 -m src.models.train \
    ${MODEL} data/dataset/Brosh2023 ${MODELSPEC} \
    --learning-rate 1E-5 \
    ## Number of fine-tuning epochs
    --epochs 10 \
    ## Number of evaluations per epoch
    --steps 100 \
    ## Number of entries evaluated simultaneously
    --batch 4
```

### Evaluate results

After fine-tuning, we can use the following code to evaluate our model predictions with the original dataset results.
It outputs a table consisting of dataset info (payload, group, fold, and activity) together with evaluated sequence offset, directly predicted signal and sum of maximum values from all annotated sites (here Sox2 LCR DHSs).

```bash
python3 -m src.models.predict \
    ${MODEL} data/dataset/Brosh2023.tsv ${MODELSPEC} \
    ## Enformer output track of interest
    --head mouse --track 10 \
    ## Number of samples and strides to be evaluated per payload
    --sample 32 --stride 1 \
    --seed $SEED1 \
    --output out/SingleHead64_64/Brosh2023.tsv
```
