## Supplemental Code for "Iterative improvement of deep learning models using synthetic regulatory genomics": array_correlation.py

```
import os
import sys
import argparse

import tqdm
import numpy as np

from dotenv import load_dotenv

def main(reference_array:str, query_array:str, outfile:str) -> int:
    if not os.path.exists(reference_array):
        print(f"Unable to locate reference array at: `{reference_array}`", file=sys.stderr)
        return(1)
    if not os.path.exists(query_array):
        print(f"Unable to locate query array at: `{query_array}`", file=sys.stderr)
        return(1)

    refarray = np.load(reference_array)
    qryarray = np.load(query_array)
    
    outfile = open(outfile, "w") if outfile != "-" else sys.stdout

    if refarray.shape != qryarray.shape:
        print(f"Reference and query arrays must have the same dimension", file=sys.stderr)
        print(f"Reference: {refarray.shape}", file=sys.stderr)
        print(f"Query:     {qryarray.shape}", file=sys.stderr)
        return(1)
    
    nentry, _, ntrack = refarray.shape
    for i in tqdm.tqdm(range(nentry)):
        corr = []
        for j in range(ntrack):
            ref = refarray[i, :, j]
            qry = qryarray[i, :, j]
            r = np.corrcoef(ref, qry)[0, 1]
            corr.append(np.round(r, 5))
        print(i, *corr, sep="\t", file=outfile)
    outfile.close()
    return(0)


if __name__ == "__main__":
    load_dotenv()

    parser = argparse.ArgumentParser("analyze.peak")
    parser.add_argument('reference_array', type=str)
    parser.add_argument('query_array', type=str)
    parser.add_argument('--outfile', type=str, default="-")

    args = parser.parse_args()
    print(args, file=sys.stderr)
    exit(main(**vars(args)))
```
