## Supplemental Code for "Iterative improvement of deep learning models using synthetic regulatory genomics": array_correlation_bw.py

```
#! /usr/bin/env python3

import os
import sys
import csv
import argparse

import tqdm
import pysam
import pyBigWig
import h5py
import numpy as np

from dotenv import load_dotenv
from typing import Dict

from src.utils import dna_1hot

STAT = dict(
    sum  = lambda x: np.sum(x, axis=1, dtype=np.float32),
    mean = lambda x: np.mean(x, axis=1, dtype=np.float32)
)

class CovReader(object):
    def __init__(self, options:Dict):
        self._options = options
        self._covfile = pyBigWig.open(self._options["file"])
        if "clip" in self._options:
            self._clip = float(self._options["clip"])
        if "scale" in self._options:
            self._scale = float(self._options["scale"])
        self._func = STAT[self._options["sum_stat"]]
        
    def measure(self, chrom:str, start:int, end:int, width:int):
        cov = self._covfile.values(chrom, start, end, numpy=True)
        ## baseline to median
        baseline = np.percentile(cov, 50)
        baseline = np.nan_to_num(baseline)
        cov[np.isnan(cov)] = baseline

        cov = cov.reshape((end-start)//width, width)
        cov = self._func(cov)

        if self._scale is not None:
            cov = self._scale * cov
        if self._clip is not None:
            cov = np.clip(cov, -self._clip, self._clip)
        ## Clip to f16 limits
        cov = np.clip(cov, np.finfo(np.float16).min, np.finfo(np.float16).max)
        return(cov.astype(np.float16))

    def close(self):
        self._covfile.close()

    def __repr__(self):
        return(f"<CovReader {str(self._options)}>")


def pearson(true, pred):
    count = true.shape[0]
    true_mean = np.mean(true, 0)
    pred_mean = np.mean(pred, 0)
    true_var  = np.sum(true**2, 0) - count * (true_mean ** 2)
    pred_var  = np.sum(pred**2, 0) - count * (pred_mean ** 2)

    covariance = (
        np.sum(true * pred, 0)
        - true_mean * np.sum(pred, 0)
        - pred_mean * np.sum(true, 0)
        + (true_mean * pred_mean * count)
    )
    tp_var = np.sqrt(true_var) * np.sqrt(pred_var)
    return covariance / tp_var


def main(array:str, bedfile:str, tracks:str, outfile:str) -> int:
    ## Check files exists
    if not os.path.exists(tracks):
        print(f"Unable to locate file at`{tracks}`", file=sys.stderr)
        return(1)

    ## Setup
    tracks = csv.DictReader(open(tracks, "r"), delimiter="\t")
    tracks = [CovReader(line) for line in tracks]

    track_index = np.array([int(track._options["index"]) for track in tracks])

    coords = [line.rstrip().split("\t") for line in open(bedfile, 'r')]
    coords = [(f[0], int(f[1]), int(f[2])) for f in coords]

    pred_array = np.log10(np.load(array) + 1)

    outfilef = open(outfile, "w")

    for i in tqdm.tqdm(range(pred_array.shape[0])):
        chrom, start, stop = coords[i]
        
        true = [track.measure(chrom, start, stop, 128) for track in tracks]
        true = np.stack(true, axis=-1)
        true = np.log10(true + 1)

        pred = pred_array[i, ..., track_index].T
        pred = np.log10(pred + 1)

        corr = pearson(true, pred)
        print(chrom, start, stop, i, *corr, sep="\t", file=outfilef)
    
    outfilef.close()
    for track in tracks:
        track.close()
    
    return(0)


if __name__ == "__main__":
    load_dotenv()
    
    parser = argparse.ArgumentParser(prog="array-correlation-bw")
    parser.add_argument('array', type=str)
    parser.add_argument('bedfile', type=str)
    parser.add_argument('tracks', type=str)
    parser.add_argument('--outfile', type=str, default="-")

    args = parser.parse_args()

    print(args, file=sys.stderr)
    exit(main(**vars(args)))
```
