## Supplemental Code for "Iterative improvement of deep learning models using synthetic regulatory genomics": call_peak.py

```
import os
import sys
import argparse
import csv
from collections import defaultdict
from typing import List, Dict, Iterable, Callable

import numpy as np
import pyBigWig


def main(bigwig:str, min_score:float, min_length:int, target:str, outfile:str, **kwargs) -> int:
    if not os.path.exists(bigwig):
        print(f"Unable to locate {bigwig}", file=sys.stderr)
        return(1)
    
    bigwig = pyBigWig.open(bigwig)
    outfile = sys.stdout if outfile == "-" else open(outfile, "w")
    
    if (target is not None) and (os.path.exists(target)):
        intervals = csv.reader(open(target, 'r'), delimiter="\t")
        intervals = [(x[0], int(x[1]), int(x[2])) for x in intervals]
    else:
        intervals = [(c, ) for c in bigwig.chroms()]
    
    for it in intervals:
        last = (0, 0)
        bstart = -1
        for (start, end, value) in bigwig.intervals(*it):
            if (bstart == -1) and (value >= min_score):
                bstart = start
            elif ((bstart != -1) and (value < min_score)) or (last[1] != start):
                if (bstart != -1) and (last[1] - bstart >= min_length):
                    print(it[0], bstart, end, sep="\t", file=outfile)
                bstart = -1
            last = (start, end)

    outfile.close()
    return(0)

if __name__ == "__main__":
    parser = argparse.ArgumentParser("call_peak")
    parser.add_argument('bigwig', type=str)
    parser.add_argument('--target', type=str)
    parser.add_argument('--min-score', type=float, default=0.5)
    parser.add_argument('--min-length', type=int, default=128)
    parser.add_argument('--outfile', type=str, default="-")

    args = parser.parse_args()
    print(args, file=sys.stderr)
    exit(main(**vars(args)))
```
