## Supplemental Code for "Iterative improvement of deep learning models using synthetic regulatory genomics": dinuc_frequency.py

```
#! /usr/bin/env python3
import os
import sys
import argparse

import tqdm
import pysam
import numpy as np

from dotenv import load_dotenv


def main(coordinates:str, reference:str, output: str) -> int:
    ## Check required files
    if not os.path.isfile(coordinates):
        print(f"Unable to locate `{coordinates}`", file=sys.stderr)
        exit(1)
    if not os.path.isfile(reference):
        print(f"Unable to locate `{reference}`", file=sys.stderr)
        exit(1)
    
    ## SETUP
    print("! Setup", file=sys.stderr)
    coordinateName = os.path.splitext(os.path.basename(coordinates))[0]

    reference = pysam.FastaFile(args.reference)
    outfile = sys.stdout if output == "-" else open(output, "w")
    coordinates = sys.stdin if coordinates == "-" else open(coordinates, "r")

    coords = [line.rstrip().split("\t") for line in coordinates]
    coords = [
        (x[0], int(x[1]), int(x[2]), coordinateName if len(x) < 4 else x[3])
        for x in coords
    ]

    bases = dict(A=0, C=1, G=2, T=3, N=4)
    toBases = "ACGTN"
    dinuc_count = dict()

    for (chrom, start, end, name) in tqdm.tqdm(coords):
        if name not in dinuc_count:
            dinuc_count[name] = np.zeros((5, 5), dtype=np.int32)
        count = dinuc_count[name]

        seq = reference.fetch(chrom, start, end)
        for i in range(1, len(seq)):
            if seq[i] not in bases or seq[i-1] not in bases:
                print(f"Unexpected base found at {chrom}:{start}-{end}", file=sys.stderr)
                continue
            fromBase = bases[seq[i-1]]
            toBase = bases[seq[i]]
            count[fromBase, toBase] += 1

    for site in dinuc_count:
        count = dinuc_count[site]
        freq = count / np.sum(count)
        for i in range(count.shape[0]):
            for j in range(count.shape[1]):
                print(site, toBases[i] + toBases[j], count[i, j], freq[i, j], sep="\t", file=outfile)
    return(0)
    

if __name__ == "__main__":
    load_dotenv()
    parser = argparse.ArgumentParser(prog="dinuc-frequency")
    parser.add_argument('coordinates', type=str)
    parser.add_argument('--reference', type=str)
    ## model parameters
    parser.add_argument('--output', type=str, default="dinuc-freq.txt")
    
    args = parser.parse_args()
    print(args, file=sys.stderr)
    exit(main(**vars(args)))
```
