## Supplemental Code for "Iterative improvement of deep learning models using synthetic regulatory genomics": npy_to_h5.py

```
import os
import sys
import argparse
import pathlib

import h5py
import numpy as np
from dotenv import load_dotenv
from src.utils import pad_or_crop


def main(bedfile:str, array:str, output:str) -> int:
    if not os.path.exists(bedfile):
        print(f"Unable to locate '{bedfile}'", file=sys.stderr)
        return(1)
    
    if not os.path.exists(array):
        print(f"Unable to locate '{array}'", file=sys.stderr)
        return(1)
    
    if output is None:
        output = str(pathlib.Path(array).with_suffix(".h5"))
    
    print(f"Converting {array} into {output}", file=sys.stderr)

    outcome = np.load(array)

    targets = [line.rstrip().split("\t") for line in open(bedfile, "r")]
    targets = [
        (x[0], int(x[1]), int(x[2]))
        for x in targets if not x[0].startswith('#') and len(x) >= 3
    ]
    
    ## Estimate length based on 128bp binsize
    length = 128 * outcome.shape[1]
    ## Standardize the length
    targets = [(x[0], *pad_or_crop(x[1], x[2], length)) for x in targets]

    chroms = [chrom for chrom, _, _ in targets]
    chroms_label = list(set(chroms))
    chroms_index = np.array([chroms_label.index(chrom) for chrom in chroms], dtype="uint8")
    starts = np.array([start for _, start, _ in targets], dtype="uint32")
    ends = np.array([end for _, _, end in targets], dtype="uint32")

    ## Save as hdf5 file
    with h5py.File(output, 'w') as h5:
        h5.create_dataset('chrom_label', data=chroms_label, dtype=h5py.string_dtype(encoding='utf-8'))
        h5.create_dataset('chrom', data=chroms_index, dtype='uint8')
        h5.create_dataset('start', data=starts, dtype='uint32', compression='gzip')
        h5.create_dataset('end', data=ends, dtype='uint32', compression='gzip')
        h5.create_dataset('prediction', data=outcome, dtype='float16', compression='gzip')
    return(0)


if __name__ == "__main__":
    load_dotenv()
    parser = argparse.ArgumentParser("npy_to_h5")
    parser.add_argument("bedfile", type=str)
    parser.add_argument("array", type=str)
    parser.add_argument("--output", type=str)

    args = parser.parse_args()
    print(args, file=sys.stderr)
    exit(main(**vars(args)))
```
