## Supplemental Code for "Iterative improvement of deep learning models using synthetic regulatory genomics": payload_stat.py

```
import os
import sys
import argparse

from typing import List, Tuple

import numpy as np
import tqdm

from dotenv import load_dotenv

FUNCS=dict(
    max=lambda x, *args, **kwargs: np.max(x),
    mean=lambda x, *args, **kwargs: np.mean(x),
    median=lambda x, *args, **kwargs: np.median(x),
    propCutoff=lambda x, cutoff, *args, **kwargs: np.mean(x > cutoff))


def read_bed(bedfile):
    def parse_bed(fs):
        fs[1] = int(fs[1])
        fs[2] = int(fs[2])
        return fs
    chunks = [line.rstrip() for line in bedfile]
    chunks = [
        parse_bed(line.split("\t"))
        for line in chunks
        if (line != "") and (not line.startswith("#"))]
    return chunks


def read_bedgraph(bedgraph):
    chunks = [line.rstrip().split("\t") for line in bedgraph]
    if len(chunks[0]) <= 4:
        chrom = []
        start = []
        stop  = []
        score = []
        for fs in chunks:
            chrom.append(fs[0])
            start.append(int(fs[1]))
            stop.append(int(fs[2]))
            score.append(float(fs[3]))
        result = dict(chrom = np.array(chrom), start = np.array(start), stop = np.array(stop), score = np.array(score))
        result = dict(default=result)
    else:
        result = dict()
        for fs in chunks:
            if fs[3] not in result:
                result[fs[3]] = dict(chrom=[], start=[], stop=[], score=[])
            result[fs[3]]['chrom'].append(fs[0])
            result[fs[3]]['start'].append(int(fs[1]))
            result[fs[3]]['stop'].append(int(fs[2]))
            result[fs[3]]['score'].append(float(fs[4]))
        result = dict(
            (payload, dict((key, np.array(result[payload][key])) for key in result[payload]))
            for payload in result
        )
    return(result)


def main(bedfile:str, bedgraphfile:List[str], outfile:str, stats:str) -> int:    
    ## Checkup
    if bedfile != "-" and not os.path.exists(bedfile):
        print(f"~ Unable to locate bedfile at`{bedfile}`", file=sys.stderr)
        return(1)
    if not os.path.exists(bedgraphfile):
        print(f"~ Unable to locate bedgraph at `{bedgraphfile}`", file=sys.stderr)
        return(1)
    
    ## Setup
    funcs = [FUNCS[stat] for stat in stats]
    
    targets  = read_bed(open(bedfile, 'r'))
    bedgraph = read_bedgraph(open(bedgraphfile, 'r'))

    bedfile = sys.stdin if bedfile == "-" else open(bedfile, "r")
    outfile = sys.stdout if outfile == "-" else open(outfile, "w")

    labels = [[payload] * len(funcs) for payload in bedgraph]
    labels = sum(labels, [])
    print("chrom", "start", "stop", "name", *labels, sep="\t", file=outfile)

    for chunk in tqdm.tqdm(targets):
        chrom, start, stop, name = chunk[:4]

        values = []
        for payload in bedgraph:
            ## Subset target region
            target = bedgraph[payload]['chrom'] == chrom
            target = target & (bedgraph[payload]['stop'] > start)
            target = target & (bedgraph[payload]['start'] < stop)
            score = bedgraph[payload]['score'][target]
            
            if len(score) > 0:
                values += [round(fn(score), 5) for fn in funcs]
            else:
                values += ['NA'] * len(funcs)
        print(chrom, start, stop, name, *values, sep="\t", file=outfile)
    return(0)

if __name__ == "__main__":
    load_dotenv()

    parser = argparse.ArgumentParser("analyze.payload_stat")
    parser.add_argument("bedfile", type=str, default="-")
    parser.add_argument("bedgraphfile", type=str)
    parser.add_argument('--outfile', type=str, default="-")
    parser.add_argument("--stats", type=str, nargs='+',
                        default=['mean'],
                        choices=FUNCS.keys())

    args = parser.parse_args()
    print(args, file=sys.stderr)
    exit(main(**vars(args)))
```
