## Supplemental Code for "Iterative improvement of deep learning models using synthetic regulatory genomics": peak.py

```
import os
import sys
import argparse
from collections import defaultdict
from typing import List, Dict, Iterable, Callable

import numpy as np


def is_number(element:str) -> bool:
    partition=element.partition('.')
    if ((partition[0].isdigit() and partition[1]=='.' and partition[2].isdigit()) or
        (partition[0]=='' and partition[1]=='.' and partition[2].isdigit()) or
        (partition[0].isdigit() and partition[1]=='.' and partition[2]=='') or
        (partition[0].isdigit() and partition[1]=='' and partition[2]=='')):
        return True
    else:
        return False


def read_score(score:Iterable[str]) -> Dict[str, np.ndarray]:
    data = defaultdict(list)

    header = next(score)
    header = header.rstrip().split("\t")

    for entry in score:
        entry = entry.rstrip().split("\t")
        for i in range(len(header)):
            value = entry[i] if i in [0, 3] else float(entry[i])
            data[header[i]].append(value)
    
    for i in range(len(header)):
        data[header[i]] = np.array(data[header[i]])

    return(data)


def ecdf(x: np.ndarray) -> Callable[[np.array], np.array]:
    ## See https://stackoverflow.com/questions/15792552/numpy-scipy-equivalent-of-r-ecdfxx-function
    xs = np.sort(x)
    n = xs.shape[0]
    def _ecdf(v: np.ndarray) -> np.array:
        return np.searchsorted(xs, v, side='right') / n
    return _ecdf


def main(score:str, prefix:str, length:int, top:int) -> int:
    if not os.path.exists(score):
        print(f"~ Unable to locate score table at `{score}`", file=sys.stderr)
        return(1)

    data = read_score(open(score, "r"))

    tracks = [key for key in data.keys() if key not in ['chrom', 'start', 'end', 'name']]
    reference = tracks[0]

    ## CDF
    minval = min(np.min(data[x]) for x in tracks)
    maxval = max(np.max(data[x]) for x in tracks)
    valspace = np.linspace(minval, maxval, 1000)
    # valspace = np.linspace(np.log10(minval + 1), np.log10(maxval + 1), 1000)
    # valspace = (10 ** valspace) - 1

    track_ecdf = dict((track, ecdf(data[track])(valspace)) for track in tracks)

    with open(f"{prefix}-ecdf.txt", "w") as outf:
        print("track", "value", "cdf", sep="\t", file=outf)
        for track in tracks:
            for i in range(len(valspace)):        
                print(track, valspace[i], track_ecdf[track][i], sep="\t", file=outf)

    ## Chromossome-wise Pearson r
    chroms = np.unique(data["chrom"])
    track_rcoef = list()
    for chrom in chroms:
        chromvec = data["chrom"] == chrom
        refvalue = data[reference][chromvec]
        for track in tracks:
            value = data[track][chromvec]
            if np.std(value) == 0 or np.std(refvalue) == 0:
                rcoef = np.nan
            else:
                rcoef = np.corrcoef(value, refvalue)[0,1]
            track_rcoef.append((track, chrom, value.shape[0], rcoef))
    
    with open(f"{prefix}-pearsonr.txt", "w") as outf:
        print("track", "chrom", "n", "rcoef", sep="\t", file=outf)
        for entry in track_rcoef:
            print(*entry, sep="\t", file=outf)

    ## Cutoff call
    quantile_call = lambda x, q: x > np.quantile(x, q)

    cutoffs = [0.5, 0.7, 0.8, 0.9]
    track_call = dict(
        (cutoff, dict((track, quantile_call(data[track], cutoff)) for track in tracks))
        for cutoff in cutoffs
    )
    track_perf = []

    for cutoff in track_call:
        trcall = track_call[cutoff]
        for chrom in chroms:
            chromvec = data["chrom"] == chrom
            refcall = trcall[reference][chromvec]
            for track in tracks:
                call = trcall[track][chromvec]

                accuracy = np.mean(refcall == call)
                true_positive = np.mean(call[refcall])
                true_negative = np.mean(np.logical_not(call)[np.logical_not(refcall)])

                track_perf.append((track, chrom, cutoff, len(value), accuracy, true_positive, true_negative))
    
    with open(f"{prefix}-accuracy.txt", "w") as outf:
        print("track", "chrom", "cutoff", "n", "accuracy", "true_positive", "true_negative", sep="\t", file=outf)
        for entry in track_perf:
            print(*entry, sep="\t", file=outf)

    ## Report highly diverging regions
    trcall = track_call[0.8]
    region_report = []
    for chrom in chroms:
        chromvec = data["chrom"] == chrom
        startvec = data["start"][chromvec]
        endvec = data["end"][chromvec]
        position = (startvec + endvec) / 2.0

        refcall  = trcall[reference][chromvec]
        regions = np.cumsum(np.diff(np.concatenate(([0], position))) > length)
        
        for region in np.unique(regions):
            regionvec = (regions == region)
            start = np.min(startvec[regionvec])
            end = np.max(endvec[regionvec])
            num = np.sum(regionvec)
            acc = [
                np.mean(refcall[regionvec] == trcall[track][chromvec][regionvec])
                for track in tracks]
            region_report.append((chrom, int(start), int(end), num, *acc))

    with open(f"{prefix}-regions.txt", "w") as outf:
        print("chrom", "start", "end", "n", *tracks, sep="\t", file=outf)
        for entry in region_report:
            print(*entry, sep="\t", file=outf)

    return(0)

if __name__ == "__main__":
    default_length = os.getenv("SEQ_LENGTH", 25600)

    parser = argparse.ArgumentParser("analyze.peak")
    parser.add_argument('score', type=str)
    parser.add_argument('--prefix', type=str, default="peak")
    parser.add_argument('--length', type=int, default=default_length)
    parser.add_argument('--top', type=int, default=20)

    args = parser.parse_args()
    print(args, file=sys.stderr)
    exit(main(**vars(args)))
```
