## Supplemental Code for "Iterative improvement of deep learning models using synthetic regulatory genomics": stat.py

```
import sys
import argparse

# import pyBigWig
import numpy as np

from dotenv import load_dotenv

FUNCS = dict(
    max = lambda x, *args, **kwargs: np.max(x),
    mean = lambda x, *args, **kwargs: np.mean(x),
    median = lambda x, *args, **kwargs: np.median(x),
    quantile = lambda x, q, *args, **kwargs: np.quantile(x, q)
)

def main(stat:str, quantile:float) -> int:
    statfn = FUNCS[stat]

    data = [float(line.rstrip()) for line in sys.stdin]
    data = np.array(data)

    print(statfn(data, quantile), file=sys.stdout)
    
    return(0)

if __name__ == "__main__":
    load_dotenv()

    parser = argparse.ArgumentParser("analyze.track")
    parser.add_argument("stat", type=str, default="mean", choices=FUNCS.keys())
    parser.add_argument("--quantile", type=float, default=None)

    args = parser.parse_args()
    print(args, file=sys.stderr)
    exit(main(**vars(args)))
```
