## Supplemental Code for "Iterative improvement of deep learning models using synthetic regulatory genomics": track.py

```
import os
import sys
import argparse
import subprocess

from typing import List, Tuple

# import pyBigWig
import numpy as np
import tqdm

from dotenv import load_dotenv

FUNCS=dict(
    max=lambda x, *args, **kwargs: np.max(x),
    mean=lambda x, *args, **kwargs: np.mean(x),
    median=lambda x, *args, **kwargs: np.median(x),
    propCutoff=lambda x, cutoff, *args, **kwargs: np.mean(x > cutoff))


class BigWig:
    def __init__(self, file:str) -> None:
        try:
            import pyBigWig
            self._bigwig = True
            self._file = pyBigWig.open(file)
        except ImportError:
            self._bigwig = False
            self._file = file
    
    def _fetch(self, chrom:str, start:int, end:int):
        ps = subprocess.run(
            ['bigWigToBedGraph', self._file, '/dev/stdout',
             f"-chrom={chrom}", f"-start={start}", f"-end={end}"],
            capture_output=True, text=True)
        return [parse_bedgraph(chunk)
                for chunk in ps.stdout.strip().split("\n")
                if chunk != ""]
    
    def fetch(self, chrom:str, start:int, end:int) -> np.ndarray:
        if self._bigwig:
            result = self._file.values(chrom, start, end, numpy=True)
            return np.nan_to_num(result)
        
        result = np.zeros(end-start)
        for _, cstart, cend, value in self._fetch(chrom, start, end):
            rstart = max(0, cstart-start)
            rend = min(end-start, cend-start)
            result[rstart:rend] = value
        return result


def parse_bedgraph(entry:str) -> Tuple[str, int, int, float]:
    entry = entry.rstrip().split("\t")
    return(entry[0], int(entry[1]), int(entry[2]), float(entry[3]))


def read_bed(bedfile):
    def parse_bed(fs):
        fs[1] = int(fs[1])
        fs[2] = int(fs[2])
        return fs
    chunks = [line.rstrip() for line in bedfile]
    chunks = [
        parse_bed(line.split("\t"))
        for line in chunks
        if (line != "") and (not line.startswith("#"))]
    return chunks


def main(bedfile:str, wigfiles:List[str], outfile:str, stats:str) -> int:    
    ## Checkup
    if bedfile != "-" and not os.path.exists(bedfile):
        print(f"~ Unable to locate bedfile at`{bedfile}`", file=sys.stderr)
        return(1)
    if any(not os.path.exists(wig) for wig in wigfiles):
        print(f"~ Unable to locate all wigfiles", file=sys.stderr)
        return(1)
    
    ## Setup
    funcs = [FUNCS[stat] for stat in stats]
    wigs  = [BigWig(file) for file in wigfiles]
    # wigs = [avgOverBed(file, bedfile) for file in wigfiles]

    bedfile = sys.stdin if bedfile == "-" else open(bedfile, "r")
    outfile = sys.stdout if outfile == "-" else open(outfile, "w")

    targets = read_bed(bedfile)
    for chunk in tqdm.tqdm(targets):
        chrom, start, end = chunk[:3]
        try:
            values = [wig.fetch(chrom, start, end) for wig in wigs]
            stats = [round(fn(value), 5) for value in values for fn in funcs]
            print(*chunk, *stats, sep="\t", file=outfile)
        except Exception as e:
            print("FAIL", e, *chunk, file=sys.stderr)
            return(1)
    return(0)

if __name__ == "__main__":
    load_dotenv()

    parser = argparse.ArgumentParser("analyze.track")
    parser.add_argument("bedfile", type=str, default="-")
    parser.add_argument("wigfiles", type=str, nargs='+')
    parser.add_argument('--outfile', type=str, default="-")
    parser.add_argument("--stats", type=str, nargs='+',
                        default=['mean'],
                        choices=FUNCS.keys())

    args = parser.parse_args()
    print(args, file=sys.stderr)
    exit(main(**vars(args)))
```
