## Supplemental Code for "Iterative improvement of deep learning models using synthetic regulatory genomics": distance.py

```
#! /usr/bin/env python3
import os
import sys
import argparse

import tqdm
import h5py
import pysam
import numpy as np
import tensorflow as tf

from dotenv import load_dotenv
from src.utils import pad_or_crop, dna_1hot
from src.models import load_model, load_model_from_modeldef, head_func


bases = dict(A=0, C=1, G=2, T=3, N=4)
toOneHot = dna_1hot("ACGTN")


def shuffle_dinuc(seq:np.array, n:int = 1, seed:int = None) -> np.array:
    rng = np.random.RandomState(seed)

    chars, tokens = np.unique(seq, axis=0, return_inverse=True)
    shuf_next_inds = [ np.where(tokens[:-1] == t)[0] + 1 for t in range(len(chars)) ]

    results = np.empty((n, seq.shape[0], seq.shape[1]), dtype=seq.dtype)
    for i in range(n):
        ## shuffle indices
        for t in range(len(chars)):
            inds = np.arange(len(shuf_next_inds[t]))
            inds[:-1] = rng.permutation(len(inds)-1)
            shuf_next_inds[t] = shuf_next_inds[t][inds]
        
        ## build result
        counters = [0] * len(chars)
        ind = 0
        
        results[i][0] = chars[tokens[ind]]
        for j in range(1, len(tokens)):
            t = tokens[ind]
            ind = shuf_next_inds[t][counters[t]]
            counters[t] += 1
            results[i][j] = chars[tokens[ind]]
    return(results)


def random_dinuc(freq:np.array, length:int, n:int = 1, seed:int = None) -> np.array:
    rng = np.random.RandomState(seed)

    results = np.empty((n, length, 4), dtype=toOneHot.dtype)
    for i in range(n):
        cur = 0
        results[i, 0, :] = toOneHot[cur, :]
        for j in range(1, length):
            cur = rng.choice(np.arange(5), p=freq[cur, :])
            results[i, j, :] = toOneHot[cur, :]
    return(results)


def parse_coord(coord:str) -> tuple:
    name, location = coord.split(":")
    start, end = location.split("-")
    return(name, int(start), int(end))


def main(model:str, sites:str, length:int, step_size:int, seed:int, reference:str,
         head: str, shuffles:int, output: str, report:str, dinuc_frequency:str,
         key_size:int, value_size:int, num_heads:int) -> int:
    ## Check required files
    if not os.path.isfile(sites):
        print(f"Unable to locate `{sites}`", file=sys.stderr)
        exit(1)
    if not os.path.isfile(reference):
        print(f"Unable to locate `{reference}`", file=sys.stderr)
        exit(1)

    ## SETUP
    print("! Setup", file=sys.stderr)
    reference = pysam.FastaFile(args.reference)
    reportfile = sys.stdout if report == "-" else open(report, "w")

    dinuc_freq = None
    if dinuc_frequency and os.path.isfile(dinuc_frequency):
        dinuc_count = [line.strip().split("\t") for line in open(dinuc_frequency, 'r')]
        dinuc_count = [(x[1], int(x[2])) for x in dinuc_count]
        
        dinuc_freq = np.ones((5, 5), dtype=np.int32)
        for dinuc, count in dinuc_count:
            dinuc_freq[bases[dinuc[0]], bases[dinuc[1]]] += count
        dinuc_freq = dinuc_freq / np.sum(dinuc_freq, axis=0)[:, np.newaxis]
        print("Dinucleotide Transition Frequency [ACGTN -> ACGTN]", file=sys.stderr)
        print(np.round(dinuc_freq, 2), file=sys.stderr)

    ## Build virtual sequences
    sitedat = [line.strip().split("\t") for line in open(sites, 'r')]
    sitedat = [
        (x[0], int(x[1]), int(x[2]), parse_coord(x[3]), parse_coord(x[4]))
        for x in sitedat
    ]
    
    print("! Load model", file=sys.stderr)
    mdict = load_model_from_modeldef(model)
    model = head_func(mdict, head) ## Accept a custom head
    
    print("! Predicting effect", file=sys.stderr)
    index = 0
    results = dict(index=[], distance=[], prediction=[])

    for chrom, start, end, fixed, mobile in tqdm.tqdm(sitedat):
        pstart, pend = pad_or_crop(start, end, length)
        offset = pstart - start

        ## Extract and encode sequence
        seq = reference.fetch(chrom, pstart, pend)
        seq = dna_1hot(seq)
        if dinuc_freq is not None:
            shufseq = random_dinuc(dinuc_freq, length, shuffles, seed)
        else:
            shufseq = shuffle_dinuc(seq, shuffles, seed)

        ## Extract fixed and mobile tiles
        fixedStart = (fixed[1] + offset)
        fixedEnd = (fixed[2] + offset)
        fixedSeq = seq[fixedStart:fixedEnd, :]

        mobileStart = (mobile[1] + offset)
        mobileEnd = (mobile[2] + offset)
        mobileSeq = seq[mobileStart:mobileEnd, :]
        mobileLen = mobileSeq.shape[0]

        ## Restore fixed site
        shufseq[:, fixedStart:fixedEnd, :] = fixedSeq

        ## Build steps
        steps = np.concatenate([
            np.arange(fixedStart % step_size, fixedStart, step_size),
            np.arange(fixedEnd, length, step_size)])
        
        for stepStart in tqdm.tqdm(steps, leave=False):
            stepEnd = stepStart + mobileLen
            distance = stepStart-fixedStart
            if stepEnd > length:
                continue
            stepSeq = shufseq.copy()
            stepSeq[:, stepStart:stepEnd, :] = mobileSeq

            pred = model(stepSeq, is_training=False)
            pred = tf.math.reduce_mean(pred, axis=0) ## Average across shuffles
            pred = tf.cast(pred, dtype=tf.float16)
            
            ## Select central 256bp
            fixedPredStart = int(np.floor(fixedStart/128))
            fixedPredEnd = int(np.ceil(fixedEnd/128))
            centralPoint = (fixedPredStart + fixedPredEnd) // 2
            fixedAvg = tf.math.reduce_mean(pred[(centralPoint-1):(centralPoint+1), :], 0)
            # fixedAvg = [f"{x:0.3f}" for x in fixedAvg]
            fixedAvg = np.round(fixedAvg, 3)
            print(chrom, start, end, distance, mobile[1], mobile[2], fixed[0], *fixedAvg, sep="\t", file=reportfile)

            mobilePredStart = int(np.floor(stepStart/128))
            mobilePredEnd = int(np.ceil(stepEnd/128))
            mobileMax = tf.math.reduce_max(pred[mobilePredStart:mobilePredEnd, :], 0)
            # mobileMax = [f"{x:0.3f}" for x in mobileMax]
            mobileMax = np.round(mobileMax, 3)
            print(chrom, start, end, distance, mobile[1], mobile[2], mobile[0], *mobileMax, sep="\t", file=reportfile)
        
            results["index"].append(index)
            results["distance"].append(distance)
            results["prediction"].append(pred)
        index += 1

    print("! Saving predictions", file=sys.stderr)
    ## Save as hdf5 file
    with h5py.File(output, 'w') as h5:
        indices = np.array(results["index"], dtype="uint8")
        distances = np.array(results["distance"], dtype="int32")
        predictions = np.array(results["prediction"], dtype="float16")

        h5.create_dataset('indices', data=indices, dtype='uint8', compression='gzip')
        h5.create_dataset('distances', data=distances, dtype='int32', compression='gzip')
        h5.create_dataset('predictions', data=predictions, dtype='float16', compression='gzip')

    if report != "-":
        reportfile.close()
    return(0)
    

if __name__ == "__main__":
    load_dotenv()
    parser = argparse.ArgumentParser(prog="distance-assay")
    parser.add_argument('model', type=str)
    parser.add_argument('sites', type=str)
    parser.add_argument('--length', type=int, default=os.getenv('SEQ_LENGTH', 25600))
    parser.add_argument('--step-size', type=int, default=10000)
    parser.add_argument('--seed', type=int, default=42)
    parser.add_argument('--reference', type=str)
    parser.add_argument('--dinuc-frequency', type=str)
    ## model parameters
    modeldef = parser.add_argument_group('model definition')
    modeldef.add_argument('--key-size', type=int, default=64)
    modeldef.add_argument('--value-size', type=int, default=64)
    modeldef.add_argument('--num-heads', type=int, default=1)
    ## output parameters
    outputdef = parser.add_argument_group('output definition')
    outputdef.add_argument('--head', type=str, default='mouse', choices=['mouse', 'human'])
    outputdef.add_argument('--shuffles', type=int, default=10)
    outputdef.add_argument('--output', type=str, default="array.h5")
    outputdef.add_argument('--report', type=str, default="-")

    args = parser.parse_args()
    print(args, file=sys.stderr)
    exit(main(**vars(args)))
```
