## Supplemental Code for "Iterative improvement of deep learning models using synthetic regulatory genomics": array_to_bed.py

```
import os
import sys
import argparse
from typing import List

import tqdm
import numpy as np

from src.utils import pad_or_crop
from dotenv import load_dotenv

def main(
        array: str, bedfile: str, prefix: str, tracks: List[int],
        step:int, length:int) -> int:
    if not os.path.exists(array):
        print(f"Unable to locate prediction array at `{array}`", file=sys.stderr)
        return(1)

    if not os.path.exists(bedfile):
        print(f"Unable to locate bedfile at `{bedfile}`", file=sys.stderr)
        return(1)

    array = np.load(array)
    outfiles = dict((track, open(f"{prefix}.{track}.bed", "w"))
                    for track in tracks)
    
    targets = [line.rstrip().split("\t") for line in open(bedfile, 'r')]
    targets = [(x[0], int(x[1]), int(x[2]), x[3])
               for x in targets
               if not x[0].startswith('#') and len(x) >= 3]
    targets = [(x[0], *pad_or_crop(x[1], x[2], length), x[3])
               for x in targets]
    
    chroms = [x[0] for x in targets for _ in range(array.shape[1])]
    starts = [x[1] + shift * step for x in targets for shift in range(array.shape[1])]
    starts = np.array(starts, dtype=np.int32)

    for track in tracks:
        print(f"! Parsing track {track}", file=sys.stderr)
        outfile = outfiles[track]
        values = (array[:, :, track]).flatten()
        for i in tqdm.tqdm(range(values.shape[0])):
            print(chroms[i], starts[i], starts[i] + step, ".", values[i],
                  sep="\t", file=outfile)
        outfile.close()
    return(0)    


if __name__ == "__main__":
    load_dotenv()

    parser = argparse.ArgumentParser("array_to_bed")
    parser.add_argument('--prefix', type=str, default="track")
    parser.add_argument('--tracks', type=int, nargs='+', default=[10])
    parser.add_argument('--step', type=int, default=os.getenv("SEQ_WINDOW", 128))
    parser.add_argument('--length', type=int, default=os.getenv("SEQ_LENGTH", 256000))
    parser.add_argument('bedfile')
    parser.add_argument('array')

    args = parser.parse_args()
    print(args, file=sys.stderr)
    exit(main(**vars(args)))
```
