## Supplemental Code for "Iterative improvement of deep learning models using synthetic regulatory genomics": array_to_bedgraph.py

```
import os
import sys
import argparse
from typing import List, Dict, Tuple
from collections import defaultdict

import tqdm
import numpy as np
from intervaltree import IntervalTree
from dotenv import load_dotenv


def build_coordinates(
        targets: List[Tuple[str, int, int]],
        step: int, num_steps: int) -> Dict[str, np.ndarray]:
    coords = defaultdict(IntervalTree)
    for (chrom, start, _) in targets:
        for i in range(num_steps):
            coords[chrom].addi(start + i * step, start + (i+1) * step)
    
    post = dict()
    for chrom in coords:
        itree = coords[chrom]
        itree.split_overlaps()

        start = np.array([itv.begin for itv in itree])
        end = np.array([itv.end for itv in itree])
        post[chrom] = np.sort(np.vstack((start, end)))
    return post


def main(
        array: str, bedfile: str, prefix: str,
        tracks: List[int], step:int) -> int:
    if not os.path.exists(array):
        print(f"Unable to locate prediction array at `{array}`", file=sys.stderr)
        return(1)

    if not os.path.exists(bedfile):
        print(f"Unable to locate bedfile at `{bedfile}`", file=sys.stderr)
        return(1)
    
    array = np.load(array)
    outfiles = dict((track, open(f"{prefix}.{track}.bedgraph", "w"))
                    for track in tracks)
    
    targets = [line.rstrip().split("\t") for line in open(bedfile, 'r')]
    targets = [(x[0], int(x[1]), int(x[2])) for x in targets if not x[0].startswith('#') and len(x) >= 3]

    num_steps = array.shape[1]
    chroms = np.array([x[0] for x in targets for _ in range(num_steps)])
    starts = np.array([x[1] + i * step for x in targets for i in range(num_steps)])

    coords = build_coordinates(targets, step, num_steps)
    print([(chrom, coords[chrom].shape) for chrom in coords])

    for track in tracks:
        print(f"! Parsing track {track}", file=sys.stderr)
        outfile = outfiles[track]
        values = (array[:, :, track]).flatten()

        coord_acc = dict((chrom, np.zeros(coords[chrom].shape[1])) for chrom in coords)
        coord_n = dict((chrom, np.zeros(coords[chrom].shape[1])) for chrom in coords)

        for i in tqdm.tqdm(range(values.shape[0])):
            cchrom = chroms[i]
            cstart = starts[i]

            to_update = coords[cchrom][0, :] >= cstart
            to_update = np.logical_and(to_update, coords[cchrom][1, :] <= cstart + step)

            coord_acc[cchrom][to_update] += values[i]
            coord_n[cchrom][to_update] += 1
        for cchrom in coords:
            avg = coord_acc[cchrom] / coord_n[cchrom]
            for i in range(avg.shape[0]):
                print(cchrom, coords[cchrom][0, i], coords[cchrom][1, i],
                      f"{avg[i]:.5f}", sep="\t", file=outfile)
        outfile.close()
    return(0)    


if __name__ == "__main__":
    load_dotenv()

    parser = argparse.ArgumentParser("array_to_bed")
    parser.add_argument('--prefix', type=str, default="track")
    parser.add_argument('--tracks', type=int, nargs='+', default=[10])
    parser.add_argument('--step', type=int, default=os.getenv("SEQ_WINDOW", 128))
    parser.add_argument('bedfile')
    parser.add_argument('array')

    args = parser.parse_args()
    print(args, file=sys.stderr)
    exit(main(**vars(args)))
```
