## Supplemental Code for "Iterative improvement of deep learning models using synthetic regulatory genomics": make_dataset.py

```
#! /usr/bin/env python3

import os
import sys
import csv
import argparse

import tqdm
import pysam
import numpy as np

from dotenv import load_dotenv
from collections import defaultdict
from typing import Union, List, Dict, Tuple

from src.utils import parse_annotation, fold_split


def read_dataset(
        file:str, payload:str,
        annotations:List[Tuple[str, int, int, str]]) -> Dict[str, dict]:
    payload = pysam.Fastafile(payload)
    
    dataset = defaultdict(lambda: dict(activity = [], foldchange = []))
    reader = csv.DictReader(open(file, "r"), delimiter="\t")
    for entry in reader:
        name = entry["MenDel.Name"]
        if entry["foldchange"] == "NA" or entry["activity"] == "NA":
            continue
        dataset[name]["group"] = entry["groups"]
        dataset[name]["foldchange"].append(float(entry["foldchange"]))
        dataset[name]["activity"].append(float(entry["activity"]))

        ## Add sequence
        safename = name.replace("~", "_")
        sequence = payload.fetch(safename).upper()
        annotation = [c for c in annotations if c[0] == safename]
        dataset[name]["sequence"] = sequence
        dataset[name]["annotation"] = annotation
    payload.close()
    return dataset


def pad_or_trim_sequence(
        dataset:Dict[str, dict], length:int,
        context_left:str, context_right:str) -> Dict[str, dict]:
    for name in dataset:
        payload_sequence = dataset[name]["sequence"]
        payload_length = len(payload_sequence)
        if payload_length <= length:
            padding = (length - payload_length)
            leftpad = padding // 2
            dataset[name]["sequence"] = (
                context_left[-leftpad:] +
                payload_sequence +
                context_right[:(padding-leftpad)])
            dataset[name]["annotation"] = [
                (c[0], c[1] + leftpad, c[2] + leftpad, c[3])
                for c in dataset[name]["annotation"]]
        else:
            trimming = (payload_length - length)
            lefttrim = trimming // 2
            dataset[name]["sequence"] = payload_sequence[lefttrim:(lefttrim+length)]
            dataset[name]["annotation"] = [
                (c[0], c[1] - lefttrim, c[2] - lefttrim, c[3])
                for c in dataset[name]["annotation"]]
            dropout = [c[3] for c in dataset[name]["annotation"]
                        if not (c[1] >= 0 and c[2] <= length)]
            dropout = ", ".join(dropout)
            dataset[name]["annotation"] = [
                c for c in dataset[name]["annotation"]
                if c[1] >= 0 and c[2] <= length]
            print(f"! Cropping sequnce for {name}", file=sys.stderr)
            print(f"! - Dropout sites: {dropout}", file=sys.stderr)
    return dataset


def main(
        dataset:str, sequences:str, annotation:str,
        reference:str, context:str, length:int, seed:int,
        folds:Dict[str, float], output:str) -> int:
    ## Check files exists
    for file in [dataset, sequences, reference]:
        if not os.path.exists(file):
            print(f"Unable to locate required file at`{dataset}`", file=sys.stderr)
            return(1)
    ## Setup
    np.random.seed(seed)

    ## Build context sequence
    context = parse_annotation(context)
    reference = pysam.Fastafile(reference)
    context_left = reference.fetch(context[0], context[1] - length // 2, context[1]).lower()
    context_right = reference.fetch(context[0], context[2], context[2] + length // 2).lower()
    reference.close()

    annotations = []
    if annotation is not None and os.path.exists(annotation):
        annotations = [line.rstrip().split("\t") for line in open(annotation, "r")]
        annotations = [(x[0], int(x[1]), int(x[2]), x[3]) for x in annotations]
    
    dataset = read_dataset(dataset, sequences, annotations)
    dataset = pad_or_trim_sequence(dataset, length, context_left, context_right)
    dataset_keys = list(dataset.keys())

    ## Assign folds
    fold_name = list(folds.keys())
    fold_ratio = list([ folds[f] for f in fold_name ])
    fold = fold_split(fold_ratio, len(dataset_keys))

    ## Print out results
    outfile = open(output, 'w')
    print(
        "index", "name", "group", "fold", "activity", "annotation", "sequence",
        sep="\t", file=outfile)
    for i, name in tqdm.tqdm(enumerate(dataset_keys)):
        activity = np.array(dataset[name]["activity"])
        activity = np.mean(activity)
        annotation = [f"{c[3]}:{c[1]}-{c[2]}" for c in dataset[name]["annotation"]]
        annotation = ";".join(annotation)
        print(i, name, dataset[name]["group"], fold_name[fold[i]],
              f"{activity:.3f}", annotation,  dataset[name]["sequence"],
              sep="\t", file=outfile)
    return(0)


class KwargsParser(argparse.Action):
    def __call__(
            self, parser: argparse.ArgumentParser,
            namespace: argparse.Namespace,
            values: List[str],
            option_string: Union[str, None] = None) -> None:
        setattr(namespace, self.dest, dict())
        for value in values:
            key, value = value.split('=')
            getattr(namespace, self.dest)[key] = float(value)


if __name__ == "__main__":
    load_dotenv()

    default_length = int(os.getenv("SEQ_LENGTH", 25600)) + 1280

    parser = argparse.ArgumentParser(
        prog="src.data.make_dataset",
        help=(
            "Builds a dataset reference table from payloads sequences and " +
            "their activity/foldchange signal. " +
            "Generates a synthetic sequence for each payload by replacing an " +
            "endogenous locus with the payload sequence."
        ))
    parser.add_argument('dataset', type=str, help="path to a payload activity table.")
    parser.add_argument('sequences', type=str, help="path to payloads fasta.")
    parser.add_argument('--annotation', type=str, help="BED file of relevant payload sites to annotate.")
    parser.add_argument('--reference', type=str, help="path to reference genome fasta where the payloads will be inserted.")
    parser.add_argument("--context", type=str, help="genomic coordinates that the payload will replace.")
    parser.add_argument('--length', type=int, default=default_length,
        help=(
            "length of the synthetic sequence generated. Sequences will be generated centered at the {context} site. " +
            "We recommend include some padding to your target sequence size to allow random sample generation."
        ))
    parser.add_argument('--seed', type=int, default=5, help="random number generator seed")
    ## output parameters
    outputdef = parser.add_argument_group('output definition')
    outputdef.add_argument(
        '--folds', nargs='+', action=KwargsParser,
        default=dict(train=8, test=1, validation=1),
        help="payload fold assignment ratios (e.g. train=8 test=1 validation=1).")
    outputdef.add_argument('--output', type=str, default="dataset.tsv",
        help="path to output file.")
    args = parser.parse_args()

    print(args, file=sys.stderr)
    exit(main(**vars(args)))
```
