## Supplemental Code for "Iterative improvement of deep learning models using synthetic regulatory genomics": make_dataset_array.py

```
#! /usr/bin/env python3

import os
import sys
import argparse

import numpy as np
from dotenv import load_dotenv
from collections import defaultdict
from src.utils import read_dataset, generate_offsets, dna_1hot


def main(dataset:str, prefix: str, length: int, stride: int, sample: float, seed:int) -> int:
    ## Check files exists
    if not os.path.exists(dataset):
        print(f"Unable to locate required file at`{dataset}`", file=sys.stderr)
        return(1)
    if sample < 0:
        print("Sample size must be a ratio between 0 and 1 or a integer", file=sys.stderr)
        return(1)
    
    ## Setup
    np.random.seed(seed)

    ## Load dataset
    dataset = read_dataset(open(dataset, "r"))
    
    sequences = defaultdict(list)
    activity = defaultdict(list)

    for entry in dataset:
        fold = entry["fold"]
        sequence = dna_1hot(entry["sequence"].upper())
        offsets = generate_offsets(sequence.shape[0], length, stride, sample)

        for offset in offsets:
            sequences[fold].append(sequence[offset:offset+length, :])
            activity[fold].append(float(entry['activity']))

    for fold in sequences:
        sequences[fold] = np.array(sequences[fold])
        activity[fold]  = np.array(activity[fold])

        np.save(f"{prefix}-sequence.{fold}.npy", np.array(sequences[fold]))
        np.save(f"{prefix}-activity.{fold}.npy", np.array(activity[fold]))
    return(0)


if __name__ == "__main__":
    load_dotenv()

    parser = argparse.ArgumentParser(
        prog="src.data.make_dataset",
        help="Generates sequence and activity array files, used by src.models.train."
    )
    parser.add_argument('dataset', type=str, help="path to dataset reference table generated by src.data.make_dataset.")
    parser.add_argument('--prefix', type=str, default="dataset", help="prefix path to output files.")
    parser.add_argument('--length', type=int, default=int(os.getenv("SEQ_LENGTH", 25600)), help="input sequence length in bp.")
    parser.add_argument('--stride', type=int, default=1, help="step size used to generate sequence sample pool.")
    parser.add_argument('--sample', type=float, default=0.4, help="ratio or number of samples to extract from each dataset entry.")
    parser.add_argument('--seed', type=int, default=5, help="random number generator seed")

    args = parser.parse_args()

    print(args, file=sys.stderr)
    exit(main(**vars(args)))
```
