## Supplemental Code for "Iterative improvement of deep learning models using synthetic regulatory genomics": make_hotspot_block.py

```
#! /usr/bin/env python3

import os
import sys
import argparse
import tqdm
import numpy as np

from src.utils import pad_or_crop

from dotenv import load_dotenv
from intervaltree import IntervalTree

from collections import defaultdict
from typing import List, Dict, Tuple, Iterator


def read_bed(file: Iterator[str]) -> Dict[str, List[Tuple[int, int]]]:
    chunks = defaultdict(list)
    for line in file:
        line = line.rstrip()
        if line == "" or line.startswith("#"):
            continue
        chrom, start, end = line.split("\t")[:3]
        chunks[chrom].append((int(start), int(end)))
    return chunks


def mst_group(location: np.ndarray, max_length: int) -> np.ndarray:
    blockmin = np.arange(len(location))
    blockmax = np.arange(len(location))
    distance = np.diff(location)
    for i in np.argsort(distance):
        if distance[i] > max_length:
            break ## NO LONGER ABLE TO MERGE
        nstart = location[blockmin[i]]
        nend = location[blockmax[i+1]]
        if nend - nstart < max_length:
            ## MERGE IS BELOW THE LIMIT
            blockmin[blockmin == blockmin[i+1]] = blockmin[i]
            blockmax[blockmax == blockmax[i]] = blockmax[i+1]
    blockmin = np.unique(blockmin)
    blockmax = np.unique(blockmax)
    return(np.column_stack((blockmin, blockmax)))


def coverage(itree:IntervalTree, start:int, end:int) -> float:
    coverage = 0
    for chunk in itree[start:end]:
        coverage += min(end, chunk.end) - max(start, chunk.begin)
    return coverage / (end-start)


def main(hotspots: str, length:int, sample:int, seed:int, prefix:str, mappable:str) -> int:
    ## Check input
    if hotspots != "-" and not os.path.exists(hotspots):
       print(f"Unable to locate hotspots bedfile at `{hotspots}`", file=sys.stderr)
       return(1)
    
    ## SETUP
    rng = np.random.default_rng(seed=seed)
    hotspotfile = open(f"{prefix}.hotspots.bed", "w")
    blockfile = open(f"{prefix}.hotspots-block.bed", "w")

    ## Collect hotspots
    hotspots = read_bed(sys.stdin if hotspots == "-" else open(hotspots, "r"))
    ## Count number of events per chromossome
    print("Number of hotspots per chromosome:" , file=sys.stderr)
    print({ chrom: len(hotspots[chrom]) for chrom in hotspots }, file=sys.stderr)

    ## Group into blocks
    blocks = []
    for chrom in hotspots:
        chrom_hotspots = hotspots[chrom]
        chrom_positions = [sum(chunk)/2 for chunk in chrom_hotspots]
        chrom_positions = np.array(chrom_positions, dtype=np.float32)
        chrom_blocks = mst_group(chrom_positions, length)
        blocks += [
            (chrom, *pad_or_crop(chrom_hotspots[i][0], chrom_hotspots[j][1], length), chrom_hotspots[i:j+1])
            for (i, j) in chrom_blocks
        ]
    
    if mappable is not None:
        print(f"Filtering by mappable '{mappable}' coverage (> 90%) " , file=sys.stderr)
        mappable = read_bed(open(mappable, 'r'))
        mappable = dict((chrom, IntervalTree.from_tuples(mappable[chrom]))
                        for chrom in mappable)
        blen = len(blocks)
        blocks = [b for b in blocks if coverage(mappable[b[0]], b[1], b[2]) > 0.90]
        alen = len(blocks)
        print(f" - Dropped {blen-alen} blocks with low mappability" , file=sys.stderr)

    ## Subsample blocks
    if len(blocks) < sample:
        print(f"! Low sample space, skippping sampling", file=sys.stderr)
    else:
        index = np.arange(len(blocks))
        rng.shuffle(index)
        index = np.sort(index[:sample])
        blocks = [blocks[i] for i in index[:sample]]
    
    ## Save blocks
    si = 0
    bi = 0
    for (chrom, start, end, sites) in tqdm.tqdm(blocks):
        bi += 1
        start, end = pad_or_crop(start, end, length)
        sites = [site for site in sites if site[1] > start and site[0] <= end]
        sites_size = []
        sites_start = []
        for (site_start, site_end) in sites:
            si += 1
            site_start = max(start, site_start)
            site_end = min(end, site_end)
            print(chrom, site_start, site_end, f"HT{si}", sep="\t", file=hotspotfile)
            sites_size.append(str(site_end - site_start))
            sites_start.append(str(site_start - start))
        print(
            chrom, start, end, f"HB{bi}", 0, ".", start, end, "255,0,0",
            len(sites), ",".join(sites_size), ",".join(sites_start),
            sep="\t", file=blockfile)

    hotspotfile.close()
    blockfile.close()
    return(0)


if __name__ == "__main__":
    load_dotenv()

    parser = argparse.ArgumentParser(prog="make-hotspot-block")
    parser.add_argument('hotspots', type=str, default="-")
    parser.add_argument('--length', type=int, default=os.getenv("SEQ_LENGTH", 25600))
    parser.add_argument('--sample', type=int, default=2000)
    parser.add_argument('--seed', type=int, default=5)
    parser.add_argument('--mappable', type=str)
    parser.add_argument('--prefix', type=str, default="data/processed/hotspots")
    args = parser.parse_args()

    print(args, file=sys.stderr)
    exit(main(**vars(args)))
```
