## Supplemental Code for "Iterative improvement of deep learning models using synthetic regulatory genomics": make_track_dataset_h5.py

```
#! /usr/bin/env python3

import os
import sys
import csv
import argparse

import tqdm
import pysam
import pyBigWig
import h5py
import numpy as np

from dotenv import load_dotenv
from typing import Dict

from src.utils import dna_1hot

STAT = dict(
    sum  = lambda x: np.sum(x, axis=1, dtype=np.float32),
    mean = lambda x: np.mean(x, axis=1, dtype=np.float32)
)

class CovReader(object):
    def __init__(self, options:Dict):
        self._options = options
        self._covfile = pyBigWig.open(self._options["file"])
        if "clip" in self._options:
            self._clip = float(self._options["clip"])
        if "scale" in self._options:
            self._scale = float(self._options["scale"])
        self._func = STAT[self._options["sum_stat"]]
        
    def measure(self, chrom:str, start:int, end:int, width:int):
        cov = self._covfile.values(chrom, start, end, numpy=True)
        ## baseline to median
        baseline = np.percentile(cov, 50)
        baseline = np.nan_to_num(baseline)
        cov[np.isnan(cov)] = baseline

        cov = cov.reshape((end-start)//width, width)
        cov = self._func(cov)

        if self._scale is not None:
            cov = self._scale * cov
        if self._clip is not None:
            cov = np.clip(cov, -self._clip, self._clip)
        ## Clip to f16 limits
        cov = np.clip(cov, np.finfo(np.float16).min, np.finfo(np.float16).max)
        return(cov.astype(np.float16))

    def close(self):
        self._covfile.close()

    def __repr__(self):
        return(f"<CovReader {str(self._options)}>")


def main(mappable:str, tracks:str, reference:str, length:int, width:int = 128, step:int, cutoff:float, size:int, batches:int, seed:int, prefix:str) -> int:
    ## Check files exists
    if mappable != "-" and not os.path.exists(mappable):
        print(f"Unable to locate file at`{mappable}`", file=sys.stderr)
        return(1)
    if not os.path.exists(tracks):
        print(f"Unable to locate file at`{tracks}`", file=sys.stderr)
        return(1)

    ## Setup
    np.random.seed(seed)
    reference = pysam.Fastafile(reference)
    total_size = batches * size

    ## Read and prepare targets
    tracks = csv.DictReader(open(tracks, "r"), delimiter="\t")
    tracks = [CovReader(line) for line in tracks]

    ## Read mappable regions
    chrom_ix = []
    start_pool = []

    mappable = sys.stdin if mappable == "-" else open(mappable, "r")
    for line in tqdm.tqdm(mappable):
        chrom, start, end = line.rstrip().split("\t")[:3]
        start = int(start)
        end = int(end)

        if end - start < length:
            continue

        if chrom not in chrom_ix:
            chrom_ix.append(chrom)

        st = np.arange(start, end - length, step=step)
        ch = np.repeat(chrom_ix.index(chrom), st.shape[0])
        start_pool.append(np.stack((ch, st), axis=-1))
    start_pool = np.concatenate(start_pool)

    ## Sample sites
    if total_size > start_pool.shape[0]:
        total_size = start_pool.shape[0]
    ## Shuffle entries
    ix = np.random.choice(np.arange(start_pool.shape[0]), size=total_size)
    np.random.shuffle(ix)
    sites = start_pool[ix, :]

    dataset = open(f"{prefix}.bed", "w")

    counter = 0
    index = 0

    for batch in tqdm.tqdm(range(batches)):
        if counter >= total_size:
            break ## Early exit

        input = []
        outcome = []

        for ch, start in sites[index:, ...]:
            index += 1
            chrom = chrom_ix[ch]
            end = start + length

            out = [track.measure(chrom, start, end, width) for track in tracks]
            out = np.stack(out, axis=-1)

            if cutoff and np.max(np.std(out, 0)) < cutoff:
                continue ## SKIP if not informative
        
            seq = reference.fetch(chrom, start, end).upper()
            seq = dna_1hot(seq, dtype=np.float16)

            input.append(seq)
            outcome.append(out)
            print(chrom, start, end, sep="\t", file=dataset)

            counter += 1
            if counter > 0 and counter % size == 0 or counter >= total_size:
                break
        
        with h5py.File(f"{prefix}.{batch}.h5", 'w') as h5:
            h5.create_dataset('input', data=np.array(input), dtype='float16', compression="gzip")
            h5.create_dataset('outcome', data=np.array(outcome), dtype='float16', compression="gzip")

    dataset.close()
    for track in tracks:
        track.close()
    
    return(0)


if __name__ == "__main__":
    load_dotenv()
    
    parser = argparse.ArgumentParser(
        prog = "make-track-dataset",
        help = (
            "Builds a track training dataset for collection of bigwig {tracks}. " +
            "It will generate examples by extracting random {length} bp regions " +
            "contained in the targeted regions defined by {mappable}. "
            "Examples corresponding sequences are extracted from the {reference} " +
            "FASTA file. " +
            "Regions with standard signal deviation below {cutoff} are excluded " +
            "to avoid presenting uninformative sites."
        )
    )
    parser.add_argument('mappable', type=str,
        help = "BED file specifying targeted regions to be considered. Specifying `-` uses stdin instead.")
    parser.add_argument('tracks', type=str,
        help = "TSV file specifying targeted bigwig tracks to be trained.")
    parser.add_argument('--reference', type=str,
        help = "FASTA file from which examples sequences are extracted.")
    parser.add_argument(
        '--length', type=int, default=os.getenv("SEQ_LENGTH", 25600),
        help = "Length in bp of examples sequences. It must be a multiple of {width}.")
    parser.add_argument('--step', type=int, default=1,
        help = "Step size when sampling fragments within the {mappable} regions.")
    parser.add_argument('--size', type=int, default=10000,
        help = "Number of examples to be collected per dataset batch.")
    parser.add_argument('--batches', type=int, default=5,
        help = "Number of dataset batches to be collected.")
    parser.add_argument('--cutoff', type=float,
        help = "Minimal signal standard deviation required to consider an example.")
    parser.add_argument('--seed', type=int, default=5,
        help = "Random number generator seed to ensure reproducibility.")
    parser.add_argument('--prefix', type=str, default="track-dataset",
        help = "Output path prefix.")

    args = parser.parse_args()

    print(args, file=sys.stderr)
    exit(main(**vars(args)))
```
