## Supplemental Code for "Iterative improvement of deep learning models using synthetic regulatory genomics": payload_to_bedgraph.py

```
import os
import sys
import argparse
from typing import List

import numpy as np
from dotenv import load_dotenv

STEP=128


def parse_coordinate(coordinate:str) -> tuple:
    chrom, coord, *anno = coordinate.split(":")
    start, end = coord.split("-")
    return (chrom, int(start), int(end), anno)


def parse_payload(name, coordinates, length):
    locs = [
        parse_coordinate(coord)[:3] for coord in coordinates.split(',')
    ]
    loclen  = sum(loc[2] - loc[1] for loc in locs)

    chromix = list()
    locc = np.zeros(shape=(length,), dtype='uint32')
    locp = np.zeros(shape=(length,), dtype='uint32')
    offset = (length - loclen) // 2
    for chrom, start, end in locs:
        if chrom in chromix:
            cix = chromix.index(chrom)
        else:
            cix = len(chromix)
            chromix.append(chrom)
        
        size = end-start
        if offset < 0 and (offset + size) > 0:
            locc[0:(size + offset)] = cix
            locp[0:(size + offset)] = np.arange(start-offset, end)
        elif offset < length and (offset + size) >= length:
            locc[offset:length] = cix
            locp[offset:length] = np.arange(start, start + (length - offset))
        elif offset >= 0 and (offset + size) < length:
            locc[offset:(offset + size)] = cix
            locp[offset:(offset + size)] = np.arange(start, end)
        offset += size
    return(name, locc, locp, chromix)


def main(array: str, payload: str, tracks: List[int], outfile:str) -> int:
    if not os.path.exists(array):
        print(f"Unable to locate `{array}`", file=sys.stderr)
        return(1)
    if not os.path.exists(payload):
        print(f"Unable to locate `{payload}`", file=sys.stderr)
        return(1)
    
    outfile = sys.stdout if outfile == "-" else open(outfile, "w")
    
    array = np.load(array)
    length = STEP * array.shape[1]

    payloads = [line.rstrip().split("\t")[:2] for line in open(payload, "r")]
    payloads = [parse_payload(*payload, length) for payload in payloads]

    for i in range(len(payloads)):
        value = np.repeat(array[i, :, tracks], STEP, axis=-1)

        name, locc, locp, chromix = payloads[i]

        blockc = np.diff(np.append(locc[0], locc)) != 0
        blockp = np.diff(np.append(locp[0], locp)) > 1
        blockv = np.diff(np.append(0, np.repeat(np.arange(array.shape[1]), STEP))) != 0
        block = np.cumsum(blockc | blockp | blockv)

        for bi in np.unique(block):
            ix = np.argwhere(block == bi).flatten()
            chrom = chromix[locc[ix[0]]]
            start = locp[ix[0]]
            end = locp[ix[-1]] + 1
            val = value[:, ix[0]]
            
            if start == 0:
                continue

            print(chrom, start, end, name, *val, sep="\t", file=outfile)
    return(0)


if __name__ == "__main__":
    load_dotenv()

    parser = argparse.ArgumentParser("payload_to_bedgraph")
    parser.add_argument('array', type=str)
    parser.add_argument('payload', type=str)
    parser.add_argument('--outfile', type=str, default="-")
    parser.add_argument('--tracks', type=int, nargs='+', default=[10])

    args = parser.parse_args()
    print(args, file=sys.stderr)
    exit(main(**vars(args)))
```
