## Supplemental Code for "Iterative improvement of deep learning models using synthetic regulatory genomics": sample_block.py

```
#! /usr/bin/env python3

import os
import sys
import argparse
import csv

import tqdm
import numpy as np

from src.utils import pad_or_crop

from dotenv import load_dotenv
from intervaltree import IntervalTree

from collections import defaultdict
from typing import List, Dict, Tuple, Iterator


def read_bed(file: Iterator[str]) -> Dict[str, List[Tuple[int, int]]]:
    chunks = defaultdict(list)
    for line in file:
        line = line.rstrip()
        if line == "" or line.startswith("#"):
            continue
        chrom, start, end = line.split("\t")[:3]
        chunks[chrom].append((int(start), int(end)))
    return chunks


def mst_group(location: np.ndarray, max_length: int) -> np.ndarray:
    blockmin = np.arange(len(location))
    blockmax = np.arange(len(location))
    distance = np.diff(location)
    for i in np.argsort(distance):
        if distance[i] > max_length:
            break ## NO LONGER ABLE TO MERGE
        nstart = location[blockmin[i]]
        nend = location[blockmax[i+1]]
        if nend - nstart < max_length:
            ## MERGE IS BELOW THE LIMIT
            blockmin[blockmin == blockmin[i+1]] = blockmin[i]
            blockmax[blockmax == blockmax[i]] = blockmax[i+1]
    blockmin = np.unique(blockmin)
    blockmax = np.unique(blockmax)
    return(np.column_stack((blockmin, blockmax)))


def coverage(itree:IntervalTree, start:int, end:int) -> float:
    coverage = 0
    for chunk in itree[start:end]:
        coverage += min(end, chunk.end) - max(start, chunk.begin)
    return coverage / (end-start)


def main(mappable:str, length:int, sample:int, seed:int, output:str, **kwargs:dict) -> int:
    if not os.path.exists(mappable):
        print(f"Unable to locate sampable regions at {mappable}", file=sys.stderr)
        return(1)

    ## Set sample region
    mappable = csv.reader(open(mappable, 'r'), delimiter="\t")
    mappable = [ (row[0], int(row[1]), int(row[2])) for row in mappable ]
    ## Exclude regions smaller than target length
    mappable = [ row for row in mappable if (row[2] - row[1]) > length ]

    chroms = list(set(row[0] for row in mappable))
    
    cidx  = np.array([chroms.index(chrom) for chrom, _, _ in mappable])
    start = np.array([start for _, start, _ in mappable])
    end = np.array([end - length for _, _, end in mappable])
    size = end - start
    cumsize = np.cumsum(size)
    total_size = np.sum(size)

    print(f"Number of blocks:\t{len(mappable):,}", file=sys.stderr)
    print(f"Total blocks size:\t{total_size + length * len(end):,} bp", file=sys.stderr)

    rng = np.random.default_rng(seed=seed)
    positions = rng.integers(0, total_size, size = (sample,))
    positions = np.sort(positions)

    blocks = []
    ix = 0
    for pos in positions:
        while cumsize[ix] <= pos:
            ix += 1
        offset = cumsize[ix] - pos
        bchrom = chroms[cidx[ix]]
        bstart = start[ix] + offset
        blocks.append((bchrom, bstart, bstart + length))
    
    with open(output, 'w') as blockfile:
        csv.writer(blockfile, delimiter="\t").writerows(blocks)
    return(0)


if __name__ == "__main__":
    load_dotenv()

    parser = argparse.ArgumentParser(prog="sample-block")
    parser.add_argument('mappable', type=str)
    parser.add_argument('--length', type=int, default=os.getenv("SEQ_LENGTH", 25600))
    parser.add_argument('--sample', type=int, default=2000)
    parser.add_argument('--seed', type=int, default=5)
    parser.add_argument('--output', type=str, default="block.bed")
    args = parser.parse_args()

    print(args, file=sys.stderr)
    exit(main(**vars(args)))
```
