## Supplemental Code for "Iterative improvement of deep learning models using synthetic regulatory genomics": __init__.py

```
import os
import sys
import json

import numpy as np
import tensorflow as tf

from .enformer import Enformer, Sequential
from .layers import Attention, MultiHeadAttention


def load_or_create_modeldef(model:str, modeldef:dict = dict()) -> dict:
    ## Drop None values
    modeldef = { k: v for k, v in modeldef.items() if v is not None }
    ## Preload and combine previous definition
    jsonfile = f"{model}.json"
    if os.path.exists(jsonfile):
        modeldef = modeldef | json.load(open(jsonfile, 'r'))
    with open(jsonfile, 'w') as writer:
        json.dump(modeldef, writer)
    return(modeldef)


def load_baseline(model:Enformer, baseline:str) -> Enformer:
    if baseline == "original":
        ## DEFAULT ORIGINAL TO BASELINE WEIGHTS
        baseline = os.environ("ENFORMERBASELINE")
        if not os.path.exists(baseline):
            print(f"! Downloading baseline weights into '{baseline}'", file=sys.stderr)
            ## DOWNLOAD FROM GOOGLE STORAGE
            os.mkdir(baseline)            
            gspath = "gs://dm-enformer/models/enformer/sonnet_weights/*"
            for file in tf.io.gfile.glob(gspath):
                name = os.path.basename(file)
                tf.io.gfile.copy(file, f"{baseline}/{name}", overwrite=True)
    ## RESTORE WEIGHTS TO CHECKPOINT
    latest_checkpoint = tf.train.latest_checkpoint(baseline)
    checkpoint = tf.train.Checkpoint(module = model)
    checkpoint.restore(latest_checkpoint).expect_partial()
    print(f"! Restored lastest checkpoint: {latest_checkpoint}", file=sys.stderr)
    return(model)


def load_model_from_modeldef(model:str) -> dict:
    if model == "original":
        model = load_baseline(Enformer(), "original")
        mdict = dict(module = model, trunk = model.trunk, heads = model.heads)
    elif model == "tensorhub":
        import tensorflow_hub as hub

        model = hub.load(os.getenv("ENFORMERTENSORHUB")).model
        mdict = dict(tensorhub = model)
    else:
        modeldef = load_or_create_modeldef(model)
        # TODO: consider add check for necessary keys in modeldef
        # if not all(key in modeldef for key in ('key_size', 'value_size', 'num_heads', 'baseline')):
        #     return(None)
        ## FIX: Backwards compatibility to older folder format
        if not os.path.exists(model):
            model = f"{model}-checkpoint"
        baseline = Enformer()
        if not tf.train.latest_checkpoint(model):
            ## Restore trunk, heads[mouse] and heads[human] from original
            ## if model has no checkpoint save
            baseline = load_baseline(baseline, "original")
        finetune = build_finetune_head(
            modeldef["key_size"], modeldef["value_size"], modeldef["num_heads"])
        heads = baseline.heads | dict(finetune=finetune)
        mdict = dict(module = baseline, trunk = baseline.trunk, heads = heads)
        
        checkpoint = tf.train.Checkpoint(**mdict)
        latest_checkpoint = tf.train.latest_checkpoint(model)
        checkpoint.restore(latest_checkpoint).expect_partial()
        print(f"! Restored lastest checkpoint: {latest_checkpoint}", file=sys.stderr)
    return(mdict)


def build_finetune_head(key_size:int, value_size:int, num_heads:int) -> Sequential:
    if num_heads == 1:
        attention = Attention(value_size, key_size)
    else:
        attention = MultiHeadAttention(value_size, key_size, num_heads)
    pooling = tf.keras.layers.Flatten()
    linear = tf.keras.layers.Dense(1, activation="softplus")
    return Sequential([attention, pooling, linear])


def head_func(mdict:dict, head:str) -> Sequential:
    if "tensorhub" in mdict:
        return lambda x, is_training: mdict["tensorhub"].predict_on_batch(x)[head]
    return(Sequential([mdict["trunk"], mdict["heads"][head]]))


def build_finetune(base: Enformer, key:int, value:int, num_heads:int) -> Sequential:
    pooling = tf.keras.layers.Flatten()
    linear = tf.keras.layers.Dense(1, activation="softplus")
    if num_heads == 1:
        attention = Attention(value, key)
    else:
        attention = MultiHeadAttention(value, key, num_heads)
    return Sequential([base.trunk, attention, pooling, linear])


def load_original() -> Enformer:
    model = Enformer()
    latest_checkpoint = tf.train.latest_checkpoint(os.getenv("ENFORMERBASELINE"))
    checkpoint = tf.train.Checkpoint(module = model)
    checkpoint.restore(latest_checkpoint).expect_partial()
    return model


def restore_model(model:str, key:int, value:int, num_heads:int) -> (Enformer, Sequential):
    expression = lambda x, is_training: np.full([x.shape[0], 1], np.nan)
    
    if model == "tensorhub":
        import tensorflow_hub as hub

        model = hub.load(os.getenv("ENFORMERTENSORHUB")).model
        model = lambda x, is_training: model.predict_on_batch(x)
    elif model == "original":
        model = load_original()
    else:
        baseline = load_original()
        expression = build_finetune(baseline, key, value, num_heads)

        if not os.path.exists(model):
            model = f"{model}-checkpoint"
        checkpoint = tf.train.Checkpoint(module = baseline, finetune = expression)
        latest_checkpoint = tf.train.latest_checkpoint(model)
        checkpoint.restore(latest_checkpoint).expect_partial()
        print(f"! Restored lastest checkpoint: {latest_checkpoint}", file=sys.stderr)
        ## Override heads to ensure they are properly restored
        model = baseline
    return (model, expression)


def load_model(
        model:str, key:int, value:int, num_heads:int, head: str
        ) -> Sequential:
    head_layer = None
    if head not in ['human', 'mouse', 'finetune']:
        if model == "tensorhub" or not os.path.exists(head):
            raise ValueError(f"Unrecognized head '{head}'")
        
        headsize = os.getenv('HEADSIZE', 5) ## Enable custom sizes with minimal impact
        head_layer = tf.keras.layers.Dense(headsize, activation="softplus")
        checkpoint = tf.train.Checkpoint(head = head_layer)
        chkmanager = tf.train.CheckpointManager(checkpoint, head, max_to_keep=5)
        if chkmanager.latest_checkpoint:
            checkpoint.restore(chkmanager.latest_checkpoint)
            print(f"-> Restore from {chkmanager.latest_checkpoint} @ {head}", file=sys.stderr)

    if model == "tensorhub":
        import tensorflow_hub as hub

        model = hub.load(os.getenv("ENFORMERTENSORHUB")).model
        modeltrack = lambda x, is_training: model.predict_on_batch(x)[head]
    elif model == "original":
        model = load_original()
        if not head_layer:
            head_layer = model.heads[head]
        modeltrack = Sequential([model.trunk, head_layer])
    else:
        baseline = load_original()
        blank = Enformer()
        finetune = build_finetune(blank, key, value, num_heads)

        if not os.path.exists(model):
            model = f"{model}-checkpoint"
        checkpoint = tf.train.Checkpoint(module = blank, finetune = finetune)
        latest_checkpoint = tf.train.latest_checkpoint(model)
        checkpoint.restore(latest_checkpoint).expect_partial()
        print(f"! Restored lastest checkpoint: {latest_checkpoint}", file=sys.stderr)

        if head == "finetune":
            modeltrack = finetune
        elif head_layer:
            modeltrack = Sequential([finetune._layers[0], head_layer])
        else:
            modeltrack = Sequential([finetune._layers[0], baseline.heads[head]])
    return modeltrack
```
