## Supplemental Code for "Iterative improvement of deep learning models using synthetic regulatory genomics": attention_module.py

````
# Copyright 2021 DeepMind Technologies Limited
#
# Licensed under the Apache License, Version 2.0 (the "License");
# you may not use this file except in compliance with the License.
# You may obtain a copy of the License at
#
#      http://www.apache.org/licenses/LICENSE-2.0
#
# Unless required by applicable law or agreed to in writing, software
# distributed under the License is distributed on an "AS IS" BASIS,
# WITHOUT WARRANTIES OR CONDITIONS OF ANY KIND, either express or implied.
# See the License for the specific language governing permissions and
# limitations under the License.
"""TransformerBlock and MultiheadAttention modules used in the paper.

"Effective gene expression prediction from sequence by integrating long-range
interactions"

Žiga Avsec1, Vikram Agarwal2,4, Daniel Visentin1,4, Joseph R. Ledsam1,3,
Agnieszka Grabska-Barwinska1, Kyle R. Taylor1, Yannis Assael1, John Jumper1,
Pushmeet Kohli1, David R. Kelley2*

1 DeepMind, London, UK
2 Calico Life Sciences, South San Francisco, CA, USA
3 Google, Tokyo, Japan
4 These authors contributed equally.
* correspondence:

Example:
```
mha = MultiheadAttention(
    value_size=96,
    key_size=64,
    num_heads=8,
    relative_position_functions=['positional_features_sin_cos'])
mha(tf.ones((2, 1024, 96*8)), is_training=True)

# Transformer block as used in the paper
transformer_block = TransformerBlock(
    channels=96 * 8,
    dropout_rate=0.4,
    attention_kwargs=dict(
        value_size=96,
        key_size=64,
        num_heads=8,
        relative_positions=True,
        relative_position_symmetric=False,
        num_relative_position_features=None,
        relative_position_functions=['positional_features_exponential',
                                     'positional_features_central_mask',
                                     'positional_features_gamma'],
        positional_dropout_rate=0.01,
        attention_dropout_rate=0.05,
        )
    )
transformer_block(tf.ones((2, 1024, 96*8)), is_training=True)
```
"""
from typing import Any, Dict, List, Optional

import numpy as np
import sonnet as snt
import tensorflow as tf


class TransformerBlock(snt.Module):
  """Full transformer module block."""

  def __init__(
      self,
      channels: int,
      dropout_rate: float,
      attention_kwargs: Dict[str, Any],
      name: str = 'transformer_block',
  ):
    super().__init__(name=name)
    self.is_trainable = True
    self.mha_ln = snt.LayerNorm(axis=-1, create_scale=True, create_offset=True)
    self.mha = MultiheadAttention(**attention_kwargs)
    self.mha_dropout = snt.Dropout(dropout_rate)

    self.mlp_ln = snt.LayerNorm(axis=-1, create_scale=True, create_offset=True)
    self.mlp_linear1 = snt.Linear(channels * 2)
    self.mlp_dropout1 = snt.Dropout(dropout_rate)
    self.mlp_linear2 = snt.Linear(channels)
    self.mlp_dropout2 = snt.Dropout(dropout_rate)

  def __call__(self, inputs: tf.Tensor, is_training: bool) -> tf.Tensor:
    x = self.mha_ln(inputs)
    x = self.mha(x, is_training=is_training)
    x = self.mha_dropout(x, is_training=is_training)
    x += inputs  # Residual
    mha_output = x

    # MLP.
    x = self.mlp_ln(mha_output)
    x = self.mlp_linear1(x)
    x = self.mlp_dropout1(x, is_training=is_training)
    x = tf.nn.relu(x)
    x = self.mlp_linear2(x)
    x = self.mlp_dropout2(x, is_training=is_training)
    return x + mha_output


class MultiheadAttention(snt.Module):
  """Multi-head attention."""

  def __init__(self,
               value_size: int,
               key_size: int,
               num_heads: int,
               scaling: bool = True,
               attention_dropout_rate: float = 0.1,
               relative_positions: bool = False,
               relative_position_symmetric: bool = False,
               relative_position_functions: Optional[List[str]] = None,
               num_relative_position_features: Optional[int] = None,
               positional_dropout_rate: float = 0.1,
               zero_initialize: bool = True,
               initializer: Optional[snt.initializers.Initializer] = None,
               name: str = None):
    """Creates a MultiheadAttention module.

    Args:
      value_size: The size of each value embedding per head.
      key_size: The size of each key and query embedding per head.
      num_heads: The number of independent queries per timestep.
      scaling: Whether to scale the attention logits.
      attention_dropout_rate: Dropout rate for attention logits.
      relative_positions: Whether to use TransformerXL style relative attention.
      relative_position_symmetric: If True, the symmetric version of basis
        functions will be used. If False, a symmetric and asymmetric versions
        will be use.
      relative_position_functions: List of function names used for relative
        positional biases.
      num_relative_position_features: Number of relative positional features
        to compute. If None, `value_size * num_heads` is used.
      positional_dropout_rate: Dropout rate for the positional encodings if
        relative positions are used.
      zero_initialize: if True, the final linear layer will be 0 initialized.
      initializer: Initializer for the projection layers. If unspecified,
        VarianceScaling is used with scale = 2.0.
      name: Name of module.
    """
    super().__init__(name=name)
    self.is_trainable = True
    self._value_size = value_size
    self._key_size = key_size
    self._num_heads = num_heads
    self._attention_dropout_rate = attention_dropout_rate
    self._scaling = scaling
    self._relative_positions = relative_positions
    self._relative_position_symmetric = relative_position_symmetric
    self._relative_position_functions = relative_position_functions
    if num_relative_position_features is None:
      # num_relative_position_features needs to be divisible by the number of
      # relative positional functions *2 (for symmetric & asymmetric version).
      divisible_by = 2 * len(self._relative_position_functions)
      self._num_relative_position_features = (
          (self._value_size // divisible_by) * divisible_by)
    else:
      self._num_relative_position_features = num_relative_position_features
    self._positional_dropout_rate = positional_dropout_rate

    self._initializer = initializer
    if self._initializer is None:
      self._initializer = snt.initializers.VarianceScaling(scale=2.0)

    key_proj_size = self._key_size * self._num_heads
    embedding_size = self._value_size * self._num_heads

    self._q_layer = snt.Linear(
        key_proj_size,
        name='q_layer',
        with_bias=False,
        w_init=self._initializer)
    self._k_layer = snt.Linear(
        key_proj_size,
        name='k_layer',
        with_bias=False,
        w_init=self._initializer)
    self._v_layer = snt.Linear(
        embedding_size,
        name='v_layer',
        with_bias=False,
        w_init=self._initializer)
    w_init = snt.initializers.Zeros() if zero_initialize else self._initializer
    self._embedding_layer = snt.Linear(
        embedding_size,
        name='embedding_layer',
        w_init=w_init)

    # Create additional layers if using relative positions.
    if self._relative_positions:
      self._r_k_layer = snt.Linear(
          key_proj_size,
          name='r_k_layer',
          with_bias=False,
          w_init=self._initializer)
      self._r_w_bias = tf.Variable(
          self._initializer([1, self._num_heads, 1, self._key_size],
                            dtype=tf.float32),
          name='r_w_bias')
      self._r_r_bias = tf.Variable(
          self._initializer([1, self._num_heads, 1, self._key_size],
                            dtype=tf.float32),
          name='r_r_bias')

  def _multihead_output(self, linear, inputs):
    """Applies a standard linear to inputs and returns multihead output."""

    output = snt.BatchApply(linear)(inputs)  # [B, T, H * KV]
    num_kv_channels = output.shape[-1] // self._num_heads
    # Split H * Channels into separate axes.
    output = snt.reshape(output,
                         output_shape=[-1, self._num_heads, num_kv_channels])
    # [B, T, H, KV] -> [B, H, T, KV]
    return tf.transpose(output, [0, 2, 1, 3])

  def __call__(self,
               inputs,
               is_training=False):
    # Initialise the projection layers.
    embedding_size = self._value_size * self._num_heads
    seq_len = inputs.shape[1]

    # Compute q, k and v as multi-headed projections of the inputs.
    q = self._multihead_output(self._q_layer, inputs)  # [B, H, T, K]
    k = self._multihead_output(self._k_layer, inputs)  # [B, H, T, K]
    v = self._multihead_output(self._v_layer, inputs)  # [B, H, T, V]

    # Scale the query by the square-root of key size.
    if self._scaling:
      q *= self._key_size**-0.5

    if self._relative_positions:
      # For relative positions, we project positions to form relative keys.
      distances = tf.range(-seq_len + 1, seq_len, dtype=tf.float32)[tf.newaxis]
      positional_encodings = positional_features_all(
          positions=distances,
          feature_size=self._num_relative_position_features,
          seq_length=seq_len,
          feature_functions=self._relative_position_functions,
          symmetric=self._relative_position_symmetric)
      # [1, 2T-1, Cr]

      if is_training:
        positional_encodings = tf.nn.dropout(
            positional_encodings, rate=self._positional_dropout_rate)

      # [1, H, 2T-1, K]
      r_k = self._multihead_output(self._r_k_layer, positional_encodings)

      # Add shifted relative logits to content logits.
      # [B, H, T', T]
      content_logits = tf.matmul(q + self._r_w_bias, k, transpose_b=True)
      # [B, H, T', 2T-1]
      relative_logits = tf.matmul(
          q + self._r_r_bias, r_k, transpose_b=True)
      #  [B, H, T', T]
      relative_logits = relative_shift(relative_logits)
      logits = content_logits + relative_logits
    else:
      # [B, H, T', T]
      logits = tf.matmul(q, k, transpose_b=True)

    weights = tf.nn.softmax(logits)

    # Dropout on the attention weights.
    if is_training:
      weights = tf.nn.dropout(weights, rate=self._attention_dropout_rate)

    # Transpose and reshape the output.
    output = tf.matmul(weights, v)  # [B, H, T', V]
    output_transpose = tf.transpose(output, [0, 2, 1, 3])  # [B, T', H, V]

    # Final linear layer.
    attended_inputs = snt.reshape(
        output_transpose, output_shape=[embedding_size], preserve_dims=2)
    output = self._embedding_layer(attended_inputs)

    return output


def relative_shift(x):
  """Shift the relative logits like in TransformerXL."""
  # We prepend zeros on the final timescale dimension.
  to_pad = tf.zeros_like(x[..., :1])
  x = tf.concat([to_pad, x], -1)
  _, num_heads, t1, t2 = x.shape
  x = tf.reshape(x, [-1, num_heads, t2, t1])
  x = tf.slice(x, [0, 0, 1, 0], [-1, -1, -1, -1])
  x = tf.reshape(x, [-1, num_heads, t1, t2 - 1])
  x = tf.slice(x, [0, 0, 0, 0], [-1, -1, -1, (t2 + 1) // 2])
  return x


# Available feature functions:
def get_positional_feature_function(name):
  """Returns positional feature functions."""
  available = {
      'positional_features_exponential': positional_features_exponential,
      'positional_features_central_mask': positional_features_central_mask,
      'positional_features_gamma': positional_features_gamma,
      'positional_features_cosine': positional_features_cosine,
      'positional_features_linear_masks': positional_features_linear_masks,
      'positional_features_sin_cos': positional_features_sin_cos,
  }
  if name not in available:
    raise ValueError(f'Function {name} not available in {available.keys()}')
  return available[name]


def positional_features_all(positions: tf.Tensor,
                            feature_size: int,
                            seq_length: Optional[int] = None,
                            bin_size: Optional[int] = None,
                            feature_functions: Optional[List[str]] = None,
                            symmetric=False):
  """Compute relative positional encodings/features.

  Each positional feature function will compute/provide the same fraction of
  features, making up the total of feature_size.

  Args:
    positions: Tensor of relative positions of arbitrary shape.
    feature_size: Total number of basis functions.
    seq_length: Sequence length denoting the characteristic length that
      the individual positional features can use. This is required since the
      parametrization of the input features should be independent of `positions`
      while it could still require to use the total number of features.
    bin_size: Bin sized used to partition the sequence. This can be used to
      compute features on the absolute scale relative to the genome.
    feature_functions: List of different feature functions to use. Each function
      will take as argument: positions, sequence length and number of features
      to compute.
    symmetric: If True, the resulting features will be symmetric across the
      relative position of 0 (i.e. only absolute value of positions will
      matter). If false, then both the symmetric and asymmetric version
      (symmetric multiplied by sign(positions)) of the features will be used.

  Returns:
    Tensor of shape: `positions.shape + (feature_size,)`.
  """
  if feature_functions is None:
    feature_functions = ['positional_features_exponential',
                         'positional_features_central_mask',
                         'positional_features_gamma']
  num_components = len(feature_functions)  # 1 per each basis function
  if not symmetric:
    num_components = 2 * num_components

  # For now, we do not allow odd sized embeddings.
  if feature_size % num_components != 0:
    raise ValueError(
        f'feature_size has to be divisible by {num_components}')

  feature_functions = [get_positional_feature_function(f)
                       for f in feature_functions]
  num_basis_per_class = feature_size // num_components
  embeddings = tf.concat([f(tf.abs(positions), num_basis_per_class,
                            seq_length, bin_size)
                          for f in feature_functions],
                         axis=-1)
  if not symmetric:
    embeddings = tf.concat([embeddings,
                            tf.sign(positions)[..., tf.newaxis] * embeddings],
                           axis=-1)
  tf.TensorShape(embeddings.shape).assert_is_compatible_with(
      positions.shape + [feature_size])
  return embeddings


def _prepend_dims(x, num_dims):
  return tf.reshape(x, shape=[1] * num_dims + x.shape)


def positional_features_exponential(positions: tf.Tensor,
                                    feature_size: int,
                                    seq_length: Optional[int] = None,
                                    bin_size: Optional[int] = None,
                                    min_half_life: Optional[float] = 3.0):
  """Create exponentially decaying positional weights.

  Args:
    positions: Position tensor (arbitrary shape).
    feature_size: Number of basis functions to use.
    seq_length: Sequence length.
    bin_size: (unused). See `positional_features_all`.
    min_half_life: Smallest exponential half life in the grid of half lives.

  Returns:
    A Tensor with shape [2 * seq_length - 1, feature_size].
  """
  del bin_size  # Unused.
  if seq_length is None:
    seq_length = tf.reduce_max(tf.abs(positions)) + 1
  # Grid of half lifes from [3, seq_length / 2] with feature_size
  # distributed on the log scale.
  seq_length = tf.cast(seq_length, dtype=tf.float32)
  max_range = tf.math.log(seq_length) / tf.math.log(2.0)
  half_life = tf.pow(2.0, tf.linspace(min_half_life, max_range, feature_size))
  half_life = _prepend_dims(half_life, positions.shape.rank)
  positions = tf.abs(positions)
  outputs = tf.exp(-tf.math.log(2.0) / half_life * positions[..., tf.newaxis])
  tf.TensorShape(outputs.shape).assert_is_compatible_with(
      positions.shape + [feature_size])
  return outputs


def positional_features_central_mask(positions: tf.Tensor,
                                     feature_size: int,
                                     seq_length: Optional[int] = None,
                                     bin_size: Optional[int] = None):
  """Positional features using a central mask (allow only central features)."""
  del seq_length  # Unused.
  del bin_size  # Unused.
  center_widths = tf.pow(2.0, tf.range(1, feature_size + 1, dtype=tf.float32))
  center_widths = center_widths - 1
  center_widths = _prepend_dims(center_widths, positions.shape.rank)
  outputs = tf.cast(center_widths > tf.abs(positions)[..., tf.newaxis],
                    tf.float32)
  tf.TensorShape(outputs.shape).assert_is_compatible_with(
      positions.shape + [feature_size])
  return outputs


def gamma_pdf(x, concentration, rate):
  """Gamma probability distribution function: p(x|concentration, rate)."""
  log_unnormalized_prob = tf.math.xlogy(concentration - 1., x) - rate * x
  log_normalization = (tf.math.lgamma(concentration) -
                       concentration * tf.math.log(rate))
  return tf.exp(log_unnormalized_prob - log_normalization)


def positional_features_gamma(positions: tf.Tensor,
                              feature_size: int,
                              seq_length: Optional[int] = None,
                              bin_size: Optional[int] = None,
                              stddev=None,
                              start_mean=None):
  """Positional features computed using the gamma distributions."""
  del bin_size  # Unused.
  if seq_length is None:
    seq_length = tf.reduce_max(tf.abs(positions)) + 1
  if stddev is None:
    stddev = seq_length / (2 * feature_size)
  if start_mean is None:
    start_mean = seq_length / feature_size
  mean = tf.linspace(start_mean, seq_length, num=feature_size)
  mean = _prepend_dims(mean, positions.shape.rank)
  concentration = (mean / stddev)**2
  rate = mean / stddev**2
  probabilities = gamma_pdf(
      tf.abs(tf.cast(positions, dtype=tf.float32))[..., tf.newaxis],
      concentration, rate)
  probabilities += 1e-8  # To ensure numerical stability.
  outputs = probabilities / tf.reduce_max(probabilities,
                                          axis=1, keepdims=True)
  tf.TensorShape(outputs.shape).assert_is_compatible_with(
      positions.shape + [feature_size])
  return outputs


def positional_features_cosine(positions: tf.Tensor,
                               feature_size: int,
                               seq_length: Optional[int] = None,
                               bin_size: Optional[int] = None):
  """Cosine positional features."""
  del bin_size  # Unused.
  del seq_length  # Unused.
  periodicity = 1.25 * tf.pow(2.0, tf.range(0, feature_size, dtype=tf.float32))
  periodicity = _prepend_dims(periodicity, positions.shape.rank)

  outputs = tf.math.cos(2 * np.pi * positions[..., tf.newaxis] / periodicity)
  tf.TensorShape(outputs.shape).assert_is_compatible_with(
      positions.shape + [feature_size])
  return outputs


def positional_features_linear_masks(positions: tf.Tensor,
                                     feature_size: int,
                                     seq_length: Optional[int] = None,
                                     bin_size: Optional[int] = None):
  """Exponentially increasing point focuses."""
  del bin_size  # Unused.
  del seq_length  # Unused.
  distances = tf.range(0, feature_size, dtype=tf.float32)
  distances = _prepend_dims(distances, positions.shape.rank)
  outputs = tf.cast(distances == tf.abs(positions[..., tf.newaxis]),
                    dtype=tf.float32)

  tf.TensorShape(outputs.shape).assert_is_compatible_with(
      positions.shape + [feature_size])
  return outputs


def positional_features_sin_cos(positions: tf.Tensor,
                                feature_size: int,
                                seq_length: Optional[int] = None,
                                bin_size: Optional[int] = None,
                                max_time=10000.0):
  """Sine/cosine positional encodings."""
  del bin_size  # Unused.
  del seq_length  # Unused.
  if feature_size % 2 != 0:
    raise ValueError('feature_size needs to be divisible by 2.')
  i = tf.range(0, feature_size, 2, dtype=tf.float32)
  i = _prepend_dims(i, positions.shape.rank)

  # Concat sines and cosines and return.
  outputs = tf.concat([
      tf.sin(positions[..., tf.newaxis] / max_time**(i / feature_size)),
      tf.cos(positions[..., tf.newaxis] / max_time**(i / feature_size))], -1)

  tf.TensorShape(outputs.shape).assert_is_compatible_with(
      positions.shape + [feature_size])
  return outputs
````
