## Supplemental Code for "Iterative improvement of deep learning models using synthetic regulatory genomics": build.py

```
#! /usr/bin/env python3

import sys
import argparse
from src.models import load_or_create_modeldef, load_model_from_modeldef

def main(model:str, baseline:str, key_size:int, value_size:int, num_heads:int) -> int:
    modeldef = load_or_create_modeldef(model, dict(
        baseline=baseline,
        key_size=key_size,
        value_size=value_size,
        num_heads=num_heads))
    print(modeldef, file=sys.stderr)
    return(1)


if __name__ == "__main__":
    parser = argparse.ArgumentParser(prog="build-model")
    parser.add_argument('model', type=str)
    ## model parameters
    modeldef = parser.add_argument_group('model parameters')
    modeldef.add_argument('--baseline', type=str)
    modeldef.add_argument('--key-size', type=int)
    modeldef.add_argument('--value-size', type=int)
    modeldef.add_argument('--num-heads', type=int)

    args = parser.parse_args()
    print(args, file=sys.stderr)
    exit(main(**vars(args)))
```
