## Supplemental Code for "Iterative improvement of deep learning models using synthetic regulatory genomics": enformer.py

```
# Copyright 2021 DeepMind Technologies Limited
#
# Licensed under the Apache License, Version 2.0 (the "License");
# you may not use this file except in compliance with the License.
# You may obtain a copy of the License at
#
#      http://www.apache.org/licenses/LICENSE-2.0
#
# Unless required by applicable law or agreed to in writing, software
# distributed under the License is distributed on an "AS IS" BASIS,
# WITHOUT WARRANTIES OR CONDITIONS OF ANY KIND, either express or implied.
# See the License for the specific language governing permissions and
# limitations under the License.
"""Tensorflow implementation of Enformer model.

"Effective gene expression prediction from sequence by integrating long-range
interactions"

Žiga Avsec1, Vikram Agarwal2,4, Daniel Visentin1,4, Joseph R. Ledsam1,3,
Agnieszka Grabska-Barwinska1, Kyle R. Taylor1, Yannis Assael1, John Jumper1,
Pushmeet Kohli1, David R. Kelley2*

1 DeepMind, London, UK
2 Calico Life Sciences, South San Francisco, CA, USA
3 Google, Tokyo, Japan
4 These authors contributed equally.
* correspondence:
"""
import inspect
from typing import Any, Callable, Dict, Optional, Text, Union, Iterable

from . import attention_module
import numpy as np
import sonnet as snt
import tensorflow as tf

SEQUENCE_LENGTH = 196_608
BIN_SIZE = 128
TARGET_LENGTH = 896
TARGET_LENGTH = 100


class Enformer(snt.Module):
  """Main model."""

  def __init__(self,
               channels: int = 1536,
               num_transformer_layers: int = 11,
               num_heads: int = 8,
               pooling_type: str = 'attention',
               name: str = 'enformer',
               target_length: int = -1):
    """Enformer model.

    Args:
      channels: Number of convolutional filters and the overall 'width' of the
        model.
      num_transformer_layers: Number of transformer layers.
      num_heads: Number of attention heads.
      pooling_type: Which pooling function to use. Options: 'attention' or max'.
      name: Name of sonnet module.
    """
    super().__init__(name=name)
    # pylint: disable=g-complex-comprehension,g-long-lambda,cell-var-from-loop
    heads_channels = {'human': 5313, 'mouse': 1643}
    dropout_rate = 0.4
    assert channels % num_heads == 0, ('channels needs to be divisible '
                                       f'by {num_heads}')
    whole_attention_kwargs = {
        'attention_dropout_rate': 0.05,
        'initializer': None,
        'key_size': 64,
        'num_heads': num_heads,
        'num_relative_position_features': channels // num_heads,
        'positional_dropout_rate': 0.01,
        'relative_position_functions': [
            'positional_features_exponential',
            'positional_features_central_mask',
            'positional_features_gamma'
        ],
        'relative_positions': True,
        'scaling': True,
        'value_size': channels // num_heads,
        'zero_initialize': True
    }

    trunk_name_scope = tf.name_scope('trunk')
    trunk_name_scope.__enter__()
    # lambda is used in Sequential to construct the module under tf.name_scope.
    def conv_block(filters, width=1, w_init=None, name='conv_block', **kwargs):
      return Sequential(lambda: [
          snt.distribute.CrossReplicaBatchNorm(
              create_scale=True,
              create_offset=True,
              scale_init=snt.initializers.Ones(),
              moving_mean=snt.ExponentialMovingAverage(0.9),
              moving_variance=snt.ExponentialMovingAverage(0.9)),
          gelu,
          snt.Conv1D(filters, width, w_init=w_init, **kwargs)
      ], name=name)

    stem = Sequential(lambda: [
        snt.Conv1D(channels // 2, 15),
        Residual(conv_block(channels // 2, 1, name='pointwise_conv_block')),
        pooling_module(pooling_type, pool_size=2),
    ], name='stem')

    filter_list = exponential_linspace_int(start=channels // 2, end=channels,
                                           num=6, divisible_by=128)
    conv_tower = Sequential(lambda: [
        Sequential(lambda: [
            conv_block(num_filters, 5),
            Residual(conv_block(num_filters, 1, name='pointwise_conv_block')),
            pooling_module(pooling_type, pool_size=2),
            ],
                   name=f'conv_tower_block_{i}')
        for i, num_filters in enumerate(filter_list)], name='conv_tower')

    # Transformer.
    def transformer_mlp():
      return Sequential(lambda: [
          snt.LayerNorm(axis=-1, create_scale=True, create_offset=True),
          snt.Linear(channels * 2),
          snt.Dropout(dropout_rate),
          tf.nn.relu,
          snt.Linear(channels),
          snt.Dropout(dropout_rate)], name='mlp')

    transformer = Sequential(lambda: [
        Sequential(lambda: [
            Residual(Sequential(lambda: [
                snt.LayerNorm(axis=-1,
                              create_scale=True, create_offset=True,
                              scale_init=snt.initializers.Ones()),
                attention_module.MultiheadAttention(**whole_attention_kwargs,
                                                    name=f'attention_{i}'),
                snt.Dropout(dropout_rate)], name='mha')),
            Residual(transformer_mlp())], name=f'transformer_block_{i}')
        for i in range(num_transformer_layers)], name='transformer')

    # Masking with identity layer to remove sequence length requirements
    # crop_final = TargetLengthCrop1D(TARGET_LENGTH, name='target_input')
    crop_final = Identity(name='target_input')
    if target_length > 0:
      crop_final = TargetLengthCrop1D(target_length, name='target_input')

    final_pointwise = Sequential(lambda: [
        conv_block(channels * 2, 1),
        snt.Dropout(dropout_rate / 8),
        gelu], name='final_pointwise')

    self._trunk = Sequential([stem,
                              conv_tower,
                              transformer,
                              crop_final,
                              final_pointwise],
                             name='trunk')
    trunk_name_scope.__exit__(None, None, None)

    with tf.name_scope('heads'):
      self._heads = {
          head: Sequential(
              lambda: [snt.Linear(num_channels), tf.nn.softplus],
              name=f'head_{head}')
          for head, num_channels in heads_channels.items()
      }
    # pylint: enable=g-complex-comprehension,g-long-lambda,cell-var-from-loop

  @property
  def trunk(self):
    return self._trunk

  @property
  def heads(self):
    return self._heads

  def __call__(self, inputs: tf.Tensor,
               is_training: bool) -> Dict[str, tf.Tensor]:
    trunk_embedding = self.trunk(inputs, is_training=is_training)
    return {
        head: head_module(trunk_embedding, is_training=is_training)
        for head, head_module in self.heads.items()
    }

  @tf.function(input_signature=[
      tf.TensorSpec([None, SEQUENCE_LENGTH, 4], tf.float32)])
  def predict_on_batch(self, x):
    """Method for SavedModel."""
    return self(x, is_training=False)


class Identity(snt.Module):
  def __init__(self, name:str = 'identity'):
    super().__init__(name=name)
  
  def __call__(self, inputs:tf.Tensor) -> tf.Tensor:
    return inputs

class TargetLengthCrop1D(snt.Module):
  """Crop sequence to match the desired target length."""

  def __init__(self,
               target_length: Optional[int],
               name: str = 'target_length_crop'):
    super().__init__(name=name)
    self._target_length = target_length

  def __call__(self, inputs):
    if self._target_length is None:
      return inputs
    trim = (inputs.shape[-2] - self._target_length) // 2
    if trim < 0:
      raise ValueError('inputs longer than target length')
    elif trim == 0:
      return inputs
    else:
      return inputs[..., trim:-trim, :]


class Sequential(snt.Module):
  """snt.Sequential automatically passing is_training where it exists."""

  def __init__(self,
               layers: Optional[Union[Callable[[], Iterable[snt.Module]],
                                      Iterable[Callable[..., Any]]]] = None,
               name: Optional[Text] = None):
    super().__init__(name=name)
    if layers is None:
      self._layers = []
    else:
      # layers wrapped in a lambda function to have a common namespace.
      if hasattr(layers, '__call__'):
        layers = layers()
      self._layers = [layer for layer in layers if layer is not None]

  def __call__(self, inputs: tf.Tensor, is_training: bool, **kwargs):
    outputs = inputs
    for _, mod in enumerate(self._layers):
      try:
        outputs = mod(outputs, is_training=is_training, **kwargs)
      except TypeError:
        outputs = mod(outputs, **kwargs)
    return outputs


def pooling_module(kind, pool_size):
  """Pooling module wrapper."""
  if kind == 'attention':
    return SoftmaxPooling1D(pool_size=pool_size, per_channel=True,
                            w_init_scale=2.0)
  elif kind == 'max':
    return tf.keras.layers.MaxPool1D(pool_size=pool_size, padding='same')
  else:
    raise ValueError(f'Invalid pooling kind: {kind}.')


class SoftmaxPooling1D(snt.Module):
  """Pooling operation with optional weights."""

  def __init__(self,
               pool_size: int = 2,
               per_channel: bool = False,
               w_init_scale: float = 0.0,
               name: str = 'softmax_pooling'):
    """Softmax pooling.

    Args:
      pool_size: Pooling size, same as in Max/AvgPooling.
      per_channel: If True, the logits/softmax weights will be computed for
        each channel separately. If False, same weights will be used across all
        channels.
      w_init_scale: When 0.0 is equivalent to avg pooling, and when
        ~2.0 and `per_channel=False` it's equivalent to max pooling.
      name: Module name.
    """
    super().__init__(name=name)
    self._pool_size = pool_size
    self._per_channel = per_channel
    self._w_init_scale = w_init_scale
    self._logit_linear = None

  @snt.once
  def _initialize(self, num_features):
    self._logit_linear = snt.Linear(
        output_size=num_features if self._per_channel else 1,
        with_bias=False,  # Softmax is agnostic to shifts.
        w_init=snt.initializers.Identity(self._w_init_scale))

  def __call__(self, inputs):
    _, length, num_features = inputs.shape
    self._initialize(num_features)
    inputs = tf.reshape(
        inputs,
        (-1, length // self._pool_size, self._pool_size, num_features))
    return tf.reduce_sum(
        inputs * tf.nn.softmax(self._logit_linear(inputs), axis=-2),
        axis=-2)


class Residual(snt.Module):
  """Residual block."""

  def __init__(self, module: snt.Module, name='residual'):
    super().__init__(name=name)
    self._module = module

  def __call__(self, inputs: tf.Tensor, is_training: bool, *args,
               **kwargs) -> tf.Tensor:
    return inputs + self._module(inputs, is_training, *args, **kwargs)


def gelu(x: tf.Tensor) -> tf.Tensor:
  """Applies the Gaussian error linear unit (GELU) activation function.

  Using approximiation in section 2 of the original paper:
  https://arxiv.org/abs/1606.08415

  Args:
    x: Input tensor to apply gelu activation.
  Returns:
    Tensor with gelu activation applied to it.
  """
  return tf.nn.sigmoid(1.702 * x) * x


def one_hot_encode(sequence: str,
                   alphabet: str = 'ACGT',
                   neutral_alphabet: str = 'N',
                   neutral_value: Any = 0,
                   dtype=np.float32) -> np.ndarray:
  """One-hot encode sequence."""
  def to_uint8(string):
    return np.frombuffer(string.encode('ascii'), dtype=np.uint8)
  hash_table = np.zeros((np.iinfo(np.uint8).max, len(alphabet)), dtype=dtype)
  hash_table[to_uint8(alphabet)] = np.eye(len(alphabet), dtype=dtype)
  hash_table[to_uint8(neutral_alphabet)] = neutral_value
  hash_table = hash_table.astype(dtype)
  return hash_table[to_uint8(sequence)]


def exponential_linspace_int(start, end, num, divisible_by=1):
  """Exponentially increasing values of integers."""
  def _round(x):
    return int(np.round(x / divisible_by) * divisible_by)

  base = np.exp(np.log(end / start) / (num - 1))
  return [_round(start * base**i) for i in range(num)]


def accepts_is_training(module):
  return 'is_training' in list(inspect.signature(module.__call__).parameters)
```
