## Supplemental Code for "Iterative improvement of deep learning models using synthetic regulatory genomics": layers.py

```
import numpy as np
import tensorflow as tf
import sonnet as snt


class Blank(tf.keras.layers.Layer):
    def __init__(self, size:int, name:str="blank"):
        self._size = size
    
    def call(self, inputs: tf.Tensor, training:bool = False) -> tf.Tensor:
        return tf.fill((inputs.shape[0], self._size), np.nan)


class Attention(tf.keras.layers.Layer):
    def __init__(self, value_size:int, key_size:int, dropout_rate:float =0.1, name:str="attention"):
        super().__init__(name=name)
        self._value_size = value_size
        self._key_size = key_size
        self._dropout_rate = dropout_rate
        self._initializer = snt.initializers.VarianceScaling(scale=2.0)
        
        self._q = tf.keras.layers.Dense(self._key_size, use_bias=False, kernel_initializer=self._initializer)
        self._k = tf.keras.layers.Dense(self._key_size, use_bias=False, kernel_initializer=self._initializer)
        self._v = tf.keras.layers.Dense(self._value_size, use_bias=False, kernel_initializer=self._initializer)
    
    def call(self, inputs: tf.Tensor, training:bool = False) -> tf.Tensor:
        q = self._q(inputs) # [B, K]
        k = self._k(inputs) # [B, K]
        v = self._v(inputs) # [B, V]
        ## scaling
        q *= self._key_size ** -0.5
        ## compute key weights
        logits = tf.matmul(q, k, transpose_b=True) # [B, B]
        weights = tf.nn.softmax(logits)
        if training:
            weights = tf.nn.dropout(weights, rate=self._dropout_rate)
        ## generate output
        return tf.matmul(weights, v)  # [B, V]


class MultiHeadAttention(tf.keras.layers.Layer):
    def __init__(self, value_size:int, key_size:int, heads:int, dropout_rate:float =0.1, name:str="attention"):
        super().__init__(name=name)
        self._value_size = value_size
        self._key_size = key_size
        self._heads = heads
        self._dropout_rate = dropout_rate
        self._initializer = snt.initializers.VarianceScaling(scale=2.0)
        
        kproj = self._key_size * self._heads
        vproj = self._value_size * self._heads

        self._q = tf.keras.layers.Dense(kproj, use_bias=False, kernel_initializer=self._initializer)
        self._k = tf.keras.layers.Dense(kproj, use_bias=False, kernel_initializer=self._initializer)
        self._v = tf.keras.layers.Dense(vproj, use_bias=False, kernel_initializer=self._initializer)

    def _transpose(self, x):
        kv = x.shape[-1] // self._heads
        x = tf.reshape(x, x.shape[:-1] + [self._heads, kv])
        if len(x.shape) > 3:
            x = tf.transpose(x, (0, 2, 1, 3))
        return x

    def _untranspose(self, x):
        if len(x.shape) > 3:
            x = tf.transpose(x, (0, 2, 1, 3))
        kv = x.shape[-1]
        x = tf.reshape(x, x.shape[:-2] + [self._heads * kv])
        return x
    
    def call(self, inputs: tf.Tensor, training:bool = False) -> tf.Tensor:
        scale = (self._key_size ** -0.5)
        q = self._transpose(self._q(inputs)) * scale # [B, H, K]
        k = self._transpose(self._k(inputs)) # [B, H, K]
        v = self._transpose(self._v(inputs)) # [B, H, V]
        ## compute key weights
        weights = tf.nn.softmax(tf.matmul(q, k, transpose_b=True)) # [B, B]
        if training:
            weights = tf.nn.dropout(weights, rate=self._dropout_rate)
        ## generate output
        return self._untranspose(tf.matmul(weights, v))  # [B, V]
```
