## Supplemental Code for "Iterative improvement of deep learning models using synthetic regulatory genomics": predict.py

```
#! /usr/bin/env python3

import os
import sys
import argparse

import tqdm
import numpy as np
import tensorflow as tf

from dotenv import load_dotenv
from src.utils import read_dataset, generate_offsets, batch_iterator, dna_1hot
from src.models import restore_model


def wrap(vec: np.ndarray, size:int) -> np.ndarray:
    nr, nc = vec.shape
    return(np.reshape(vec, (nr, nc // size, size)).max(axis=2))


def main(model:str, dataset:str, seed:int, batch:int, length:int,
         key_size:int, value_size:int, num_heads:int,
         head:str, track:int, stride:int, sample:float, output:str) -> int:
    if not os.path.exists(dataset):
        print(f"Unable to locate dataset at: `{dataset}`", file=sys.stderr)
        return(1)
    if model not in ['original', 'tensorhub'] and not os.path.exists(model):
        print(f"Unable to locate model checkpoints at: `{model}`", file=sys.stderr)
        return(1)
    if sample < 0:
        print("Sample size must be a ratio between 0 and 1 or a integer", file=sys.stderr)
        return(1)
    ## SETUP
    np.random.seed(seed)
    outfile = open(output, "w") if output != "-" else sys.stdout

    ## Load dataset
    print("! Load dataset", file=sys.stderr)
    dataset = read_dataset(open(dataset, "r"))
    
    ## Load model
    print("! Load model", file=sys.stderr)
    model_track, model_expression = restore_model(model, key_size, value_size, num_heads)
    
    ## Run batch analysis
    print("! Generating predictions", file=sys.stderr)
    print("index", "name", "group", "fold", "offset", "activity", "predicted",
          "peaksum", "peak", sep="\t", file=outfile)
    
    for entry in tqdm.tqdm(dataset):
        sequence = dna_1hot(entry['sequence'].upper())
        annotations_labels = [anno[0] for anno in entry['annotation']]
        annotations_matrix = []
        for anno in entry['annotation']:
            vec = np.zeros(sequence.shape[0])
            vec[anno[1]:anno[2]] = 1
            annotations_matrix.append(vec)
        if len(entry['annotation']) == 0:
            ## Add blank when no annotation available
            annotations_matrix.append(np.zeros(sequence.shape[0]))
        annotations_matrix = np.array(annotations_matrix)
        ## Generate strides and subsample
        offsets = generate_offsets(sequence.shape[0], length, stride, sample)

        for batch_offsets in batch_iterator(offsets, batch):
            batch_sequence = np.array([
                sequence[offset:offset+length, :] for offset in batch_offsets ])
            prediction = model_expression(batch_sequence, is_training=False)
            prediction_track = model_track(batch_sequence, is_training=False)[head][:, :, track]

            wrap_size = int(length / prediction_track.shape[1])
            wrap_anno = np.array([
                wrap(annotations_matrix[:, offset:offset+length], wrap_size)
                for offset in batch_offsets ])
            for i in range(len(batch_offsets)):
                peak = np.max(prediction_track[i, :] * wrap_anno[i, :, :], axis=1)
                peaksum = np.sum(peak)
                peak = [f"{label}={peak[i]:.3f}" for i, label in enumerate(annotations_labels)]
                print(
                    entry['index'], entry['name'], entry['group'], entry['fold'],
                    batch_offsets[i], entry['activity'],
                    f"{prediction[i, 0]:.3f}", f"{peaksum:.3f}",
                    ";".join(peak), sep = "\t", file=outfile)
    return(0)


if __name__ == "__main__":
    load_dotenv()

    parser = argparse.ArgumentParser(
        prog="predict",
        description=(
            "Predicts {model} activity (from finetuning head) and maximum signal " +
            "at annotated sites for each {dataset} entry. " +
            "Predictions of each entry are conducted on a random sample of all " +
            "possible sequence strides."
        ))
    parser.add_argument('model', type=str,
        help="path to load finetuned model. Indicating `original` will load weights from $ENFORMERBASELINE and `tensorhub` will load weights from ENFORMERTENSORHUB.")
    parser.add_argument('dataset', type=str, help="path to dataset table.")
    parser.add_argument('--seed', type=int, default=5, help="random number generator seed")
    parser.add_argument('--batch', type=int, default=4,
        help="number of samples to be evaluated simultaneously. Note this has large implications to the memory requirements.")
    parser.add_argument('--length', type=int, default=os.getenv('SEQ_LENGTH', 25600),
        help="input/output sequence length in bp.")
    
    ## model parameters
    modeldef = parser.add_argument_group('model definition')
    modeldef.add_argument('--key-size', type=int, default=64, help='finetuning attention layer key size')
    modeldef.add_argument('--value-size', type=int, default=64, help='finetuning attention layer value size')
    modeldef.add_argument('--num-heads', type=int, default=1, help='number of independent finetuning attention heads')

    ## output parameters
    outputdef = parser.add_argument_group('output definition')
    outputdef.add_argument('--head', type=str, choices=['mouse', 'human'], default='mouse',
        help="Enformer output head (either mouse or human) to evaluate annotated sites signal.")
    outputdef.add_argument('--track', type=int, default=10,
        help="index of Enformer output track of interest to evaluate annotated sites signal.")
    outputdef.add_argument('--stride', type=int, default=1, help="step size used to generate sequence sample pool.")
    outputdef.add_argument('--sample', type=float, default=0.4,
        help="ratio or number of samples to extract from each dataset entry.")
    outputdef.add_argument('--output', type=str, default="-",
        help="resulting table output path. Print to stdout if `-`.")

    args = parser.parse_args()
    print(args)
    exit(main(**vars(args)))
```
