## Supplemental Code for "Iterative improvement of deep learning models using synthetic regulatory genomics": predict_array.py

```
#! /usr/bin/env python3

import os
import sys
import argparse

import tqdm
import pysam
import numpy as np
import tensorflow as tf

from dotenv import load_dotenv
from src.utils import pad_or_crop, dna_1hot
from src.models import load_model


def main(
        model:str, bedfile:str,
        reference:str, length:int,
        head: str, strides:int, output: str,
        key_size:int, value_size:int, num_heads:int) -> int:
    ## Check input
    for file in [bedfile, reference]:
        if not os.path.exists(file):
            print(f"! Unable to locate `{file}`", file=sys.stderr)
            return(1)
        
    if head not in ['mouse', 'human', 'finetune'] and not os.path.exists(file):
        print(f"! Unrecognized head `{head}`", file=sys.stderr)
        return(1)

    ## SETUP
    stride_it = [(i, i+length) for i in range(0, strides + 1)]
    reference = pysam.FastaFile(args.reference)

    ## Prepare track
    print("! Setup track", file=sys.stderr)
    total_length = length + strides
    targets = [line.rstrip().split("\t") for line in open(bedfile, "r")]
    targets = [(x[0], int(x[1]), int(x[2]))
               for x in targets if not x[0].startswith('#') and len(x) >= 3]
    targets = [(x[0], *pad_or_crop(x[1], x[2], total_length)) for x in targets]

    print("! Load model", file=sys.stderr)
    model = load_model(model, key_size, value_size, num_heads, head)
  
    ## Predict track signal
    print("! Predict tracks", file=sys.stderr)
    traget_prediction = []
    shape = None
    for (chrom, start, end) in tqdm.tqdm(targets):
        sequence = dna_1hot(reference.fetch(chrom, start, end).upper())
        sequence = np.array([sequence[st:ed] for st, ed in stride_it])
        try:
            prediction = []
            for bi in range(0, sequence.shape[0], 3):
                bs = sequence[bi:min(sequence.shape[0], bi+3), :, :]
                pd = model(bs, is_training=False)
                prediction.append(pd)
            prediction = np.concatenate(prediction)
            # prediction = model(sequence, is_training=False)
            prediction = np.mean(prediction, axis=0)
            shape = prediction.shape
            traget_prediction.append(prediction)
        except Exception as err:
            ## It will fail if none are present
            print(f"Unable to generate predictions at {chrom}:{start}-{end}", file=sys.stderr)
            print(err, file=sys.stderr)
            if shape is None:
                return(1)
            traget_prediction.append(np.full(shape, np.nan))            
    np.save(output, np.array(traget_prediction))    
    return(0)
    

if __name__ == "__main__":
    load_dotenv()

    parser = argparse.ArgumentParser(
        prog="predict-array",
        description = (
            "Generated the {model} predictions over targeted regions specified " +
            "in the {bedfile} and saves them into a numpy array. " +
            "Each region are resized to be a multiple of {length} and broken " +
            "into non-overlapping fragments for prediction. " +
            "Each fragment sequence are extracted from {reference} FASTA file " +
            "and predictions are measured as the average of {strides}x 1-bp " +
            "shifts centered at the fragment region."
        )
    )
    parser.add_argument('model', type=str,
        help = "Path to model weights or either `original` or `tensorhub` which will load weights from ENFORMERBASELINE and ENFORMERTENSORHUB, respectively.")
    parser.add_argument('bedfile', type=str, help = "BED file specifying targeted regions.")
    parser.add_argument('--reference', type=str,
        help = "Path to reference FASTA file from which the targeted regions sequences are extracted.")
    parser.add_argument(
        '--length', type=int, default=os.getenv('SEQ_LENGTH', 25600),
        help = "Output fragments length in bp.")
    ## model parameters
    modeldef = parser.add_argument_group('model definition')
    modeldef.add_argument('--key-size', type=int, default=64,
        help='Finetuning attention layer key size')
    modeldef.add_argument('--value-size', type=int, default=64,
        help='Finetuning attention layer value size')
    modeldef.add_argument('--num-heads', type=int, default=1,
        help='Number of independent finetuning attention heads')
    ## output parameters
    outputdef = parser.add_argument_group('output definition')
    outputdef.add_argument('--head', type=str, default='mouse',
        help = "Enformer output head (either mouse or human) or path to a head weights.")
    outputdef.add_argument('--strides', type=int, default=10,
        help = "Number of 1bp shift to average the predicted signal. Note that it will compute all strides in a single batch.")
    outputdef.add_argument('--output', type=str, default="array.npy",
        help = "Path to save resulting prediction array.")

    args = parser.parse_args()
    print(args, file=sys.stderr)
    exit(main(**vars(args)))
```
