## Supplemental Code for "Iterative improvement of deep learning models using synthetic regulatory genomics": predict_bedgraph.py

```
#! /usr/bin/env python3

import os
import sys
import argparse

import tqdm
import pysam
import numpy as np
import tensorflow as tf

from dotenv import load_dotenv
from src.utils import dna_1hot
from src.models import load_model, load_model_from_modeldef, head_func


def resize_and_break(chrom:str, start:int, end:int, length:int) -> list[tuple[str, int, int]]:
    size = (end-start)
    nblock = int(np.ceil(size / length))
    nstart = (start + end - (nblock * length)) // 2
    return([
        (chrom, nstart + i * length, nstart + (i+1) * length)
        for i in range(nblock)
    ])


def main(
        bedfile:str, model:str, reference:str, length:int,
        key_size:int, value_size:int, num_heads:int, extend:bool,
        head:str, strides:int, tracks:list[int], prefix:str) -> int:
    ## check input
    if not os.path.exists(reference):
        print(f"! Unable to locate `{reference}`", file=sys.stderr)
        return(1)
    if bedfile != "-" and not os.path.exists(bedfile):
        print(f"! Unable to locate `{bedfile}`", file=sys.stderr)
        return(1)
    
    ## Setup run
    it_stride = [(i, i+length) for i in range(0, strides + 1)]
    reference = pysam.FastaFile(reference) 
    if (prefix is None):
        prefix = os.path.basename(model)

    ## Prepare track
    print("! Setup targets", file=sys.stderr)
    lshift = strides // 2
    rshift = strides - lshift

    targets = sys.stdin if bedfile == "-" else open(bedfile, "r")
    targets = [line.rstrip().split("\t") for line in targets]
    targets = [
        (*loc, reference.fetch(loc[0], loc[1] - lshift, loc[2] + rshift))
        for x in targets
        for loc in resize_and_break(x[0], int(x[1]), int(x[2]), length)
    ]

    print("! Load model", file=sys.stderr)
    mdict = load_model_from_modeldef(model)
    model = head_func(mdict, head)
    # model = load_model(model, key_size, value_size, num_heads, head)
  
    ## Predict track signal
    print("! Predict tracks", file=sys.stderr)
    results = []

    for (chrom, start, end, seq) in tqdm.tqdm(targets):
        seq = dna_1hot(seq)
        seq = np.array([seq[st:ed] for (st, ed) in it_stride])
        seq = tf.convert_to_tensor(seq, np.float32)

        predicted = model(seq, is_training=False)
        variance  = tf.math.reduce_variance(predicted, axis = 0)
        predicted = tf.math.reduce_mean(predicted, axis = 0)

        step = 128
        ssize = (end-start)
        psize = predicted.shape[0] * step
        if psize < ssize:
            offset = (ssize - psize) // 2
            pstart = start + offset
            pend = pstart + psize
        else:
            pstart = start
            pend = end
        # step = (end-start) // predicted.shape[0]
        pos = np.arange(pstart, pend, step, dtype=np.int32)
        res = [predicted[:, t] for t in tracks]
        var = [variance[:, t] for t in tracks]

        results.append((chrom, pos, step, res, var))
    
    ## Predict track signal
    print("! Generating bedgraph", file=sys.stderr)

    output = [open(f"{prefix}.{head}-{track}.bedGraph", "w") for track in tracks]

    for (chrom, start, step, signal, variance) in  tqdm.tqdm(results):
        for i in range(len(tracks)):
            result = [
                f"{chrom}\t{start[j]}\t{start[j] + step}\t{signal[i][j]:.5f}\t{variance[i][j]:.5f}"
                for j in range(start.shape[0])
            ]
            print("\n".join(result), file=output[i])
    return(0)


if __name__ == "__main__":
    load_dotenv()
    
    parser = argparse.ArgumentParser(prog="predict-bedgraph")
    parser.add_argument('bedfile', type=str)
    parser.add_argument('model', type=str)
    parser.add_argument('--reference', type=str)
    parser.add_argument('--length', type=int, default=os.getenv('SEQ_LENGTH', 25600))
    parser.add_argument('--extend', action='store_false')
    ## model parameters
    modeldef = parser.add_argument_group('model definition')
    modeldef.add_argument('--key-size', type=int, default=64)
    modeldef.add_argument('--value-size', type=int, default=64)
    modeldef.add_argument('--num-heads', type=int, default=1)
    ## output parameters
    outputdef = parser.add_argument_group('output definition')
    outputdef.add_argument('--head', type=str, default='mouse')
    outputdef.add_argument('--strides', type=int, default=10)
    outputdef.add_argument('--tracks', type=int, nargs="+", default=[10, 11])
    outputdef.add_argument('--prefix', type=str)

    args = parser.parse_args()
    print(args, file=sys.stderr)
    exit(main(**vars(args)))
```
