## Supplemental Code for "Iterative improvement of deep learning models using synthetic regulatory genomics": predict_payload.py

```
#! /usr/bin/env python3

import os
import sys
import argparse

import tqdm
import pysam
import numpy as np
import tensorflow as tf

from dotenv import load_dotenv
from src.utils import pad_or_crop, dna_1hot
from src.models import load_model


def parse_coordinate(coordinate:str) -> tuple:
    chrom, coord, *anno = coordinate.split(":")
    start, end = coord.split("-")
    return (chrom, int(start), int(end), anno)


def find(seq:str, sub:str) -> int:
    if sub in seq:
        return(seq.index(sub))
    return(-1)


def parse_payload(name, coord, length, context_sequence, reference):
    sequences = []
    labels = []
    for c in coord.split(","):
        chrom, start, end, anno = parse_coordinate(c)
        refs = [a for a in anno if a.startswith("f")]
        if len(refs) > 0:
            ref = pysam.FastaFile(refs[0][1:])
        else:
            ref = reference
        
        ## fill with Ns
        if any(a.startswith("d") for a in anno):
            sequences.append("N" * (end-start))
        else:
            sequences.append(ref.fetch(chrom, start, end))
        
        labs = [a for a in anno if a.startswith("n")]
        if len(labs) > 0:
            labels.append(labs[0][1:])
        else:
            labels.append(None)
    seqlen = [len(seq) for seq in sequences]
    payload_sequence = "".join(sequences)
    sequence = context_sequence[0] + payload_sequence + context_sequence[1]
    (cropstart, cropend) = pad_or_crop(0, len(sequence), length)
    sequence = sequence[cropstart:cropend]
    offset = len(context_sequence[0]) - cropstart

    labels = [
        (labels[i], find(sequence, sequences[i]), seqlen[i])
        for i in range(len(labels))
        if labels[i] is not None
    ]

    return(name, sequence, (offset, offset + len(payload_sequence)), labels)
    

def main(model:str, payload:str, 
        context:str, length:int, reference:str,
        head: str, strides:int, output:str, report:str,
        key_size:int, value_size:int, num_heads:int) -> int:
    ## Check input
    for file in [payload, reference]:
        if not os.path.exists(file):
            print(f"! Unable to locate `{file}`", file=sys.stderr)
            return(1)
        
    ## SETUP
    stride_it = [(i, i+length) for i in range(0, strides + 1)]
    reference = pysam.FastaFile(args.reference)
    report = sys.stdout if report == "-" else open(report, "w")

    ## Prepare track
    print("! Setup track", file=sys.stderr)
    total_length = length + strides

    if context is None:
        context_sequence = ("N" * total_length, "N" * total_length)
    else:
        chrom, start, end, _ = parse_coordinate(context)
        ctxleft = reference.fetch(chrom, start - total_length, start).upper()
        ctxright = reference.fetch(chrom, end, end + total_length).upper()
        context_sequence = (ctxleft, ctxright)

    payloads = [line.rstrip().split("\t")[:2] for line in open(payload, "r")]
    payloads = [parse_payload(*payload, total_length, context_sequence, reference)
                for payload in payloads]

    print("! Load model", file=sys.stderr)
    model = load_model(model, key_size, value_size, num_heads, head)
  
    ## Predict track signal
    print("! Predict tracks", file=sys.stderr)
    prediction = []
    shape = None
    for (name, sequence, offset, labels) in tqdm.tqdm(payloads):
        sequence = dna_1hot(sequence)
        sequence = np.array([sequence[st:ed] for st, ed in stride_it])
        sequence = tf.convert_to_tensor(sequence, np.float32)

        try:
            pd = model(sequence, is_training=False)
            pd = tf.math.reduce_mean(pd, axis=0)
            shape = pd.shape

            prediction.append(pd)
        except Exception as err:
            ## It will fail if none are present
            print(f"Unable to generate predictions at {chrom}:{start}-{end}", file=sys.stderr)
            print(err, file=sys.stderr)
            if shape is None:
                return(1)
            pd = tf.fill(shape, np.nan)
            prediction.append(pd)
        
        if len(labels) > 0:
            pd = np.repeat(pd, 128, 0)
            for (label, st, sz) in labels:
                if st == -1 or sz == 0:
                    continue ## Skip if sequence is not fully included
                val = np.max(pd[(st-strides):(st+sz+strides), :], 0)
                val = [f"{v:.3f}" for v in val]
                val = "\t".join(val)
                print(name, label, val, sep="\t", file=report)
    
    np.save(output, np.array([tensor.numpy() for tensor in prediction]))    
    return(0)


if __name__ == "__main__":
    load_dotenv()

    parser = argparse.ArgumentParser(prog="predict-array")
    parser.add_argument('model', type=str)
    parser.add_argument('payload', type=str)
    parser.add_argument('--context', type=str)
    parser.add_argument('--length', type=int, default=os.getenv('SEQ_LENGTH', 25600))
    parser.add_argument('--reference', type=str)
    ## model parameters
    modeldef = parser.add_argument_group('model definition')
    modeldef.add_argument('--key-size', type=int, default=64)
    modeldef.add_argument('--value-size', type=int, default=64)
    modeldef.add_argument('--num-heads', type=int, default=1)
    ## output parameters
    outputdef = parser.add_argument_group('output definition')
    outputdef.add_argument('--head', type=str, choices=['mouse', 'human'], default='mouse')
    outputdef.add_argument('--strides', type=int, default=10)
    outputdef.add_argument('--output', type=str, default="array.npy")
    outputdef.add_argument('--report', type=str, default="-")

    args = parser.parse_args()
    print(args, file=sys.stderr)
    exit(main(**vars(args)))
```
