## Supplemental Code for "Iterative improvement of deep learning models using synthetic regulatory genomics": train.py

```
#! /usr/bin/env python3

import os
import sys
import argparse

import tqdm
import numpy as np
import sonnet as snt
import tensorflow as tf

from dotenv import load_dotenv
from src.utils import read_dataset, generate_offsets, batch_iterator, dna_1hot
from src.models import load_original, build_finetune, restore_model
from src.models.enformer import Enformer


def load_dataset(prefix, folds):
    sequence = []
    activity = []
    for fold in folds:
        sequence.append(np.load(f"{prefix}-sequence.{fold}.npy"))
        activity.append(np.load(f"{prefix}-activity.{fold}.npy"))
    sequence = np.concatenate(sequence)
    activity = np.concatenate(activity)
    index = np.arange(sequence.shape[0])
    np.random.shuffle(index)

    dataset = tf.data.Dataset.from_tensor_slices((sequence[index, :, :], activity[index]))
    return(dataset)


def main(model: str, dataset: str, seed: int,
         kfold: int, num_kfolds: int, learning_rate: float,
         epochs: int, steps: int, batch: int,
         key_size: int, value_size: int, num_heads: int,
         baseline: str) -> int:
    ## Setup
    np.random.seed(seed)

    logfile = f"{model}-log.txt"
    logfile = open(logfile, "w")
    checkpointfile = f"{model}"

    ## Load dataset
    print("! Loading dataset", file=sys.stderr)
    if num_kfolds == 1:
        valid_dataset = load_dataset(args.dataset, ["fold0"])
        train_dataset = load_dataset(args.dataset, ["fold0"])
    elif num_kfolds > 0 and kfold >= 0 and kfold < num_kfolds:
        valid_dataset = load_dataset(args.dataset, [f"fold{kfold}"])
        train_dataset = load_dataset(
            args.dataset, [f"fold{i}" for i in range(num_kfolds) if i != kfold])
    else:
        train_dataset = load_dataset(dataset, ["train"])
        valid_dataset = load_dataset(dataset, ["validation"])
    train_dataset = iter(train_dataset.batch(batch).repeat())
    valid_dataset = iter(valid_dataset.batch(batch).repeat())

    ## Build model
    print("! Building model", file=sys.stderr)
    ## Define checkpoint and restore if available
    if baseline and os.path.exists(baseline):
        print(f"-> Baseline from {baseline}", file=sys.stderr)
        original = Enformer()
        checkpoint = tf.train.Checkpoint(module = original)
        checkpoint.restore(tf.train.latest_checkpoint(baseline)).expect_partial()
    else:
        print(f"-> Restored Enformer from from original", file=sys.stderr)
        original = load_original()
    finetune = build_finetune(original, key_size, value_size, num_heads)
    checkpoint = tf.train.Checkpoint(module = original, finetune = finetune)
    chkmanager = tf.train.CheckpointManager(checkpoint, checkpointfile, max_to_keep=5)
    if chkmanager.latest_checkpoint:
        checkpoint.restore(chkmanager.latest_checkpoint)
        print(f"-> Restore from {chkmanager.latest_checkpoint} @ {checkpointfile}", file=sys.stderr)
    else:
        print(f"-> Initializing from scratch @ {checkpointfile}.", file=sys.stderr)

    ## Setup training
    print("! Setup training", file=sys.stderr)
    learning_rate = tf.Variable(learning_rate, trainable=False, name="lr")
    optimizer = snt.optimizers.Adam(learning_rate = learning_rate)

    @tf.function
    def loss(inputs, output):
        pred = finetune(inputs, is_training=False)
        pred = tf.reshape(pred, (-1,))
        return tf.reduce_mean(tf.keras.losses.MSE(output, pred))
    
    @tf.function
    def train(inputs, output):
        with tf.GradientTape() as tape:
            pred = finetune(inputs, is_training=True)
            pred = tf.reshape(pred, (-1,))
            loss = tf.reduce_mean(tf.keras.losses.MSE(output, pred))
        gradient = tape.gradient(loss, finetune.trainable_variables)
        optimizer.apply(gradient, finetune.trainable_variables)
        return loss
    
    ## Run epochs
    train_loss = tf.keras.metrics.Mean(name="train_loss")
    valid_loss = tf.keras.metrics.Mean(name="validation_loss")
    print("! Running training", file=sys.stderr)
    for epoch in range(epochs):
        train_loss.reset_states()
        valid_loss.reset_states()
        progbar = tqdm.tqdm(range(steps))
        for _ in progbar:
            seq, activity = next(train_dataset)
            train_loss(train(seq, activity))
            seq, activity = next(valid_dataset)
            valid_loss(loss(seq, activity))
            progbar.set_description(f"Epoch #{epoch+1}\tTrain MSE={train_loss.result():.5f}")
        print(
            f"Epoch {epoch+1} / {args.epochs}, ",
            f"Train Loss: {train_loss.result():.5f}, ",
            f"Valid. Loss: {valid_loss.result():.5f}.", file=sys.stderr)
        print(epoch, float(train_loss.result()), float(valid_loss.result()), file=logfile)
        chkmanager.save()

    logfile.close()
    return(0)


if __name__ == "__main__":
    load_dotenv()

    parser = argparse.ArgumentParser(
        prog="src.models.train",
        description=(
            "Trains a finetuned Enformer model based on " +
            "{dataset}-sequence.{fold}.npy and {dataset}-activity.{fold}.npy " +
            "arrays."
        ))
    parser.add_argument('model', type=str, help="path to save or restore a finetuned model")
    parser.add_argument('dataset', type=str, help="training/validation dataset path prefix")
    parser.add_argument('--seed', type=int, default=5, help="random number generator seed")
    ## model parameters
    traindef = parser.add_argument_group('training options')
    traindef.add_argument('--baseline', type=str, help="Path to Enformer weights to use as baseline. Defaults to $ENFORMERBASELINE if not set")
    traindef.add_argument('--kfold', type=int, default=-1, help="fold to be used as validation in a kfold divided dataset.")
    traindef.add_argument('--num_kfolds', type=int, default=-1, help="total number of kfolds the dataset was divided into.")
    traindef.add_argument('--learning-rate', type=float, default=1E-5, help="finetuning training learning rate")
    traindef.add_argument('--epochs', type=int, default=10, help="number of training epochs to be conducted.")
    traindef.add_argument("--steps", type=int, default=100, help="number of trainins steps conducted per epoch.")
    traindef.add_argument("--batch", type=int, default=4, help="number of samples to be evaluated simultaneously per step. Note this has large implications to the memory requirements.")
    ## model parameters
    modeldef = parser.add_argument_group('model definition')
    modeldef.add_argument('--key-size', type=int, default=64, help='finetuning attention layer key size')
    modeldef.add_argument('--value-size', type=int, default=64, help='finetuning attention layer value size')
    modeldef.add_argument('--num-heads', type=int, default=1, help='number of independent finetuning attention heads')

    args = parser.parse_args()
    print(args)
    exit(main(**vars(args)))
```
