## Supplemental Code for "Iterative improvement of deep learning models using synthetic regulatory genomics": train_track_h5.py

```
#! /usr/bin/env python3

import os
import sys
import argparse

import tqdm
import h5py
import numpy as np
import sonnet as snt
import tensorflow as tf
import matplotlib.pyplot as plt

from dotenv import load_dotenv
from src.models import load_original
from src.models.enformer import Enformer, Sequential


class LazyH5Generator:
    def __init__(self, files:list):
        self._files = list(files)
    
    def __call__(self):
        for file in self._files:
            h5 = h5py.File(file, 'r')
            n = h5["input"].shape[0]
            for i in range(n):
                input = np.float32(h5["input"][i, ...])
                outcome = np.float32(h5["outcome"][i, ...])
                yield (input, outcome)
            h5.close()


def load_dataset(prefix:str, batches:int, validation_rate:float):
    dtype = (tf.float32, tf.float32)
    shape = None
    channels = None
    with h5py.File(f"{prefix}.0.h5", "r") as base:
        input_shape = base["input"].shape[1:]
        outcome_shape = base["outcome"].shape[1:]
        shape = (tf.TensorShape(input_shape), tf.TensorShape(outcome_shape))
        channels = outcome_shape[-1]

    files = [f"{prefix}.0.h5"]
    for i in range(1, batches):
        filename = f"{prefix}.{i}.h5"
        file = h5py.File(filename, "r")
        file_input_shape = file["input"].shape[1:]
        file_outcome_shape = file["outcome"].shape[1:]
        if file_input_shape == input_shape and file_outcome_shape == outcome_shape:
            files.append(filename)
    # files = [f"{prefix}.{i}.h5" for i in range(batches)]
    # split = np.max([1, int(batches * validation_rate)])
    split = np.max([1, int(len(files) * validation_rate)])
    valid_dataset = LazyH5Generator(files[:split])
    valid_dataset = tf.data.Dataset.from_generator(valid_dataset, output_types=dtype, output_shapes=shape)
    train_dataset = LazyH5Generator(files[split:])
    train_dataset = tf.data.Dataset.from_generator(train_dataset, output_types=dtype, output_shapes=shape)

    return((train_dataset, valid_dataset, channels))


def main(
        model:str, dataset:str, dataset_batches:int, seed:int, plot:bool,
        learning_rate:float, epochs:int, steps:int, batch:int,
        validation:float, baseline:str):
    if dataset_batches < 2:
        print("Number of batches must be 2 or above", file=sys.stderr)
        return(1)
    
    ## Setup
    np.random.seed(seed)

    logfile = open(f"{model}-log.txt", "a")
    checkpointfile = f"{model}"

    ## Load dataset
    print("! Loading dataset", file=sys.stderr)
    train_dataset, validation_dataset, channels = load_dataset(dataset, dataset_batches, validation)
    train_dataset = iter(train_dataset.batch(batch).repeat())
    validation_dataset = iter(validation_dataset.batch(batch).repeat())

    ## Prepare model
    print("! Building model", file=sys.stderr)
    if baseline and os.path.exists(baseline):
        print(f"-> Baseline from {baseline}", file=sys.stderr)
        original = Enformer()
        checkpoint = tf.train.Checkpoint(module = original)
        checkpoint.restore(tf.train.latest_checkpoint(baseline)).expect_partial()
    else:
        print(f"-> Restored Enformer from from original", file=sys.stderr)
        original = load_original()

    trunk = original._trunk
    trunk.trainable = False
    head = tf.keras.layers.Dense(channels, activation="softplus")
    model = Sequential([trunk, head])

    checkpoint = tf.train.Checkpoint(head = head)
    chkmanager = tf.train.CheckpointManager(checkpoint, checkpointfile, max_to_keep=5)
    if chkmanager.latest_checkpoint:
        checkpoint.restore(chkmanager.latest_checkpoint)
        print(f"-> Restore from {chkmanager.latest_checkpoint} @ {checkpointfile}", file=sys.stderr)
    else:
        print(f"-> Initializing from scratch @ {checkpointfile}.", file=sys.stderr)

    ## Setup training
    print("! Setup training", file=sys.stderr)
    learning_rate = tf.Variable(learning_rate, trainable=False, name="lr")
    optimizer = snt.optimizers.Adam(learning_rate = learning_rate)

    @tf.function
    def loss(inputs, output):
        predicted = model(inputs, is_training=False)
        error = tf.reduce_mean(tf.keras.losses.poisson(output, predicted))
        return error
    
    @tf.function
    def train(inputs, output):
        with tf.GradientTape() as tape:
            predicted = model(inputs, is_training=True)
            error = tf.reduce_mean(tf.keras.losses.poisson(output, predicted))
        gradient = tape.gradient(error, head.trainable_variables)
        optimizer.apply(gradient, head.trainable_variables)
        return error

    ## Run epochs
    print("! Running training", file=sys.stderr)
    train_loss = tf.keras.metrics.Mean(name="train_loss")
    valid_loss = tf.keras.metrics.Mean(name="validation_loss")
    
    for epoch in range(epochs):
        train_loss.reset_states()
        valid_loss.reset_states()
        progbar = tqdm.tqdm(range(steps))
        ct = 0
        for _ in progbar:
            seq, activity = next(train_dataset)
            train_loss(train(seq, activity))
            seq, activity = next(validation_dataset)
            valid_loss(loss(seq, activity))
            
            if ct % 20 == 0:
                progbar.set_description(f"Epoch #{epoch+1}\tTrain MSE={train_loss.result():.5f}")
            ct += 1
        print(
            f"Epoch {epoch+1} / {args.epochs}, ",
            f"Train Loss: {train_loss.result():.5f}, ",
            f"Valid. Loss: {valid_loss.result():.5f}.", file=sys.stderr)
        print(epoch, float(train_loss.result()), float(valid_loss.result()), file=logfile)
        chkmanager.save()

        if plot:
            inputs, outcome = next(validation_dataset)
            predicted = model(inputs, is_training=False)

            outcome = outcome.numpy()
            predicted = predicted.numpy()

            x = np.arange(predicted.shape[1])
            f, xs = plt.subplots(
                predicted.shape[2], predicted.shape[0],
                figsize=(4 * predicted.shape[0], 8), sharex=True)
            for i in range(predicted.shape[0]):
                for j in range(predicted.shape[2]):
                    xs[j, i].step(x, outcome[i, :, j], color="tab:blue")
                    xs[j, i].step(x, predicted[i, :, j], color="tab:red")
                    xs[j, i].set_ylim(0, 5)
            f.tight_layout()
            f.savefig(f"{checkpointfile}-{epoch}.png")
    logfile.close()
    return(0)


if __name__ == "__main__":
    load_dotenv()

    parser = argparse.ArgumentParser(
        prog="train-track",
        help = (
            "Trains a new output head based on a training dataset, while " +
            "preserving the original Enformer trunk weights. " +
            "During training, a portion of the training dataset are reserved " +
            "for validation and used only to measure each epoch performance. " +
            "Training is performed in epochs, where batches of examples are " +
            "presented in steps."
        )
    )
    parser.add_argument('model', type=str,
        help = "Path to model weights or either `original` or `tensorhub` which will load weights from ENFORMERBASELINE and ENFORMERTENSORHUB, respectively.")
    parser.add_argument('dataset', type=str,
        help = "Path prefix for a training dataset. It looks for {dataset}.*.h5 files.")
    parser.add_argument('--dataset-batches', type=int, default=1,
        help = "Number of training dataset batches to consider during training.")
    parser.add_argument('--seed', type=int, default=5,
        help = "Random number generator seed to ensure reproducibility.")
    parser.add_argument('--plot', action='store_true',
        help = "Flag to generate reporting plot [WIP].")
    ## model parameters
    traindef = parser.add_argument_group('training options')
    traindef.add_argument('--learning-rate', type=float, default=1E-4,
        help = "Training learning rate.")
    traindef.add_argument('--epochs', type=int, default=10,
        help = "Number of training epochs to conduct.")
    traindef.add_argument("--steps", type=int, default=100,
        help = "Number of training steps to be conducted per epoch.")
    traindef.add_argument("--batch", type=int, default=4,
        help = "Number of examples to evaluate simultaneously per training step.")
    traindef.add_argument("--validation", type=float, default=0.2,
        help = "Ratio of training datasets to be reserved for validation. Validation examples are only used to measure performance at the end of each epoch.")
    traindef.add_argument('--baseline', type=str,
        help = "Path to weights to use as baseline. Defaults to ENFORMERBASELINE if not set")

    args = parser.parse_args()
    print(args)
    exit(main(**vars(args)))
```
