## Supplemental Code for "Iterative improvement of deep learning models using synthetic regulatory genomics": utils.py

```
import numpy as np

from typing import Tuple, List, Iterable


def parse_annotation(annotation:str) -> Tuple[str, int, int]:
    name = annotation[:annotation.find(":")]
    start, end = annotation[annotation.find(":")+1:].split("-")
    return((name, int(start), int(end)))


def pad_or_crop(start:int, end:int, target_length:int) -> Tuple[int, int]:
    padding = target_length - (end - start)
    leftpad = padding // 2
    start -= leftpad
    end += padding - leftpad
    return (start, end)


def dna_1hot(
        seq:str, alphabet:str = "ACGT",
        neutral:str='N', neutral_value:float = 0.25,
        dtype:np.dtype = np.float32) -> np.ndarray:
    """
    based on kipoiseq
    """
    def to_uint8(string):
        return np.frombuffer(string.encode('ascii'), dtype=np.uint8)
    hasht = np.zeros((np.iinfo(np.uint8).max, len(alphabet)), dtype=dtype)
    hasht[to_uint8(alphabet)] = np.eye(len(alphabet), dtype=dtype)
    hasht[to_uint8(neutral)] = neutral_value
    hasht = hasht.astype(dtype)
    return hasht[to_uint8(seq.upper())]


def fold_split(ratio:List[float], size:int) -> np.ndarray:
    ratio_sum = sum(ratio)
    fold_size = np.array(
        [ np.ceil(size * r / ratio_sum) for r in ratio ], dtype=np.int32)
    fold = np.repeat(np.arange(len(ratio)), fold_size)
    np.random.shuffle(fold)
    return fold[:size]


def batch_iterator(it:Iterable, size:int) -> None:
    batch = []
    for i in it:
        batch.append(i)
        if len(batch) >= size:
            yield batch
            batch = []
    if len(batch) > 0:
        yield batch


def read_dataset(file:Iterable[str]) -> List[dict]:
    header = next(file).rstrip().split("\t")
    data = []
    for line in file:
        row = line.rstrip().split("\t")
        entry = dict(zip(header, row))
        ## Decode annotation
        if entry['annotation'] == "":
            entry["annotation"] = []
        else:
            entry["annotation"] = entry["annotation"].split(";")
            entry["annotation"] = [
                parse_annotation(annotation)
                for annotation in entry["annotation"]]
        data.append(entry)
    return(data)


def generate_offsets(
        total_length:int, target_length:int, stride:int,
        sample:float) -> np.ndarray:
    offsets = np.arange(0, total_length-target_length, stride)
    if sample < 1:
        sample_size = int(np.floor(offsets.shape[0] * sample))
    else:
        sample_size = int(min(sample, offsets.shape[0]))
    np.random.shuffle(offsets)
    offsets = offsets[:sample_size]
    offsets.sort()
    return offsets
```
